# Directed evolution of hyperactive integrases for site specific insertion of transgenes

**DOI:** 10.1101/2024.06.10.598370

**Authors:** Brian E. Hew, Sabranth Gupta, Ryuei Sato, David F. Waller, Ilko Stoytchev, James E. Short, Lisa Sharek, Christopher T. Tran, Ahmed H. Badran, Jesse B. Owens

**Affiliations:** Department of Cell and Molecular Biology, Institute for Biogenesis Research, John A. Burns School of Medicine, University of Hawaii at Manoa, Honolulu, Hawaii, 96814 USA; Department of Chemistry, Department of Integrative Structural and Computational Biology, Beckman Center for Chemical Sciences, The Scripps Research Institute, La Jolla, California, 92037 USA

## Abstract

The ability to deliver large transgenes to a single genomic sequence with high efficiency would accelerate biomedical interventions. Current methods suffer from low insertion efficiency and most rely on undesired double-strand DNA breaks. Serine integrases catalyze the insertion of large DNA cargos at attachment (att) sites. By targeting att sites to the genome using technologies such as prime editing, integrases can target safe loci while avoiding double-strand breaks. We developed a method of phage-assisted continuous evolution we call IntePACE, that we used to rapidly perform hundreds of rounds of mutagenesis to systematically improve activity of PhiC31 and Bxb1 serine integrases. Novel hyperactive mutants were generated by combining synergistic mutations resulting in integration of a multi-gene cargo at rates as high as 80% of target chromosomes. Hyperactive integrases inserted a 15.7 kb therapeutic DNA cargo containing Von Willebrand Factor. This technology could accelerate gene delivery therapeutics and our directed evolution strategy can easily be adapted to improve novel integrases from nature.

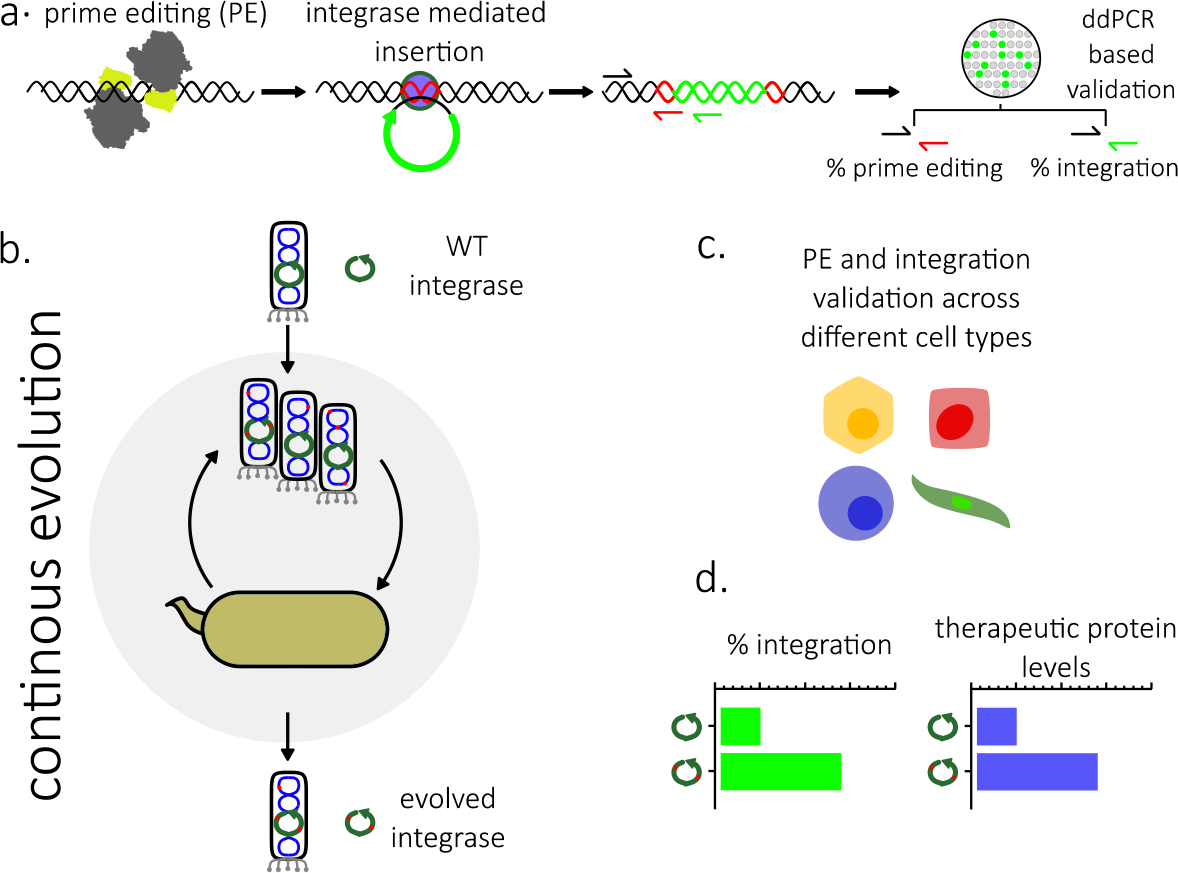

## INTRODUCTION

A longstanding challenge for the genome editing field has been the insertion of large DNA cargo at a specific target sequence at high efficiency and without unprotected double-strand breaks (DSBs) (1). Homology-directed repair (HDR) insertion efficiency drops significantly for larger cargos and HDR is inefficient or nonfunctional in non-dividing cells which make up most therapeutically relevant cell types (2,3). Nonhomologous end joining (NHEJ)-based insertion methods have been developed but, like HDR, rely on DSBs that can cause uncontrolled large deletions or chromosomal rearrangements (4,5).

By overcoming the lack of efficiency of current methods, numerous applications would be accelerated, including (i) integration into genomes of cell lines used in industrial biotechnology for the production of pharmaceuticals and other proteins (6); (ii) introduction of desirable traits, such as disease resistance or enhanced nutritional content, into plant genomes (7,8); (iii) insertion of disease-related genes into genomes of animal models to study the mechanisms of various disorders and test therapeutic interventions (9); (iv) integration of genetic modules into synthetic genomes, allowing the creation of biological memory devices or genetic logic gates (10,11); (v) attachment of epitope tags or fusion proteins to study gene regulation and localization *in situ* or *in vivo* (12); and (vi) delivering therapeutic genes into the genomes of patients’ cells (13–15). Tailoring a therapy to correct an individual’s unique mutation is challenging, hence a full replacement of the coding sequence of a recessive gene is often needed for a single, common therapy to be effective (16). As treatments for increasingly complex diseases are undertaken, multi-gene insertions will be necessary requiring larger cargos beyond the limits of current virus-based approaches (17).

Site-specific recombination involves the exchange of DNA strands between short, specific DNA sequences recognized by site-specific recombinases (SSRs) (18). SSRs cleave the DNA backbone at these sites, exchange the two DNA helices involved, and rejoin the DNA strands. These systems are employed naturally in a variety of cellular processes, including bacterial genome replication, differentiation and pathogenesis, and movement of mobile genetic elements (19).

Serine integrases are types of SSRs that are capable of actively inserting large DNA cargos at defined DNA sequences (20–22). During integrase recombination between attP (donor sequence) and attB (target sequence) attachment sites, DNA is protected during cleavage, strand exchange, and re-ligation (23,24). No synthesis or degradation of DNA occurs and no sequence homology is required for the donor DNA (21). This process does not rely on rate-limiting host factors and does not provoke error-prone DNA repair processes caused by nuclease cleavage that can induce a p53-mediated DNA damage response and lead to unwanted selection for cells with an impaired p53 pathway (21,25,26). A limitation of serine integrase technology is that the att site must first be inserted at the target sequence.

Fortunately, the small size of att sites has permitted their insertion using HDR into the genomes of mice and human cells to act as a landing pad for integrase insertion (27–30).

Prime editing (PE) is an established tool for precise targeting of short sequences to desired loci with advantages over HDR including high efficiency in both dividing and non-dividing cells as well as a reduction in DSBs (1,22). PE uses a Cas9 nickase that targets the DNA using a prime editing guide RNA (pegRNA) that encodes a reverse transcriptase template (RTT) which is “written” into the target site by a reverse transcriptase (RT) (Supplementary Fig. S1a). PE can be used for DNA substitutions, short insertions, and deletions (1). Several recent improvements to PE, including pegRNA engineering, optimized PE architecture, and twin prime editing have enabled the insertion of sequences >50 bp at high efficiencies (22,31–33).

PE has been used to insert att site landing pads for insertion of large DNA cargos by integrases (Fig. 1a and Supplementary Fig. S1a). Although high PE efficiency was achieved, the efficiency of insertion in mammalian cells by wildtype integrases has thus far been limited (1,34). We developed a directed evolution strategy using phage-assisted continuous evolution (PACE) (35) capable of rapidly evolving integrase activity, that we call IntePACE. We used IntePACE to perform over 200 rounds of mutagenesis in just a few days, a process that would take years using traditional directed evolution methods (35). By first delivering the att site using PE, we used evolved integrases to insert large DNA cargos at efficiencies several-fold higher than wildtype integrases into the human genome. We demonstrated activity in diverse cultured cell lines and primary human fibroblasts. A large 15.7 kb therapeutic DNA cargo encoding von

**Fig. 1.**
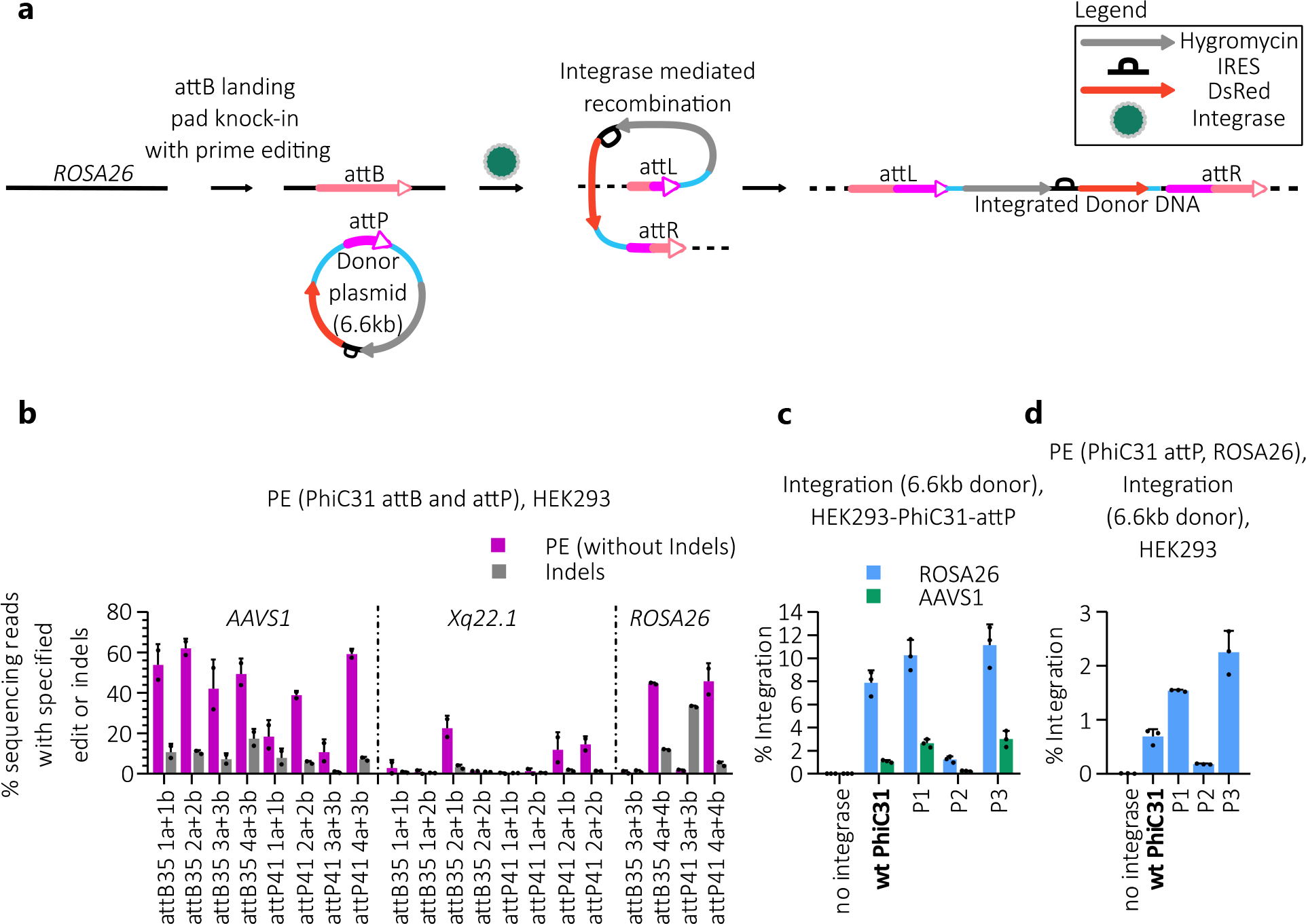
**PE optimization and PhiC31 integrase targeted insertion into the human genome. a**, Schematic of PE directed integrase gene insertion. PE efficiently inserts sequences <100bp. PE is first used to insert the 35bp attB site at a desired genomic locus, such as ROSA26. The attB site acts as a landing pad for PhiC31 integrase which can efficiently insert multi-kb sequences. During a second step, the integrase recombines the attB site found in the genome with an attP site found on the 6.6 kb donor plasmid. Following recombination, the entire donor plasmid becomes inserted into the genome and the attB and attP sites are rearranged to form attL and attR sites flanking the insertion. Because the integrase cannot recombine the newly formed attL and attR sites, the donor plasmid containing the Hygromycin antibiotic resistance gene and DsRed florescent protein gene is permanently inserted into the genome. **b**, Amplicon sequencing was used to compare efficiencies of PE for insertion of PhiC31 integrase attB (35 bp) or attP (41 bp) sites at AAVS1, Xq22.1, and ROSA26 in HEK293 cells. For each target sequence, twin epegRNA were used (listed on the x-axis). Amplicon sequencing primers were designed to flank each target sequence. PE efficiencies reflect sequencing reads containing the intended insertion of the att site. Indel efficiencies reflect reads with unintended insertions or deletions. N = 2. **c**, Integration efficiency of PhiC31 integrase mutants from Keravala et al. (39) at pre-inserted attP sites (AAVS1 and ROSA26) in clonal HEK293 cell lines. Cells previously modified to contain a PhiC31 attP site were co-transfected with a 6.6 kb donor plasmid containing an attB site, Hygromycin gene, and DsRed gene along with a helper plasmid containing one of four integrase variants. Integration of the donor plasmid was measured by ddPCR using a forward primer and probe designed to bind the genome and reverse primer designed to bind the inserted sequence. n = 3. **d**, PE plasmids, including attP41 4a+4b pegRNA from (b) designed to insert the PhiC31 attP site, were co-transfected with a donor plasmid containing an attB site and a helper plasmid encoding PhiC31 integrase variants in HEK293 cells. Integration efficiency at ROSA26 was measured by ddPCR. n = 3. Data are shown as mean + s.d. All ddPCR measurements were normalized using the product/reference ratio to account for differences in the number of target sequences in the genome (Supplementary Note S1 and Supplementary Table S1).

Willebrand Factor (vWF) was assessed for genome targeting and protein expression. We modelled evolution of the previously characterized PhiC31 and Bxb1 integrases. However, any integrase with a known att sequence has the potential for significant enhancement using this system.

## MATERIAL AND METHODS

### Plasmid design

#### General methods

Oligonucleotides and synthetic DNA fragments were purchased from Integrated DNA Technologies. PCR amplification of DNA fragments was performed with KOD One PCR Mastermix (Toyobo). Plasmids were constructed using NEBuilder HiFi DNA Assembly or USER cloning (New England Biolabs) and purified with the Zyppy Plasmid Miniprep Kit (Zymo).

DNA for transfection was purified by Zymopure Plasmid Miniprep Kit (Zymo), Zymopure Express Plasmid Midiprep Kit (Zymo), or Qiagen Plasmid Plus Midiprep Kit (Qiagen). Plasmids used in these experiments are listed in Supplementary Table S10.

#### Engineered pegRNA (epegRNA) design

Spacer sequences within selected human genomic target sites (AAVS1, Xq22.1, and ROSA26) were selected using CRISPOR (36). The protective tevopreQ1 motif (22,32) was added to the optimized cr772 scaffold (33) with a linker designed using pegLIT (32). Sequences for epegRNA designed for this study are listed in Supplementary Table S11. Designed epegRNA coding sequences were ordered as gBlocks (Integrated DNA Technologies) and subcloned into the pU6 pegRNA GG acceptor plasmid (Addgene #132777) (1).

#### Mammalian expression plasmids

Helper plasmids for mammalian expression called pCMV2 PhiC31 wt, pCMV2 HuOpt Bxb1, and pCMV2 HuOpt Pa01 encode wildtype PhiC31 (37), Bxb1 (22), and Pa01 (38) integrases, respectively. Hyperactive mutations in PhiC31 and Bxb1 integrases identified using directed evolution are listed in Supplementary Table S7 and S8, respectively. These mutations were incorporated into pCMV2 PhiC31 wt, and pCMV2 HuOpt Bxb1 for use in genome targeting experiments with mammalian cells. Hyperactive mutations originally identified by Keravala et al. (39) were also incorporated into pCMV2 PhiC31 wt to generate plasmids called pCMV2 PhiC31 P1, pCMV2 PhiC31 P2, and pCMV2 PhiC31 P3. Donor plasmids contain an att site and cargo DNA for genomic insertion. Donor plasmids pCMV PhiC31 attB DsRed Hygro, pCMV Bxb1 attP DsRed Hygro, and pCMV Pa01 attP DsRed Hygro contain att sites for PhiC31, Bxb1, and Pa01 integrases, respectively. All att sites are listed in Supplementary Table S4. The donor plasmids each contain the DsRed fluorescent protein and hygromycin resistance genes. In addition to DsRed and hygromycin, the donor plasmid pCDNA3.1-WT-VWF Bxb1 attP HIR Donor contains the therapeutic vWF gene. The PE5max plasmid (Addgene #174828) encodes an *S. pyogenes* Cas9 (SpCas9) nickase fused to a reverse transcriptase used for PE (31). Plasmids used for expressing the epegRNA include a spacer sequence that directs the SpCas9 nickase to the genomic target sequence (22,32). The epegRNA expression plasmids include a reverse transcription template (RTT) for the att site that becomes incorporated at the target sequence by PE. The epegRNA expression plasmids were used for PE insertion of the att site at human genomic target sequences AAVS1, Xq22.1, and ROSA26.

Helper, donor and epegRNA plasmids will be made available through Addgene.

#### PACE plasmids

Accessory plasmids (AP) for PACE called AP LinkRec PhiC31 attP and AP LinkRec Bxb1 attP48 contain the PhiC31 attP site (60 bp) or Bxb1 attP site (48 bp) adjacent to the 1,209 bp encoding the C-terminal 403 amino acids of M13 bacteriophage gene III (gIII).

Complementary plasmids (CP) called CP LinkDon PhiC31 attB or CP LinkDon Bxb1 attB contain a swappable promoter (Supplementary Fig. S8a and Supplementary Table S9) upstream of the 66 bp encoding the N-terminal 22 amino acids of gIII immediately adjacent to the PhiC31 attB site (70 bp) or Bxb1 attB site (50 bp). Att sites are listed in Supplementary Table S4. Variants of APs and CPs were constructed with different origins of replication to vary plasmid copy number as listed in Supplementary Table S10. Selection phage (SP) plasmids SP RecA PhiC31 and SP RecA Bxb1 contain the M13 phage genome in which gIII has been removed and replaced with PhiC31 and Bxb1 integrase genes respectively, and the gene VI promoter was replaced with the RecA promoter (Supplementary Note S4 and Supplementary Fig. S8b). Hyperactive variants of PhiC31 integrase described by Keravala et al. (39) were incorporated into SP RecA PhiC31 P1, SP RecA PhiC31 P2, and SP RecA PhiC31 P3. The mutagenesis plasmid (MP6 – Addgene #69669) provides increased mutagenesis rates by expressing proteins involved in mismatch repair (ugi and SeqA),

DNA base excision (dam and cda1), DNA base selection (emrR), and proofreading (dnaQ926) under the control of the arabinose inducible pBAD system (40).

### Mammalian cell culture

#### General culture conditions

HEK293 (ATCC CRL-1573) and HEK293T (Thermo Fisher Scientific NC0260915) cells were maintained in DMEM + GlutaMAX (Gibco), HeLa (Sigma 93021013) cells were maintained in EMEM with L-glutamine (ATCC), K562 (Sigma 89121407d) human lymphoblast cells were maintained in RPMI-1640 media (Sigma), and U2OS (ATCC 30-2007) human osteosarcoma cells were maintained in McCoy’s 5A media (Gibco). Media for HEK293, HEK293T, HeLa, K562, and U2OS cells was supplemented with 10% heat inactivated FBS (Neuromics), 1x Penicillin/Streptomycin (Gibco), and 0.75 µg/ml Amphotericin B (Gibco). PBS, trypsin, and Opti-MEM were purchased from Gibco. Adult human dermal fibroblast (HDFa) cells (Cascade Biologics C-013-5C) were maintained in Human Fibroblast Expansion Basal Medium (Gibco) supplemented with Low Serum Growth Supplement (Gibco).

#### Transfection reagents

HEK293, HEK293T, and K562 cells were transfected with TransIT-2020 (Mirus Bio). HeLa cells were transfected with HeLa-Monster (Mirus Bio). HDFa cells were transfected with Lipofectamine 3000 (Invitrogen). U2OS cells were electroporated using Neon Transfection system (Invitrogen MPK10025).

#### Clonal HEK293 cell lines

PE was used to generate single cell clones containing an att site at AAVS1 or ROSA26. HEK293 cells were transfected with 450 ng PE5max and 150 ng each of twin epegRNA plasmids (a and b) directed to either AAVS1 or ROSA26 in 24-well plates. Cells were serially diluted and visually confirmed to have a single colony per well. The presence of the att site was confirmed by ddPCR using a forward primer and probe found in the genome and a reverse primer found on the att site. For transfections with clonal lines containing a pre-inserted att site, cells (1 x 10^5^) were seeded in 24-well plates and transfected the next day with 360 ng of helper plasmid and 240 ng of donor plasmid. After three days, cells were lysed for analysis by ddPCR to determine the integration efficiency of the donor plasmid. Primer sequences are listed in Supplementary Table S6 and sequences of epegRNA expression plasmids are listed in Supplementary Table S11.

#### Simultaneous PE and integration assays in HEK293, HEK293T, HeLa, K562, U2OS, and HDFa cell lines

HEK293, HEK293T, and HeLa cells (1 x 10^5^) were seeded in 24-well plates. The next day, wells were transfected with 300 ng of PE5max, 30 ng of each epegRNA plasmid, 120 ng of helper integrase expression plasmid, and 120 ng of donor plasmid. K562 cells (4 x 10^6^) were seeded in 6-well plates. U2OS cells (1.4 x 10^6^) were electroporated (described above) with 2.5 µg of PE5max, 250 ng of each epegRNA plasmid, 1 µg of integrase expression plasmid, and 1 µg of donor plasmid. After electroporation the cells were transferred to a 6-well plate. HDFa cells (4.7 x 10^5^) were seeded in 6-well plates and the next day cells were transfected with 3.6 µg of PE5max, 360 ng of each epegRNA plasmid, 1.44 µg of integrase expression plasmid, and 1.44 µg of donor plasmid DNA using Lipofectamine 3000. Four hours later, media was replaced to remove excess reagent-DNA complexes. Conditions for the Neon transfections of U2OS cells were adapted from the ThermoFisher Scientific protocol. Briefly, 5 µg of DNA was added to 1.4 x10^6^ cells with electroporation parameters set to 1230V, 10 ms and 4 pulses. After electroporation, cells were transferred to 6-well plates. After three days, cells were lysed for analysis. All donor plasmids expressed DsRed. To reduce the influence of transfection efficiency, HEK293T, K562, HeLa, U2OS, and HDFa cells in experiments shown in Fig. 6 were sorted for DsRed to ensure uptake of the DNA.

#### Cell lysis

Cell pellets were lysed by addition of 90 µl DirectPCR Cell lysis reagent (Viagen Biotech) with 1.0 mg/mL freshly prepared proteinase K (Meridian Bioscience) and rotated in a hybridization oven for three hours at 55°C, then heat-inactivated for 45 min at 85°C and stored at -20°C prior to assaying.

#### FACS

All donor plasmids constitutively expressed DsRed. For experiments requiring sorting of transfected cells and exclusion of non-transfected cells, including select HEK293T transfections (Fig. 6a and 6g) as well as Hela, K562, U2OS, and HDFa transfections (Fig. 6b-e) a FACSAria Iiu flow cytometer (BD Biosciences) was used to sort cells based on DsRed fluorescence intensity (excitation 488 nm, emission 610 nm). Following FACS, cells were immediately pelleted and lysed for ddPCR analysis, described below.

### Amplicon sequencing

#### Sample preparation for amplicon sequencing

HEK293 cells were transfected with 600 ng of PE5max and 60 ng of each epegRNA plasmid (a and b). After three days, cells were lysed as described above. To determine editing outcomes using amplicon sequencing, a first round of PCR (PCR1) was performed using KOD One polymerase (Toyobo), 0.3 µM of locus specific barcoded primers containing a Nextera P5- and P7-tag and 1µL of cell lysate in a total volume of 20 µL. An 8 bp barcode sequence was inserted between the Nextera tag sequence and the annealing sequence of the primers. The barcode sequences are shown preceding the lowercase annealing sequences in Supplementary Table S6. PCR cycle conditions were 98°C for 2 min followed by 30 cycles of 98°C for 10 sec, 60°C for 5 sec, and 68°C for 5 sec with a final extension at 68°C for 30 sec. Products from PCR1 were run on a 2% agarose gel in TAE (Tris-acetate-EDTA) buffer and fragments were extracted using the Zymo Gel DNA recovery Kit (Zymo). Eluted products were pooled in equimolar amounts and were submitted to the UC Davis DNA Technologies and Expression Analysis Core lab for a second round of PCR (PCR2) and amplicon sequencing.

#### Analysis of amplicon sequencing data

Amplicon sequencing results were first demultiplexed using ultraplex (41) based on barcodes allowing one barcode mismatch. The barcode sequences were trimmed, and low-quality reads were discarded using a q-score of 20 (default is 30). To sort the forward and reverse reads in the same order, the sequences were re-paired with the repair function in Bbtools. Editing and indel frequencies were determined as previously described (42). Briefly, CRISPResso2 was run in HDR mode, discard_indel_reads flag was on and quantification window coordinates were set to 10 bp upstream of each nick site. Unedited template and PE sequences were input. Data from the “CRISPResso_quantification_of_editing_frequency” file was used to determine editing frequency by dividing “Read_aligned” (in the HDR tab) by “Reads_aligned_all_amplicons”. The indel frequency was determined by dividing total discarded reads (Reference + HDR) by “Reads_aligned_all_amplicons”. Examples of PE sequences as well as primers can be found in Supplementary Table S5 and S6.

### ddPCR

ddPCR was performed according to manufacturer’s instructions (BioRad Laboratories). Briefly, each reaction (22 µl) contained 0.5 µl of cell lysate, 11 µl of 2X ddPCR Supermix for Probes (no dUTP), 10 units XhoI (New England Biolabs), primers for target and reference at 900 nM and probes at 250 nM. The plate was sealed with PX1 PCR Plate Sealer (BioRad), vortexed thoroughly to mix, and centrifuged one min at 1000 rpm prior to droplet generation using the QX200 AutoDG Droplet Generator System (BioRad). PCR was performed as follows: denaturation at 95°C for 10 min, 45 cycles of 95°C for 30 sec, 60°C for two min and 72°C for two min, followed by a final incubation at 98°C for 10 min. Droplets were read on a QX200 Droplet Reader (BioRad) and analyzed using QX Manager 1.2 Standard Edition software (BioRad). Primers are listed in Supplementary Table S6.

### IntePACE

#### IntePACE strategy and selection stringency

During IntePACE, the phage gIII that is necessary for phage proliferation is removed from the selection phage (SP) genome and placed on plasmids in host cell *E. coli*. In place of gIII, the SP contains the coding sequence for the integrase to be evolved. The intended activity of the integrase is the recombination of the integrase attB site with the attP site. This activity is linked to the generation of gIII such that successful integrases capable of acting on their att sites enable propagation of infectious phage harboring their coding sequence. Host cells contain a mutation plasmid (MP), described below, and gIII expression plasmids called the accessory plasmid (AP) and complementary plasmid (CP) to enable selection for integrases with improved activity. Leaky gIII expression could lead to propagation of phage without intended activity. To prevent leaky gIII expression, the gene was split. The AP contains the full-size wildtype attP site (Supplementary Table S4) upstream of the C-terminus (N1-C domains) of split gIII. The CP contains a promoter upstream of the N-terminus of gIII (the leader sequence) and the full-size wildtype attB site (Supplementary Fig. S8c, Supplementary Note S4, and Supplementary Table S4). Successful crossover of the att sites on the AP and CP by an active integrase variant forms a single plasmid with a complete gIII (Fig. 2). The resulting attL site is in frame and translated, residing in a linker region between the leader sequence and N1 domain to prevent disruption of gIII activity (Supplementary Fig. S8d) (43).

**Fig. 2.**
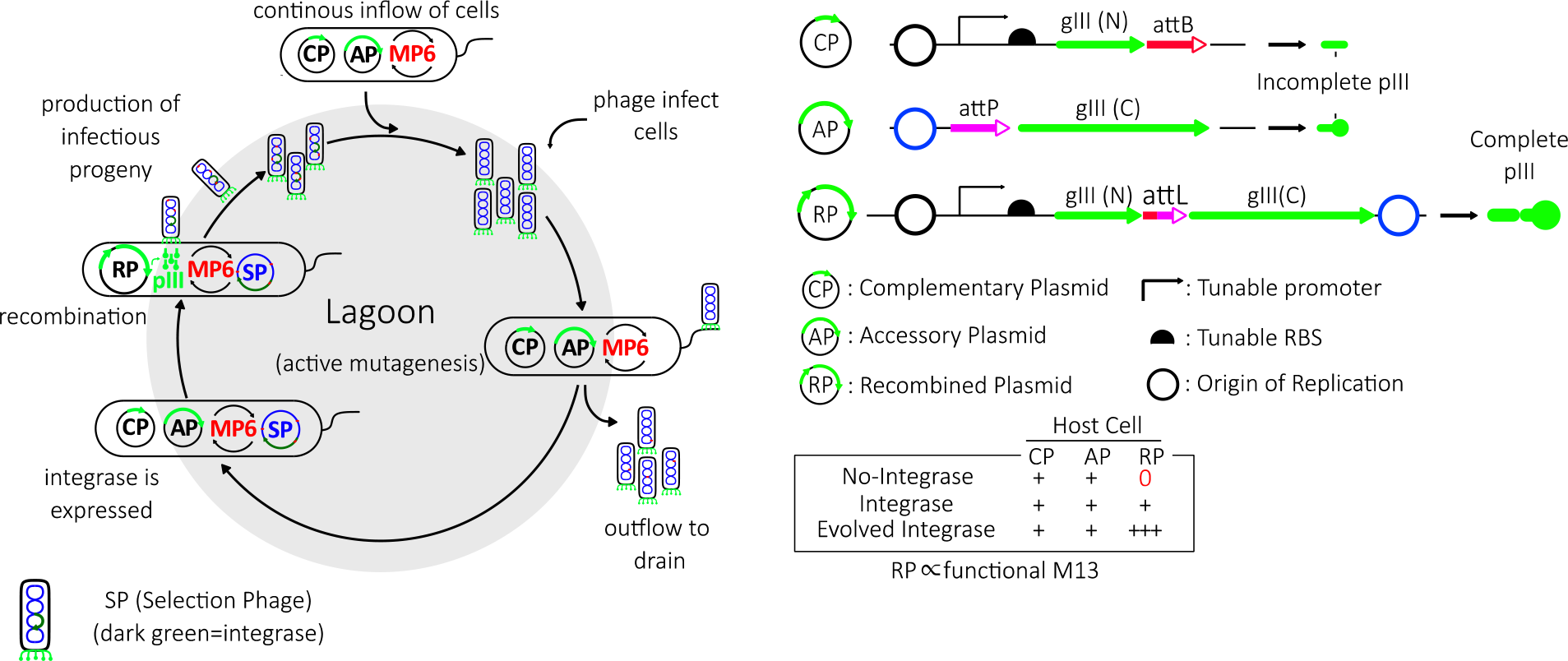
**IntePACE.**, Schematic of IntePACE. The integrase that is to be evolved is encoded in the phage genome. The phage gIII, which is essential for phage infectivity, is removed from the genome. PIII, encoded by gIII, is naturally separated into N-terminal and C-terminal domains by a flexible linker. The accessory plasmid (AP) contains the attP site and C-terminal portion of gIII. The complementary plasmid (CP) contains the attB site and a promoter driving the N-terminal portion of gIII called the leader sequence. Expression can be tuned by replacing a promoter or ribosome binding site (RBS) of desired strength. Both AP and CP are maintained in the host *E. coli* cells, and because gIII is split, no functional pIII is expressed. Key to IntePACE is the link between gIII expression and the desired activity of the integrase, specifically, the recombination of the attB site on the CP with the attP site on the AP. Upon crossover between the att sites the AP and CP join to form a single recombined plasmid (RP) and the promoter, leader sequence, and C-terminal portion of gIII join to permit the expression of pIII. During IntePACE, host cells harboring both the AP and CP are continuously added to a fixed volume vessel called the ‘lagoon’. The cells are then infected by the selection phage (SP) leading to expression of the integrase. Highly active integrase variants are expected to recombine more plasmids than less active variants. This results in higher levels of pIII and higher numbers of infectious progeny from the phage that encode the integrase variant capable of recombining the att sites. The lagoon is continuously drained at the same speed as the host cells are added. Integrase variants with poor efficiency of recombination do not produce enough infectious progeny and are diluted from the system. Stringency can be modulated by controlling the speed of the outflow of the host cells. Stringency can also be increased by reducing the strength of the promoter on the CP to limit the amount of gIII expression. During stringent conditions, only highly active integrases can recombine enough AP and CP at low pIII levels or recombine quickly enough to produce progeny before the lagoon drains the cells. New mutations are incorporated into progeny from phage encoding successful integrase variants due to the presence of the mutation plasmid (MP6) (40) in the host cells. Therefore, each cycle of phage replication generates a new mutation library from the fittest variants of the previous library.

During IntePACE, host cells are continually added at a specified rate to a fixed volume flask (the lagoon) and drained at an equal rate. Only phage capable of recombining AP and CP express gIII and produce phage fast enough to keep from being drained from the system. By increasing the flow rate of the lagoon, the evolving integrase variants are challenged by higher stringency. By decreasing the promoter strength of gIII, more recombination events are required to generate enough gIII expression to support phage proliferation. Lowering the copy number of the AP and CP reduces the number of available plasmids for recombination and provides a third method for increasing stringency. Plasmid copy number was optimized using AP and CP that each contained a different origin of replication. Early, less-stringent experiments used higher copy plasmids pUC and RSF1030-A8 (Supplementary Fig. S8e). Five promoters were used to tune gIII expression levels (Supplementary Fig. S8a). Stringency was periodically increased during each PACE experiment by decreasing promoter strength, decreasing plasmid copy number, or increasing flow rate (Fig. 3a and 4b, and Supplementary Fig. S4a and 6a). A complete list of AP and CP plasmids used for each PACE experiment is found in Supplementary Table S2.

**Fig. 3.**
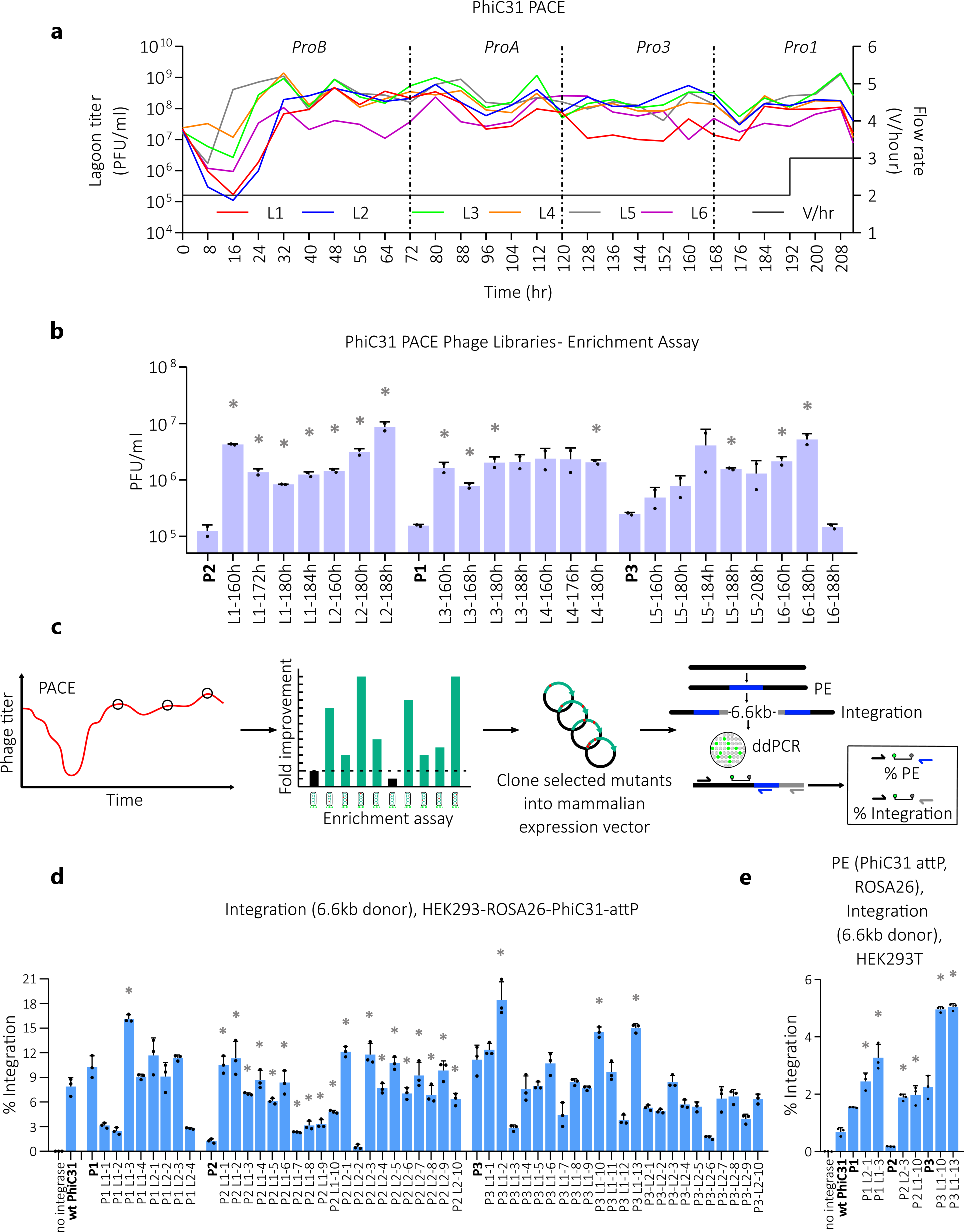
**IntePACE evolved hyperactive PhiC31 integrase. a**, IntePACE lagoon phage titers. Six lagoons were inoculated with phage containing SP encoding PhiC31 P1, P2, and P3 integrase (two lagoons each). Host cells containing the MP, AP, and CP with one of four gIII promotors (in order of strongest to weakest) ProB, ProA, Pro3, and Pro1 (53) were exchanged to increase stringency at 0, 76, 124, and 176 hour time points (black dotted lines), respectively. The flow rate was increased from 2 to 3 lagoon volumes per hour (black solid line) at the 188 hour time point. Phage were collected at least every 4 hours. Titers counted by activity-independent plaque assay are shown on the y-axis in log scale. **b**, Activity in *E. coli* of PhiC31 integrase evolved libraries from IntePACE time points assayed by overnight enrichment assay. Cells containing AP and CP were infected by phage encoding integrase variants isolated from IntePACE. Active integrase variants recombine the AP and CP resulting in pIII expression and increased phage progeny. Recombination activity was measured by counting plaques generated by the phage. P2 (lagoons L1 and L2), P1 (lagoons L3 and L4), and P3 (lagoons L5 and L6) (shown in bold font) represent starting point phage at the 0-hour time point (x-axis). Time point labels begin with lagoon number (L1-L6) followed by the hour the phage was collected (x-axis) n = 2. **c**, Schematic for strategy for assaying IntePACE derived clones in human cells. During IntePACE, integrase variants are challenged by stepwise increases in stringency as phage concentration is measured at least every 4 hours. Promising mutation libraries are chosen from time points with high phage titer (black circles). Next, the activity of the different libraries is compared in *E. coli* using the overnight enrichment assay. Integrase variants from top libraries are cloned into mammalian expression plasmids. PE and integrase plasmids are transfected into HEK293 cells. Following insertion of the att site by PE, integrase variants insert the donor plasmid by recombining the att site found on the donor plasmid with the att site now found in the genome. The ddPCR probe and forward primer are located in the genome. Both PE and insertion efficiency are measured using ddPCR using reverse primers found on the att site or donor plasmid, respectively. **d**, Helper plasmids encoding evolved PhiC31 integrase mutants were co-transfected with a 6.6 kb donor plasmid into a clonal HEK293 cell line containing a pre-installed attP site at ROSA26. Two lagoons were used for each starting variant. Helper plasmid clone names containing starting variant P1, P2, or P3, and the first or second lagoon number L1 or L2 for each, and clone number are listed on the x-axis. All mutations are listed in Supplementary Table S7. n = 3. **e**, Integration of 6.6 kb donor plasmid by select evolved PhiC31 integrase mutants with PE insertion of the attP site into unmodified HEK293 cells at ROSA26 (data for all mutants shown in Supplementary Fig. S2D). n = 3. Data are shown as mean + s.d. (*) = P value < 0.05 derived from a Student’s two- tailed t-test if significantly higher than the pre-evolved integrase control (shown in bold on the left).

**Fig. 4.**
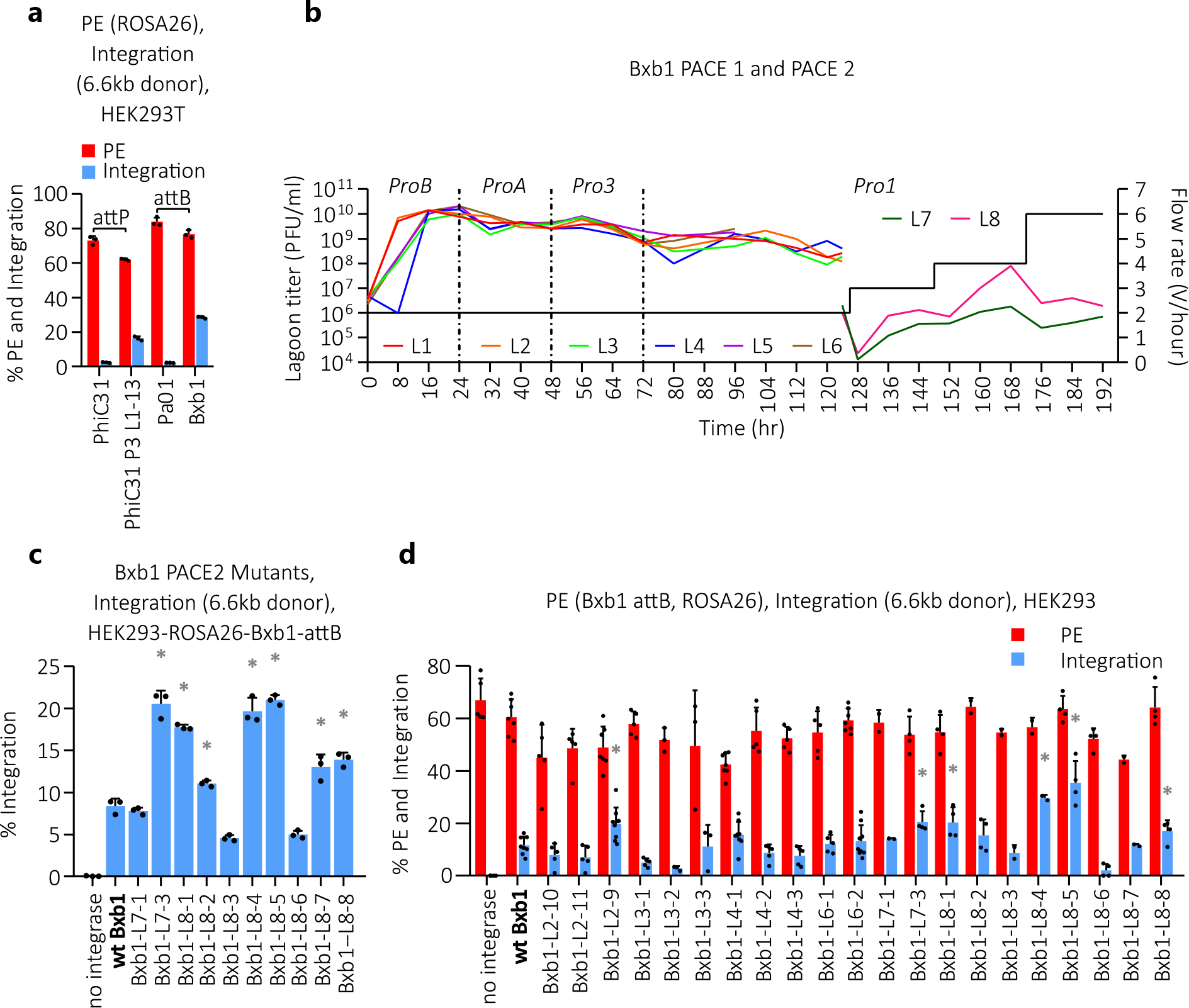
**IntePACE evolved hyperactive Bxb1 integrase. a**, Comparison of both PE efficiency of insertion of the att site and integration efficiency of a 6.6 kb donor plasmid of wildtype PhiC31, evolved PhiC31, Pa01, and Bxb1 integrases in HEK293T cells measured by ddPCR. n = 3. **b**, Bxb1 IntePACE lagoon phage titer (colored lines) measured by activity-independent plaque assay shown on the y-axis in log scale. Similar to PhiC31 IntePACE (described in detail in Fig. 3A), stringency was increased throughout the experiment. The gIII promoter strength was decreased by switching to host cells containing sequentially weaker promoters: ProB, ProA, Pro3 and Pro1. Host cells were changed every two days in six lagoons during the first experiment (black dotted lines) which lasted 5 days. In PACE 2, two lagoons were seeded with selected phage from PACE 1 (L7 was seeded with PACE 1 L4-120hr and L8 was seeded with PACE 1 L6-100hr). Flow rates were increased incrementally each day (black solid line). **c**, Bxb1 integrase mutants isolated from IntePACE were cloned into mammalian expression plasmids and used to measure insertion efficiency of a 6.6 kb donor plasmid (process described in Fig. 3C). Integration efficiency of evolved Bxb1 integrase mutants in a clonal HEK293 cell line with a preinstalled ROSA26 attB site was measured by ddPCR. n = 3. **d,** PE efficiency of insertion of the att site and integration efficiency of a donor plasmid by evolved Bxb1 integrases in unmodified HEK293 cells was measured by ddPCR. n ≥ 4. Data are shown as mean + s.d. (*) = P value < 0.05 derived from a Student’s two-tailed t-test if significantly higher than the pre-evolved integrase control (shown in bold on the left).

#### IntePACE conditions

*E. coli* S2060 (F’ proA+B+ Δ(lacIZY) zzf::Tn10 lacIQ1 PN25-tetR luxCDE Ppsp(AR2) lacZ luxR Plux groESL / endA1 recA1 galE15 galK16 nupG rpsL ΔlacIZYA araD139 Δ(ara,leu)7697 mcrA Δ(mrr-hsdRMS-mcrBC) proBA::pir116 araE201 ΔrpoZ Δflu ΔcsgABCDEFG ΔpgaC λ–) (44) cells were transformed with AP, CP, and MP6 plasmids and cultured in Davis Rich Media (DRM) (US Biological) supplemented with trace metals 3 nM ammonium molybdate (Ward’s Science), 400 nM Boric Acid (Beantown Chemical), 30 nM Cobalt(II) Chloride (Millipore), 10 nM Copper(II) Sulphate (STREM), 80 nM Manganese(II) Chloride (Acros), 10 nM Zinc Sulfate (Beantown Chemical), 500 nM Calcium Chloride (Ward’s Science), 0.1% tween-80 (Sigma), and antibiotics to maintain plasmids (AP – 50 µg/ml Carbenicillin, CP – 50 µg/ml Kanamycin, MP – 25 µg/ml Chloramphenicol) (Gold Bio) in addition to 10 µg/ml Tetracycline (Gold Bio), 10 µg/ml Flucconazole (Caymus), and 10 µg/ml Amphotericin B (Gold Bio). Host cells were cultured in chemostats to maintain a supply of fresh uninfected cells for the selection scheme. Over time, mutagenesis of the MP can inactivate the plasmid rendering the cells insensitive to arabinose and decrease the mutation during SP replication. To prevent this, host cell colonies were assayed for sensitivity to arabinose before expansion to chemostats. Cells plated on arabinose form smaller, slower growing colonies than when plated on glucose due to the mutagenesis activity of a functional (arabinose sensitive) MP (45). Each chemostat was maintained at 200-250 ml at a OD600 of 0.3-0.7 (flow rate ∼3-5 ml/min) and replaced with fresh arabinose sensitive host cells every two days. Two to six lagoons per experiment were maintained at 40 ml (except at high flow rates where volumes were decreased) with 100mM L-arabinose (Gold Bio) and flow rates ranged from 1-10 volumes/hour. A schematic of the PACE apparatus is found in Supplementary Fig. S8f.

#### Phage preparation and mutagenesis

The SP backbone was modified by replacing the gene VI promoter with the RecA promoter to decrease generation of recombinant wildtype M13 phage (Supplementary Fig. S8b). To begin IntePACE, host cells containing AP, CP, and MP6 were added to the lagoons and infected with 10^8^ plaque forming units (PFU) of starter phage. Active phage capable of recombining the AP and CP express gIII and generate progeny containing a functional integrase variant. Further mutagenesis of the integrase coding sequence is achieved by expression from the MP6 plasmid resulting in decreased proofreading and increased error-prone bypass during phage replication (40). Upon host cell entry to the lagoon, mutagenesis via the MP6 plasmid was induced at concentrations of 100 mM arabinose. Lagoons were sampled every 1-4 hours throughout each PACE experiment.

#### Overnight phage enrichment assay

*E. coli* cultures containing both AP and CP (2 ml, OD600 = 0.3-0.6) were inoculated with 10^5^ PFU/ml of purified M13 phage and incubated at 37°C at 180 rpm for 18 hours. Cultures were centrifuged at 12,000 x g for two minutes and supernatants were collected. Phage counts were obtained by activity-independent plaque assay (46). Briefly, 150 µl (OD600 = 0.5-0.8) of S2208 cells were inoculated with 10 µl of serially diluted phage-containing media. Within 2 minutes, 0.9 ml of 50°C top agar (0.5% agar in 2xYT broth with 0.5 mg/ml X-gal) was added and the entire mixture was plated on a 60 mm petri dish containing 1.5% agar in 2xYT media with 50 µg/ml carbenicillin. *E. coli* contain the pJC175e plasmid (46) that provides gIII under the control of a phage shock promoter allowing for activity-independent expression of gIII upon infection.

Expression of gIII enables the generation of infectious progeny and the formation of plaques (45). The S2208 *E. coli* strain contains the LacZ gene under the control of a phage shock promoter in the genome for visualization of phage plaques. Plaques appear as blue spots on a bacterial lawn and were counted manually.

### Flipper flow cytometry assay

Helper plasmids encoding integrase variants (100 ng) and a flipper plasmid, pH1 EukFlip zsGreen PhiC31 Hygro, containing ZsGreen flanked by attB and attP sites (500 ng) were transfected into 24-well plates seeded with HEK293T cells as described above. Following three days of culture, cells were resuspended using cold Live Cell Imaging Solution (Thermo Fisher Scientific) and ZsGreen expression was assessed by flow cytometry (excitation 488 nm, emission 530 nm) of >20,000 live, single cells per condition using an Attune NXT flow cytometer (Thermo Fisher Scientific). Representative flow cytometry analysis can be seen in Supplementary Fig. S2a.

### Off-target analysis

Genomic PCR primers were generated to previously predicted Bxb1 pseudo sites (22,34). For each site, two primer sets were designed. The first (control) set was made to amplify the genome to verify ample template was used. The second set was made to amplify the junction of the genome and donor. Additional primers were generated to amplify successful insertion at ROSA26. Lysates from HEK293T transfections performed on two separate days were used as template for genomic PCR using the Q5 polymerase (NEB) and visualized on a 2% agarose gel.

Primers are listed in Supplementary Table S6.

### Detection of vWF expression

To measure expression and export of recombinant vWF protein, HEK293T cells were seeded in 6-well plates and the next day were transfected with 1200 ng of PE5max, 120 ng of each epegRNA plasmid, 480 ng of helper plasmid, 480 ng of donor plasmid, and 240 ng of DsRed expression plasmid (for sorting). A fraction of the cells was sorted by FACS three days post transfection to measure integration efficiency by ddPCR. Cells were cultured for 14 days to avoid transient expression. On day 14, media was changed to serum free media. On day 15, the media was harvested and vWF was detected using the vWF Human ELISA Kit (Invitrogen) according to manufacturer specifications.

## RESULTS

### PhiC31 integrase variants insert a 6.6 kb cargo DNA at safe harbor sites with prime editing

Previously, directed evolution was used to improve PhiC31 integrase activity on its native att sites (39). We reasoned that these evolved mutant variants of PhiC31 integrase could mediate targeted insertion into the genome at higher efficiency than wildtype. We first optimized PE for inserting the PhiC31 att sites to act as a landing pad for insertion. We chose previously characterized safe harbor loci that were distant from cancer-related genes and shown to permit expression of inserted genes, including AAVS1 (47), Xq22.1 (48), and ROSA26 (49). PE uses a fusion protein consisting of a Cas9 nickase and reverse transcriptase called PE5, as well as a prime editing guide RNA (pegRNA) (31). The pegRNA is a modified version of a single guide RNA (sgRNA) that both directs PE5 to the target genomic sequence and encodes the desired edit. The pegRNA contains a 5’ spacer sequence that directs PE5 to nick the target sequence (Supplementary Fig. S1a). The primer binding site (PBS) found on the 3’ end of the pegRNA anneals to the non-target DNA strand following the nick in the target strand. The pegRNA contains a reverse transcriptase template (RTT) that is reverse transcribed to generate a 3’ single-strand flap of DNA that encodes the inserted DNA, such as the att site. During twin PE, a second pegRNA directs the insertion of an adjacent flap that includes overlapping homologous sequence so that the two flaps form a double-stranded intermediate. Upon resolution of the intermediate, the new sequence becomes permanently inserted.

Because PE efficiency is generally higher for shorter inserts (1,34), we designed pegRNA for inserting the minimal PhiC31 attB site (35 bp) or attP site (41 bp) (37). To maximize PE efficiency, we used engineered pegRNA (epegRNA) containing a structured RNA motif to prevent degradation (22,32), an optimized cr772 guide scaffold (33), and expressed PE5 from the PE5max plasmid with an optimized editor architecture that includes a dominant negative mutant of human mutL homolog 1 (MLH1) involved in DNA mismatch repair which has been shown to enhance PE (31). PE5max and twin epegRNA expression plasmids for each target site were transfected into HEK293 cells.

PE insertion of att sites ranged from 0.3% to 62.1% as measured by amplicon sequencing using genomic primers flanking the insert (Fig. 1b) and confirmed by droplet digital PCR (ddPCR), a form of quantitative PCR that reports highly accurate copy number of genomic inserts over a wide range of template concentrations (Supplementary Fig. S1b) (50). We next generated clonal HEK293 cell lines with the most efficient and accurate epegRNA pairs targeted to AAVS1 and ROSA26, which we used to compare integration efficiency of PhiC31 integrase variants. Wildtype PhiC31 integrase was compared to three hyperactive variants called P1, P2, and P3 generated by Keravala et al. (39). Two of these three variants (P1 and P3) demonstrated increased activity measured by ddPCR relative to the wildtype integrase at both loci with integration efficiencies for P3 reaching 11.1% and 3.0% at ROSA26 and AAVS1, respectively (Fig. 1c). In place of using a cell line containing a pre-inserted att site, we next combined PE and integrase expression plasmids in a single transfection. This strategy allowed for the insertion of an att site using PE followed by the insertion of the donor plasmid by PhiC31 integrase at the target sequence (Fig. 1a). We co-transfected PE5max and epegRNA plasmids to insert the 41 bp PhiC31 attP site at ROSA26 along with a 6.6 kb donor plasmid containing the attB site and a helper plasmid expressing PhiC31 integrase. Encouragingly, we detected PhiC31 integrase insertion of the donor plasmid for all four integrase variants with a favorable efficiency of 2.3% for P3 compared to 0.7% for wildtype. These results demonstrated that hyperactive variants outperform wildtype PhiC31 integrase during PE mediated gene insertion (Fig. 1d). We also tested a range of attP site lengths delivered to ROSA26 and confirmed that 41 bp was optimal (Supplementary Note S2 and Supplementary Fig. S1c). Guided by these results, we reasoned that directed evolution of PhiC31 integrase variants might further increase the efficiency of gene insertion.

### Directed evolution of PhiC31 integrase

The previous directed evolution campaigns that generated PhiC31 P1, P2, and P3 required laborious steps of mutagenesis, gene expression, screening, and replication (39). Alternatively, PACE rapidly and iteratively selects, replicates, and mutates genes at rates of >70 generations per day without human intervention (35). PACE achieves selection by linking the desired activity of the gene of interest to the production of infectious phage progeny. Expression of M13 phage’s gene III (gIII), that encodes protein III (pIII), is proportionally related to proliferation. By removing gIII from the phage genome and placing it on a plasmid, gIII expression is solely driven by the desired activity of the evolving gene. Each selection phage (SP) encodes the gene to be evolved. Phage encoding active variants of the gene produce infectious progeny, and phage encoding less-active variants fail to produce as many progeny and become outcompeted. Beneficial mutations multiply with successful phage life cycles. An inducible mutagenesis plasmid (MP) increases the mutation rate during phage replication by decreasing proofreading and increasing error-prone bypass. IntePACE was devised as a PACE method for increasing the activity of integrase recombination on defined att sites. For the development of IntePACE, we encoded the integrase on a SP and linked gIII expression to a successful crossover event between the attB and attP sites. We placed a promoter and the N-terminus of gIII, called the leader sequence, followed by the attB site on a complementary plasmid (CP). A second plasmid called the accessory plasmid (AP), contained the attP site and the remaining portion of gIII (N1- C domains). During a crossover event between the attB and attP sites, the promoter and leader sequence join with gIII and subsequently leads to expression. The resulting attL site is encoded within a flexible loop so as not to disrupt the function of pIII (43). Successful events require precise and directional recombination to generate the attL site. During IntePACE, host cells containing the AP, CP, and MP are continually added to a fixed-volume flask (called the ‘lagoon’) where they encounter phage. After infection, host cells begin expressing the integrase protein from the SP. A successfully evolved integrase, bearing beneficial mutations enhancing its recombination activity, joins the promoter and gIII. Activated gIII expression leads to propagation of the SP that encoded the improved integrase. Due to the presence of the MP, these SP are again mutated during the next cycle of DNA replication, effectively generating a new library from the fittest variants of the previous library. Fresh host cells are then infected by this new library within the lagoon and the process is repeated. The proliferation of the phage scales with increasing levels of gIII expression over a range of two orders of magnitude (51). This establishes a competition between mutation library members resulting in the most active variants outcompeting the others (Fig. 2).

### IntePACE generates hyperactive PhiC31 integrase mutants

IntePACE experiments were initiated with SP encoding P1, P2, and P3 integrase and run for 212 hours (∼222 cycles of mutagenesis) (35). The activity of wildtype PhiC31 integrase in *E. coli* is relatively high, and recombination rates can approach 100% within an hour (52). Therefore, we continually increased the stringency of our system to challenge the evolving integrases by decreasing gIII promoter strength and increasing the flow rate (Fig. 3a). Host cells containing CP with four different gIII promoters were sequentially added to the lagoon. In order from strongest to weakest, promoters ProB, ProA, Pro3, and Pro1 (53) were used to challenge the evolving integrases. Because each recombination event produces low levels of pIII when using a weak gIII promoter, only highly active integrase variants capable of recombining more plasmids than less active variants can generate enough pIII to make sufficient infectious progeny to maintain in the system. Less active integrase variants fail to perform enough recombination events to produce sufficient numbers of progeny and are drained away. To further challenge the evolving integrases, the flow rate was increased from 2 to 3 lagoon volumes per hour, thereby reducing the time the integrase variants had to generate progeny. The presence of the phage in each lagoon was monitored by collecting the media outflow and performing the activity-independent plaque assay (46). Dilutions of media containing phage were plated on the S2208 *E. coli* strain that produces blue plaques (45) that were counted to calculate phage concentration. For each of the four host cell transfers, outflow media was collected at 0, 1, 2, and 4 hour time points and every 4 hours afterwards. We observed the expected pattern of an initial drop in titer as many of the early phage were drained. This was followed by a gradual increase in titer, presumably due to the integrase variants accumulating beneficial mutations. Next, we observed phage numbers plateau, likely due to the integrase variants reaching a point of hyperactivity that they were no longer challenged by the current level of stringency, necessitating increases in stringency for continued productive evolution. We next isolated phage from selected time points throughout the IntePACE experiment and separately measured recombination activity using the overnight phage enrichment assay. Cells containing the AP and CP using the ProA promoter to drive gIII were inoculated with 10^5^ PFU/ml of purified phage isolated from IntPACE. The cells were cultured in the presence of the phage overnight and the next day the phage concentration was measured using the activity-independent plaque assay, similar to above. Time points containing phage encoding hyperactive integrase variants are expected to recombine more AP and CP and generate more phage progeny than phage encoding less active variants. This method allows for individual time points to be directly compared. We observed evolved libraries increase efficiency by 15.4, 70.2, and 20.9-fold over P1, P2, and P3, respectively (Fig. 3b). To test activity in human cells, evolved integrase variants were isolated from single plaques and cloned into expression plasmids (Fig. 3c). We designed a plasmid-based ‘flipper’ assay in which a crossover event between the attP and attB sites flips the orientation of ZsGreen to join with the H1 promoter and activate expression (Supplementary Fig. S2a). In agreement with results from *E. coli*, we observed increased numbers of ZsGreen expressing cells for evolved integrases, with the greatest improvement by P2 from 9.9% to 44.9% (Supplementary Fig. S2b).

Evolved variants were compared in clonal HEK293 cell lines with a pre-inserted PhiC31 attP site at the AAVS1 (Supplementary Fig. S2c) or ROSA26 (Fig. 3d) locus. Encouragingly, gene insertion by evolved integrases outcompeted their starting variants. The greatest improvement was between P2 (1.3%) and a 36-hour evolved variant P2-L2-1 (12.1%). The highest integration rates came from a P3 variant (L1-2) that integrated 18.4% at the ROSA26 site. Similarly, evolved integrases co-transfected with PE components for attP insertion resulted in higher gene targeting efficiencies. Specifically, a 36-hour variant evolved from P2 (L1-10) increased by 10.9- fold and a 184 hour variant from P3 (L1-13) reached 5.0% efficiency at the ROSA26 target sequence (up from 2.3% from the P3 starting variant) (Fig. 3e). These results demonstrate that IntePACE is capable of generating novel integrase mutants with improved genomic insertion efficiency.

### IntePACE generates hyperactive Bxb1 integrase mutants

Using our optimized PE parameters for inserting att sites, we compared evolved PhiC31 integrase integration rates with previously described Bxb1 (22) and Pa01 (38) integrases. To our surprise, wildtype Bxb1 integrase had greater efficiency compared to evolved PhiC31 integrase mutants (Fig. 4a). We reasoned that Bxb1 integrase might serve as a superior starting point for evolution. Encouraged by our results with PhiC31 integrase, we adapted the same IntePACE strategy to increase Bxb1 integrase activity on its native att sites. The full Bxb1 attP site and minimal attB site (22,54) were substituted for the PhiC31 att sites in the AP and CP respectively (Supplementary Table S4). In total, phage were evolved for 9 days (∼276 cycles of mutagenesis) (35) (Fig. 4b). Similar to IntePACE with PhiC31 integrase, enrichment assays demonstrated that evolved variants of Bxb1 integrase vastly improved in the bacterial system (Supplementary Fig. S3a). New variants were cloned into mammalian expression plasmids which were then co-transfected with a donor plasmid into a clonal HEK293 cell line containing a pre-inserted Bxb1 attB site at ROSA26. We observed efficiencies up to 21.0% with the highest activity variant (Bxb1-L8-5) increasing 2.5-fold over wildtype Bxb1 integrase (Fig. 4c and Supplementary Fig. S3b). Using unmodified HEK293 cells, simultaneous PE and evolved Bxb1 integrase-mediated gene insertion resulted in efficiencies ranging from 2.1%-35.4%, up to a 3.1-fold increase in efficiency over wildtype Bxb1 integrase (Fig. 4d). Additional rounds of evolution did not result in variants with increased efficiency (Supplementary Fig. S4 and Supplementary Note S5).

### Bxb1 integrase variants with combinations of selected IntePACE mutations increase hyperactivity

Five mutations from top-performing Bxb1 integrase variants isolated from IntePACE were assembled in all combinations to generate 32 new mutants in total. To compare activity between mutants independent of PE, helper plasmids encoding combination mutants were co-transfected with a donor plasmid into a clonal HEK293 cell line with a pre-inserted attB site at ROSA26. Encouragingly, combination mutants increased activity over wildtype Bxb1 integrase by up to 9.2-fold (Fig. 5a). The different mutations had a variety of synergistic and antagonistic effects on one another. Of the five mutations, I87L and H95Y had the greatest individual effect on integration efficiency (7.2 and 6.4-fold higher than wildtype Bxb1 integrase, respectively), but when combined they were less effective (4.7-fold higher than wildtype Bxb1 integrase).

**Fig. 5.**
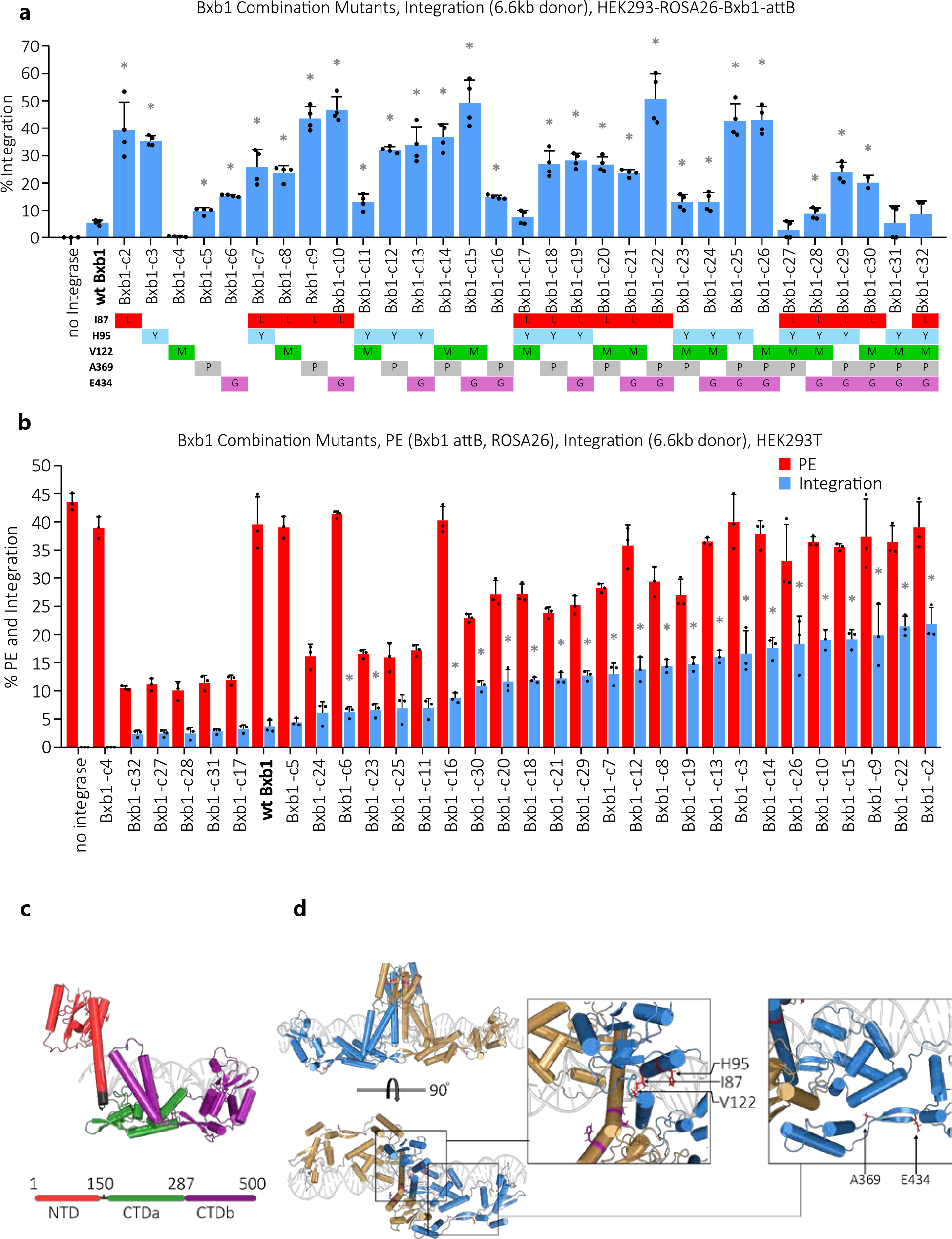
**Bxb1 integrase combination mutants. a**, Five beneficial mutations were selected from IntePACE for further characterization. Insertion efficiency measured by ddPCR of integrases with all combinations of one to five mutations was compared in a clonal HEK293 cell line containing a Bxb1 attB site at ROSA26. n = 4. **b**, PE efficiency of Bxb1 attB site insertion and integration efficiency of a donor plasmid was measured by ddPCR to compare combination mutants in unmodified HEK293 cells. n = 4. Data are shown as mean + s.d. (*) = P value < 0.05 derived from a Student’s two-tailed t-test if significantly higher than the pre-evolved integrase control **c**, Alphafold model of Bxb1 integrase. Bxb1 integrase domains defined by Ghosh et al. (52). Pymol was used to align with LI prophage integrase (pdb 6dnw) co-crystalized with attP half-site DNA (69). **d**, Alphafold model of Bxb1 integrase dimer aligned with LI prophage integrase to visualize DNA. The five mutations selected for combination variants are shown in red and purple.

Mutations A369P and E434G had a modest effect on integration activity on their own but exhibited a synergistic effect with every other mutation tested, including V122M, which decreased activity relative to wildtype Bxb1 integrase on its own.

We next used PE to insert the attB site at ROSA26 in HEK293 cells by co-transfecting PE plasmids with combination mutant helper plasmids and a donor plasmid. When combined with PE, combination mutants demonstrated increased activity over wildtype Bxb1 integrase up to 6-fold (Fig. 5b). Different patterns of activity were observed for the combination mutants between experiments performed with and without PE. For example, Combo 25 had high activity on the clonal cell line and low activity when paired with PE, indicating that the mutations had a negative effect on PE which in turn decreased the integrase’s insertion efficiency despite being hyperactive. Similarly, in all cases where H95Y and V122M were combined, PE efficiency decreased significantly (2.3 to 4-fold relative to wildtype Bxb1 integrase). Anzalone et al. previously showed that the att site found on the epegRNA expression plasmid could recombine with the att site on the donor plasmid, rendering both plasmids nonfunctional (22). This is expected to reduce the amount of epegRNA available for PE as well as reduce the number of donor plasmids available for integrase insertion. To reduce the effect of Bxb1 integrase recombining these plasmids, our epegRNA expression plasmids contain a partial att site. During twin PE, although each inserted 3’ single strand flap contains only part of the att site, there is enough overlapping sequence to form a double-stranded intermediate. The remaining single-stranded sequence of the att sites can subsequently be filled in to promote insertion of the full att site (22). While this strategy appears sufficient for preventing unwanted wildtype Bxb1 integrase recombination, we speculate that certain hyperactive integrase mutants gained activity on the partial att site sequence and recombined the epegRNA expression plasmids, which may explain the reduction in PE efficiency.

Alphafold (55,56) modeling of Bxb1 integrase as a dimer (57) revealed that mutations I87L, H95Y and V122M are in close proximity to each other on the N-terminal catalytic domain. Mutations A369P and E434G are located together proximal to the coiled-coiled motif which has been implicated in the regulation interactions between monomers of the tetrameric interface during synapsis. (Fig. 5c and 5d) (58,59). Further studies are necessary to elucidate how these individual mutations improve activity.

We observe higher transfection rates for HEK293T compared to HEK293 cells. Top variants were re-assayed using HEK293T cells resulting in insertion efficiencies up to 44.1%, with combination mutants outperforming parent variants (Supplementary Fig. S5a). Furthermore, no off-target insertions were detected by genomic PCR at predicted Bxb1 pseudo sites (22,34) (Supplementary Fig. S5b). Combination mutants were evolved using IntePACE but variants with further increases in efficiency were not recovered (Supplementary Fig. S6).

### Hyperactive integrases outperform wildtype Bxb1 integrase fused to prime editing components

Yarnall et al. described the PASTE system that uses single pegRNA PE and a fusion of wildtype Bxb1 integrase to the Cas9 nickase and reverse transcriptase (34). Similar to others, we observed lower PE efficiencies using a single pegRNA (22,60), with PASTE PE and integration into the ROSA26 locus reaching 11.2% and 2.4%, respectively. We improved PASTE by incorporating twin PE pegRNA (22) and observed that the fused integrase architecture indeed outperformed traditional twin PE with wildtype Bxb1 integrase. However, neither traditional PASTE nor twin PE PASTE reached the levels of our evolved Bxb1 integrase. We introduced several hyperactive mutations into both PASTE and twin PE PASTE but did not observe a further increase in efficiency (Supplementary Fig. S7).

### Evolved integrases function in multiple human cell types and insert large therapeutically relevant DNA cargo at ROSA26

To exclude the influence of incomplete transfection efficiency, top evolved variants were assayed in HEK293T cells and sorted for DsRed expression. These optimized conditions resulted in insertion rates as high as 80% of target sites (Fig. 6a). Evolved Bxb1 integrase variants were active on HeLa, U2OS human osteosarcoma, and K562 human lymphoblast cell lines as well as primary human fibroblasts. Surprisingly, although PE was efficient in HeLa cells (>40%), we were unable to detect integration using wildtype Bxb1 integrase. However, evolved integrases successfully performed integration, albeit at low levels (Fig. 6b). In U2OS cells, evolved integrases outperformed wildtype Bxb1 integrase by 6.7-fold, and in K562 cells, the improvement was a remarkable 11.2-fold (2.7% vs 30.3%) (Fig. 6c and 6d). Although we were only able to PE ∼15% of primary HDFa cells, normalized integration by the hyperactive Bxb1-c22 integrase represented 35.2% of PE cells (Supplementary Note S6 and Supplementary Table S3) and was a 2-fold improvement over wildtype Bxb1 integrase (Fig. 6e).

**Fig. 6.**
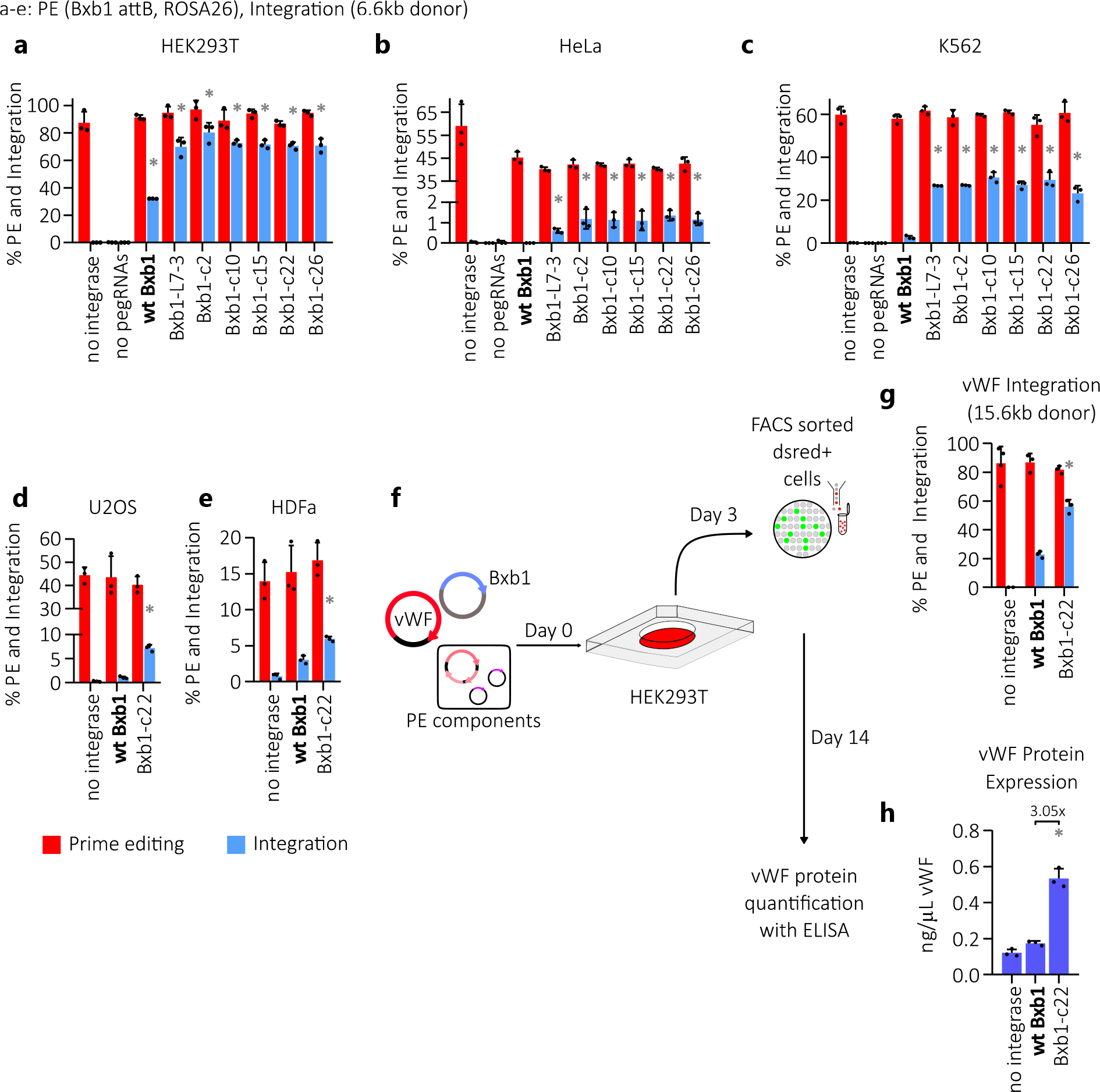
**Insertion of large cargos by evolved Bxb1 integrase with PE. a-e**, PE efficiency of insertion of the att site and integrase insertion efficiency from combination mutants (Fig. 5a) of a 6.6 kb donor plasmid during optimized transfections of (**a**) HEK293T (**b**) HeLa (**c**) K562 (**d**) U2OS and (**e**) HDFa cells. Cells were sorted three days post transfection for DsRed expression to exclude non-transfected cells before analysis by ddPCR. n = 3. **f**, Schematic depicting delivery of therapeutic cargo to HEK293T cells. Cells were transfected with PE components, integrase helper plasmid, and a 15.6 kb vWF donor plasmid. Cells were sorted for DsRed expression three days post transfection to exclude non-transfected cells. Half the cells were pelleted for ddPCR detection of vWF donor integration at ROSA26 (**g**) and the remaining cells were cultured for an additional 11 days to minimize transient expression of the therapeutic transgene. n = 3. **h**, Detection of secreted vWF in culture media by ELISA. Data are shown as mean + s.d. (*) = P value < 0.05 derived from a Student’s two-tailed t-test if significantly higher than the pre-evolved integrase control (shown in bold on the left).

Integrase insertion of large cargos would enable therapies inaccessible to current methods. Specifically, the full-length human vWF gene (9.3 kb cDNA) is not amenable for HDR (61), AAV (62), or lentiviral delivery (63). We measured PE and integrase insertion of a 15.7 kb cargo encoding vWF and observed 56.2% integration by the evolved Bxb1 integrase compared to 22.8% for wildtype Bxb1 integrase. Following 15 days of culture to prevent detection of non-integrated, transient vWF expression, we detected a 3-fold increase in protein expression for the evolved integrase (Fig. 6f-6h). Collectively, these data show that evolved integrases generated by IntePACE outperform their wildtype counterparts in multiple cell types and can efficiently insert large therapeutic cargos into the human genome.

## DISCUSSION

Genome engineering by integrases does not rely on host repair and is highly specific for the att sequence that is not present in the unmodified human genome (22). By pre-inserting the att site into cell lines, hyperactive integrases could mediate highly efficient gene insertion to a single known locus. Cell lines containing a pre-inserted att site could be used to rapidly generate clonal lines for producing monoclonal antibodies or other Good Manufacturing Practice (GMP)- grade products (6).

For therapeutic applications in cells without a pre-inserted att site, PE is an attractive tool for delivering the att site to a desired locus. PE functions in both dividing and non-dividing cells and efficiently inserts sequences <50 bp while minimizing DSBs (1). Simultaneous PE and integrase insertion supports large DNA cargo delivery to nearly any genomic target sequence (22).

Therapeutics would benefit from improved integration efficiencies greater than current wildtype enzymes. For example, CAR-T cancer immunotherapy is often limited by the amount of the patients’ donated T-cells for *ex-vivo* editing (64,65). Higher numbers of accurately edited cells available for re-introduction to the patient could improve prognosis.

In this study we developed IntePACE, a continuous directed evolution approach that we used to rapidly generate hyperactive integrases with targeting efficiencies that greatly exceed current approaches. A key feature of IntePACE is the tethering of the generation of phage progeny to successful recombination of att sites. We devised a strategy for stepwise increases in stringency to continually challenge evolving integrases to promote increasing activity. We reasoned that tuning the system by applying selective pressure would increase the speed and frequency of recombination that are required by the evolving integrases. Following IntePACE, we observed increased phage production that was used as an indicator of recombination activity from evolved PhiC31 integrases compared to wildtype integrases in *E. coli*. We used a fluorescent protein-based assay to show that selected mutants outperformed their unevolved starting variants in human cells. Similarly, by co-transfecting PE components for the insertion of the PhiC31 att site into the genome, we demonstrated improvements to genome integration of a 6.6 kb donor plasmid by evolved PhiC31 integrases compared to wildtype. This represents the first report using PhiC31 integrase in conjunction with PE. Because PE precedes integrase genome insertion, high efficiency PE is crucial, and careful choice of the target sequence and epegRNA is necessary. By optimizing PE, we identified two promising target sequences in AAVS1 and ROSA26. Should another target region be required, we suggest testing several epegRNA and using amplicon sequencing to exclude sequences that cause indel formation. Although IntePACE successfully improved PhiC31 integrase activity, we observed higher rates for wildtype Bxb1 integrase, which prompted us to perform IntePACE on Bxb1 integrase in an attempt to isolate hyperactive mutants. Similar to PhiC31 integrase experiments, selected mutants from Bxb1 IntePACE outcompeted their pre-evolved variants in *E. coli* and human cells. By combining select integrase mutations, we observed insertion efficiencies of up to 80% of target sequences. In a therapeutic context, given that normal cells contain a target site on each of two chromosomes, this efficiency equates to an estimated 96% probability that a cell will receive at least one insertion (Supplementary Note S3). This boost in efficiency did not result in measurable off-target insertions at predicted pseudo-sites (Supplementary Fig. S5b). We theorize that improvements in delivery and further optimization of DNA/RNA components would increase integration rates even further. For example, Anzalone et. al, demonstrated that delivery of pegRNA as RNA increases PE and integration rates, most likely due to preventing interactions between attP and attB sites on the pegRNA expression plasmids and the donor plasmid (22).

Additionally, the CMV promoter (which drives our integrase expression plasmid) has shown to be less effective in HeLa cells relative to HEK293 cells in which we did our optimizations (66). Nevertheless, in every cell line tested, IntePACE evolved integrases outperformed the wildtype integrase that is the current benchmark for site specific delivery of large DNA cargos to the human genome (22,34). A challenge to this technology, that is shared by most other genome engineering approaches, is efficient plasmid delivery. We observed relatively low PE (∼15%) for primary HFDa cells which limited potential integration. We observed variable integration efficiencies for the cell types tested which likely was influenced by different transfection efficiencies for the diverse lines. Technologies that improve delivery will be crucial for boosting both PE and integration in challenging cell types and tissues. Notwithstanding, a key finding was that evolved integrases outcompeted wildtype integrases particularly well during pre-optimized, suboptimal transfection conditions (>6 fold) (Fig. 5a and 5b) indicating that this technology might be suited for *in-vivo* gene delivery where tissue transfection efficiency can be difficult. In the K562 lymphoblast cell line in particular, evolved integrases had a 11-fold increase in targeting efficiency over wildtype. Von Willebrand Disease, the most common blood clotting disorder affecting 1% of individuals worldwide, is caused by mutations in vWF (67). Delivery of large therapeutic genes such as vWF can be intractable for current methods. We demonstrated that our evolved integrase efficiently inserted a 15.7 kb cargo encoding a functional vWF gene.

Mining of metagenomes has revealed alternate integrases with high efficiency (34,38). The directed evolution method described here could easily be adapted to further improve these integrases analogous to PhiC31 and Bxb1 integrases. Furthermore, IntePACE could be adapted to evolve integrases with increased specificity on pseudosites found in the human genome eliminating the need for coupling integrase insertion to other gene editing technologies for the delivery of landing pad att sites (68). We envision that this highly efficient technology will be tailored to diverse research tools and future therapies.

## DATA AVAILABILITY

The data underlying this article are available in the article and in its online supplementary material.

## SUPPLEMENTARY DATA

Supplementary Data are available at NAR online.

## AUTHOR CONTRIBUTIONS

Conceptualization: B.E.H., S.G., R.S., A.H.B., J.B.O. Methodology: B.E.H., S.G., R.S., A.H.B., J.B.O.

Investigation: B.E.H., S.G., R.S., D.F.W., I.S., J.E.S., L.S., C.T.T. Visualization: B.E.H, S.G., R.S., J.B.O.

Funding acquisition: J.B.O. Project administration: J.B.O. Supervision: J.B.O. Writing – original draft: B.E.H., S.G., R.S., J.B.O. Writing – review & editing: B.E.H., S.G., R.S., A.H.B., J.B.O.

## ACKNOWLEDGEMENTS

We thank all members of the Owens Laboratory for helpful discussions and the core facilities at the University of California – Davis and University of Hawaii for technical support.

