## Supplemental Files for "Directed evolution of hyperactive integrases for site specific insertion of transgenes"

†Joint Authors.

Supplementary Notes S1-S6

Supplementary Figures S1-S8

Supplementary Tables S1-S11

**Supplementary Note S1.**

**Rationale for normalization using the product/reference ratio (PRR).**

Two options exist for the location of the target probe: 1) The probe can be placed in the donor, with the forward primer in the donor and the reverse primer in the genome, or 2) The probe can be placed in the genome, with the forward primer in the donor and the reverse primer in the genome. An issue with the donor-located probe is that insertion at an off-target sequence positions the probe and reverse primer downstream of an unknown genomic sequence. Mispriming from the genome could generate a false signal. A second issue is that a donor-located probe cannot be used to determine the ratio of potential target sequences to reference sequences. If a greater number of target sequences exist than reference sequences, then a greater number of opportunities exist for targeting, and the final targeting efficiency will be over-reported. Accounting for this issue is especially important when assaying cell lines with abnormal numbers of chromosomes and where the reference sequence often is located on a different chromosome than the target sequence. To prevent ddPCR over- or under-reporting, we have defined the ratio of target product to reference signal as the product/reference ratio (PRR). Targeting efficiencies for unknown samples are divided by the PRR to normalize for differences in available targets and references. To determine the PRR, untransfected cell lines were assayed using FAM probes designed to bind the genomic target sequence. Unlike typical ddPCR used for identifying the target, both the forward and reverse primers are located in the genome flanking the probe. The ddPCR is run with standard HEX probes designed to bind the genomic reference sequence that also are flanked by genomic primers. The value from the FAM target is divided by the HEX reference to give the PRR. Subsequent ddPCR reactions are normalized by dividing their values by the PRR.

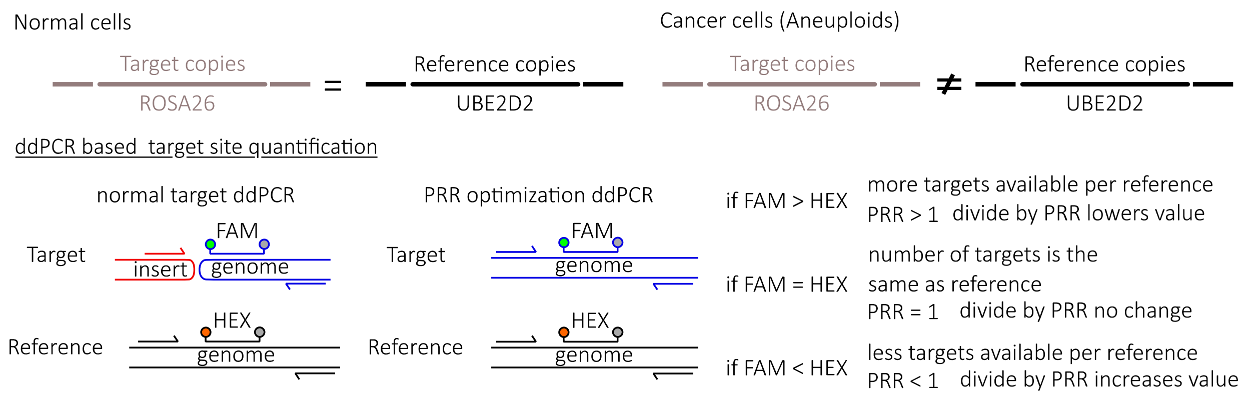

| ref | cell line | target | ave PRR | stdev |
|---|---|---|---|---|
| RPP30* | HEK293 | ROSA 1a | 0.72 | 0.034 |
| | HEK293 | ROSA 3a | 0.76 | 0.037 |
| | HEK293 | Xq22 1a | 1.01 | 0.087 |
| | HEK293 | AAV 3a | 0.47 | 0.007 |
| | HEK293 | AAV 2a | 0.43 | 0.007 |
| | HEK293 | AAV 1a | 0.42 | 0.017 |
| UBE2D2** | HEK293 | ROSA 3a | 1.01 | 0.006 |
| | HEK293 | AAV 3a | 0.63 | 0.017 |
| | HEK293T | ROSA 3a | 1.22 | 0.011 |
| | HeLa | ROSA 3a | 0.92 | 0.008 |
| | K562 | ROSA 3a | 1.24 | 0.004 |
| | U2OS | ROSA 3a | 1.54 | 0.062 |
| | HDFa | ROSA 3a | 0.96 | 0.054 |
| | HEK293 clone | PhiC31 attP41 ROSA 3a | 0.67 | 0.004 |
| | HEK293 clone | PhiC31 attP41 AAV 3a | 0.65 | 0.004 |
| | HEK293 clone | Bxb1 attB38 ROSA 3a | 0.38 | 0.011 |

**Supplementary Table S1. PRR values for target sequences used in this work.** Templates were lysates isolated from wells of untransfected cells from the indicated cell lines. n = 3. HEK293 clones were first edited by prime editing (PE) for insertion of the indicated att site. Probes were designed to bind the genomic target sequence. Primers for the HEK293 clones included the forward primer in the att site and the reverse primer in the genome. Primers for the remaining cell lines included the forward primer and reverse primer in the genome. * Reference used in Supplementary Fig. S1b. ** Reference used for all other figures.

**Supplementary Note S2. Minimal PhiC31 attB and attP sites.** PE components were co-transfected into HEK293T cells with a PhiC31 P3 integrase helper expression plasmid and a 6.6 kb donor plasmid. The reverse transcriptase templates included variable length attB and attP sites on the epegRNA expression plasmids. Each experiment used a donor containing the full att site. Prime editing and integration rates were measured by ddPCR after three days. Optimal PE and integration was observed for the attP site measuring 41bp that was used for further optimization of PhiC31 integrase-based DNA cargo delivery (Supplementary Fig. S1c).

### Supplementary Note S3.
### Probability of at least one integration per cell.

Given an insertion efficiency of 80% of available targets and two targets/cell.

P(≥1 insert/cell) = 1 – P(0 inserts/cell)

P(0 inserts/cell) = P(miss target 1) x P(miss target2)

P(miss target) = 1 – P(hit target) = 1 – 0.80 = 0.20

∴ P(0 inserts/cell) = 0.20 x 0.20 = 0.04

∴ P(≥1 insert/cell) = 1 – 0.04 = 0.96 = 96%

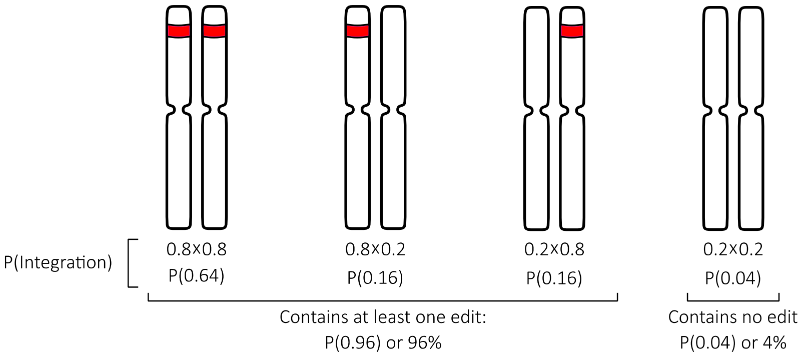

**Supplementary Note S4.**
**IntePACE optimizations.**
*PhiC31 Integrase AP/CP Ori Matrix*. The addition of the CP required optimization of a two-plasmid system in which successful recombination by an active integrase merges the AP and CP into one large plasmid with two origins of replication. To determine which combinations of origins best promote phage production, all compatible combinations of seven origins (each with a different copy number) were tested in phage enrichment assays using an evolved PhiC31 mutant phage. Phage titers after overnight enrichment spanned four orders of magnitude allowing for tunable stringency for IntePACE (Supplementary Fig. S8e).

*Changing SP gVI promoter to the RecA promoter decreases occurrence of recombinant wildtype M13 contamination*. Early attempts to run IntePACE on PhiC31 resulted in recombinant wildtype M13 phage "poisoning" the lagoons. This occurred when gIII from the AP/CP infiltrated the SP (typically within the gene of interest). While the N-terminus of the gIII insertion into the SP was variable, the C-terminus of gIII matched perfectly with the gVI promoter region which is an exact match to the C-terminal 183 bp of gIII. Replacement of the gVI promoter region with the bacterial RecA promoter maintained similar activities in phage propagation experiments (Supplementary Fig. S8b) and prevented further occurrences of recombinant wildtype phage.

*IntePACE gIII linker reduces leaking and activity independent phage propagation*. In a previous IntePACE experiment with PhiC31 integrase, leakiness of the gIII led to low levels of activity independent phage propagation. To alleviate this issue, gIII was separated at the linker region between the N-terminal leader sequence (LS) and the remaining truncated gIII C-terminus (N1-c). A promoter and the LS were moved to a separate plasmid (CP) immediately adjacent to the attB site. The attP site followed immediately by the N1-C end of gIII remained on the AP. Successful recombination of these two plasmids by an active integrase aligns the two parts of gIII, expressing the intact pIII with a small peptide insertion encoded by the newly formed attL site. To test this system, overnight enrichment assays were performed using SP containing the S12F catalytically inactivated PhiC31 integrase (1). The AP with the fully intact gIII produced 183-fold more phage than the split gIII linker system (Supplementary Fig. S8c).

*Increased stringency in PACE host cells achieved by decreasing gIII promoter strength*. Insulated promoters of variable strengths (2) were cloned into CP plasmids driving the expression of the gIII LS and tested with a single AP in an overnight phage enrichment assay. ProB produced the most phage of the promoters tested, which was used to start both PhiC31 and Bxb1 PACE. The range of phage titers resulting from different gIII promoters spanned five orders of magnitude, allowing for an additional layer of tunable stringency for IntePACE (Supplementary Fig. S8a).

*High volume PACE allows for more lagoons and higher flow rates*. Two chemostats of 2L each containing host cells were maintained at volumes ranging from 200-250 ml. Because the MP can become inactive over time due to the accumulation of mutations, the chemostats were changed every two days to maintain arabinose sensitivity. A 500 ml mixer was maintained at a volume of 50-100 ml and used to mix the two cultures before distribution to 2-6 100 ml lagoons maintained at 40 ml (except when high flow rates were used to increase stringency and volumes were reduced to compensate). L-Arabinose was supplied to the lagoons at a constant rate of 0.5 µl/min. The concentration was dependent on flow rate to maintain 100 mM L-arabinose in the lagoon (Supplementary Fig. S8f).

| PACE | Time | AP | CP | Flow Rate | Starting material |
|---|---|---|---|---|---|
| PhiC31 PACE | 0-72hr | AP pUC PhiC31 attP | CP RSF1030 PhiC31 attB ProB | 2 V/hr | Lagoon 1 SP RecA PhiC31 (wt) |
| | 72-120hr | AP pUC PhiC31 attP | CP RSF1030 PhiC31 attB ProA | 2 V/hr | Lagoon 2 SP RecA PhiC31 P2 |
| | 120-168hr | AP pUC PhiC31 attP | CP RSF1030 PhiC31 attB Pro3 | 2 V/hr | Lagoon 3 SP RecA PhiC31 P1 |
| | 168-192hr | AP pUC PhiC31 attP | CP RSF1030 PhiC31 attB Pro1 | 2 V/hr | Lagoon 4 SP RecA PhiC31 P1 |
| | 192-212hr | AP pUC PhiC31 attP | CP RSF1030 PhiC31 attB Pro1 | 3 V/hr | Lagoon 5 SP RecA PhiC31 P3 |
| | | | | | Lagoon 6 SP RecA PhiC31 P3 |
| Bxb1 PACE 1 | | | | | Lagoon 1 SP RecA Bxb1 (wt) |
| | 0-24hr | AP pUC Bxb1 attP | CP RSF1030 Bxb1 attB ProB | 2 V/hr | Lagoon 2 SP RecA Bxb1 (wt) |
| | 24-48hr | AP pUC Bxb1 attP | CP RSF1030 Bxb1 attB ProA | 2 V/hr | Lagoon 3 SP RecA Bxb1 (wt) |
| | 48-72hr | AP pUC Bxb1 attP | CP RSF1030 Bxb1 attB Pro3 | 2 V/hr | Lagoon 4 SP RecA Bxb1 (wt) |
| | 72-124hr | AP pUC Bxb1 attP | CP RSF1030 Bxb1 attB Pro1 | 2 V/hr | Lagoon 5 SP RecA Bxb1 (wt) |
| | | | | | Lagoon 6 SP RecA Bxb1 (wt) |
| Bxb1 PACE 2 | 0-24hr | AP pUC Bxb1 attP | CP RSF1030 Bxb1 attB Pro1 | 3 V/hr | Lagoon 7 SP RecA Bxb1 PACE1 L4-120hr |
| | 24-48hr | AP pUC Bxb1 attP | CP RSF1030 Bxb1 attB Pro1 | 4 V/hr | Lagoon 8 SP RecA Bxb1 PACE1 L6-100hr |
| | 48-72hr | AP pUC Bxb1 attP | CP RSF1030 Bxb1 attB Pro1 | 6 V/hr | |
| Bxb1 PACE 3 | 0-24hr | AP pUC Bxb1 attP | CP RSF1030 Bxb1 attB Pro1 | 2 V/hr | Lagoon 9 SP RecA Bxb1 PACE2 pool |
| | 24-48hr | AP pUC Bxb1 attP | CP RSF1030 Bxb1 attB Pro1 | 4 V/hr | Lagoon 10 SP RecA Bxb1 L2-9 |
| | 48-72hr | AP pUC Bxb1 attP | CP RSF1030 Bxb1 attB Pro1 | 6 V/hr | |
| | 72-96hr | AP pUC Bxb1 attP | CP RSF1030 Bxb1 attB Pro1 | 8 V/hr | |
| | 96-98hr | AP pUC Bxb1 attP | CP RSF1030 Bxb1 attB Pro1 | 10 V/hr | |
| Bxb1 PACE 4 | | | | | Lagoon 11 SP RecA Bxb1 (wt) |
| | 0-48hr | AP R6k Bxb1 attP | CP RSF1030 Bxb1 attB Pro1 | 2 V/hr | Lagoon 12 SP RecA Bxb1 Combo 2 |
| | 48-72hr | AP R6k Bxb1 attP | CP RSF1030 Bxb1 attB Pro1 | 3 V/hr | Lagoon 13 SP RecA Bxb1 Combo 3 |
| | 72-96hr | AP R6k Bxb1 attP | CP RSF1030 Bxb1 attB Pro1 | 4 V/hr | Lagoon 14 SP RecA Bxb1 Combo 9 |
| | | | | | Lagoon 15 SP RecA Bxb1 Combo 22 |
| | | | | | Lagoon 16 SP RecA Bxb1 L7-3 |

**Supplementary Table S2. Summary of PACE experiments.** A total of five IntePACE experiments were performed. The results of PhiC31 PACE are represented in Figure 3a. The results of Bxb1 PACE 1 and 2 are shown in Figure 4b, Bxb1 PACE 3 in Supplementary Fig. S4a, and Bxb1 PACE 4 in Supplementary Fig. S6b.

**Supplementary Note S5. Bxb1 PACE 3 did not result in improved variants in human cells.** A third Bxb1 PACE experiment was conducted in an attempt to further increase the activity of evolved Bxb1 mutants. A single phage clone from PACE 1 (Bxb1-L2-9) and a pool of time points from PACE 2 were evolved in host cells containing the weakest promoter (Pro 1) for 98 hours increasing the flow rate each day (Supplementary Fig. S4a). Although increased integrase activity was observed in bacterial phage propagation assays (Supplementary Fig. S4b), no mutants selected from these pools demonstrated increased activity in human cells (Supplementary Fig. S4c and 4d).

**Supplementary Note S6. Summary of PE and integration rates in different cell lines.** Multiple cell lines were simultaneously transfected with PE components and unevolved or evolved integrases to insert a donor plasmid at the ROSA26 locus with efficiencies ranging from 1-70%. In every cell line transfected, the evolved Bxb1-c22 integrase facilitated higher rates of integration compared to wildtype Bxb1 integrase. Normalizing the integration rate to the PE efficiency demonstrates that the efficiencies of integrase activity and prime editing activity vary independently between cell types.

| | | Bxb1 | | | Bxb1-c22 | | | fold increase |
|---|---|---|---|---|---|---|---|---|
| Figure | Cell line | % PE | % Integration | Int/PE | % PE | % Integration | Int/PE | Bxb1-c22/Bxb1 |
| 5b | HEK293 | 39.57 | 3.66 | 9.25% | 36.49 | 21.44 | 58.76% | 5.9 |
| 6a | HEK293T | 91.32 | 32.09 | 35.14% | 86.67 | 70.58 | 81.44% | 2.2 |
| 6b | HeLa | 45.20 | 0.00 | 0.00% | 40.33 | 1.36 | 3.37% | undefined |
| 6c | K562 | 57.95 | 2.71 | 4.68% | 55.04 | 29.33 | 53.29% | 10.8 |
| 6d | U2OS | 43.49 | 1.08 | 2.48% | 40.26 | 7.22 | 17.93% | 6.7 |
| 6e | HDFa | 15.23 | 3.05 | 20.03% | 16.87 | 5.95 | 35.27% | 2.0 |

**Supplementary Table S3. Summary of PE and integration rates in different cell lines.** % PE and % Integration are averages of at least 3 biological replicates taken from data shown in the figures listed in the column on the left. The fold increase refers to the total integration rate of the Bxb1-c22 mutant vs. wild type Bxb1.

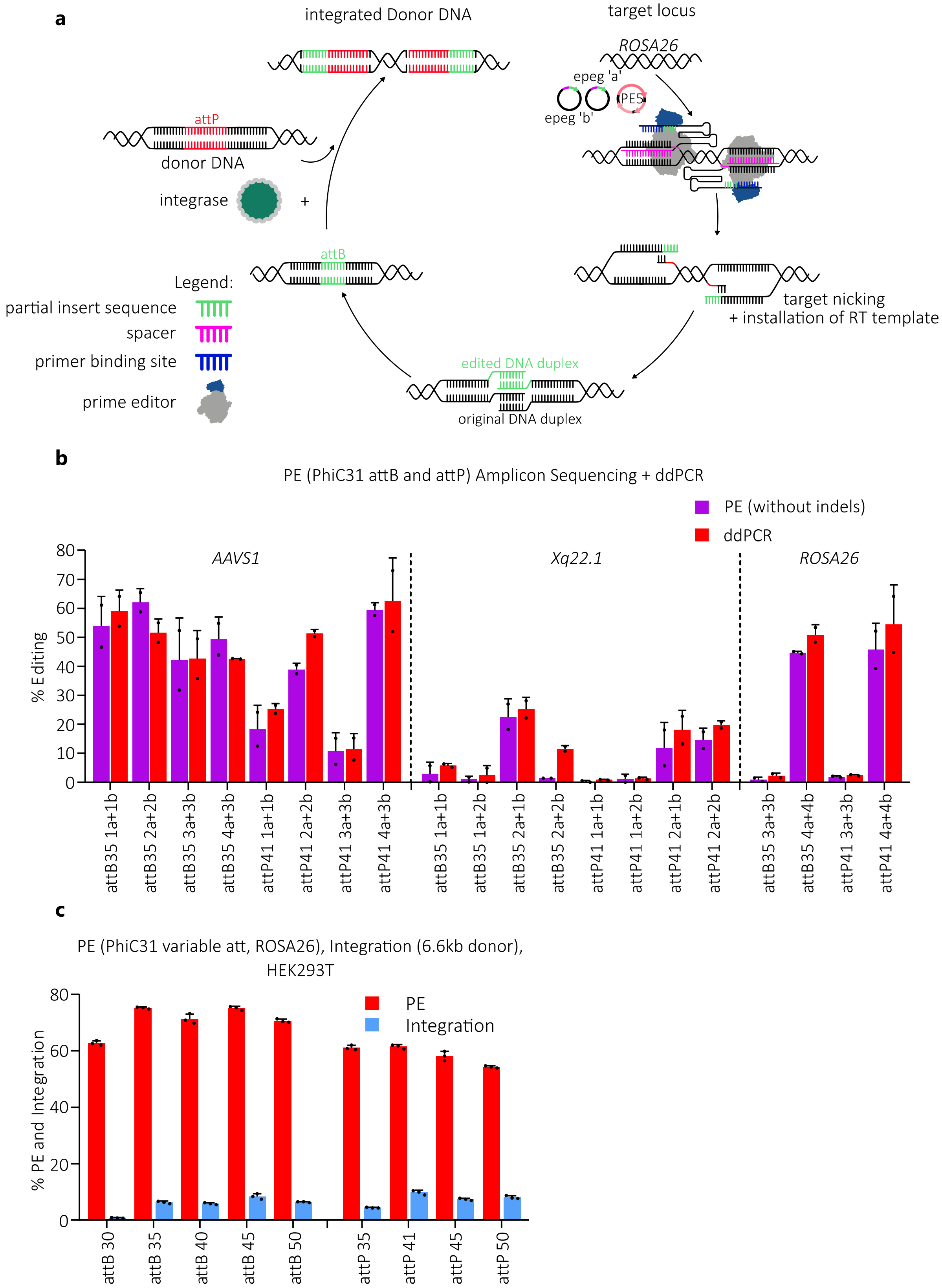
a
integrated Donor DNA
target locus
ROSA26
epeg 'a'
epeg 'b'
PE5
attP
donor DNA
integrase
+
attB
Legend:
partial insert sequence
spacer
primer binding site
prime editor
target nicking
+ installation of RT template
edited DNA duplex
original DNA duplex
b
PE (PhiC31 attB and attP) Amplicon Sequencing + ddPCR
PE (without indels)
ddPCR
% Editing
AAVS1
Xq22.1
ROSA26
attB35 1a+1b
attB35 2a+2b
attB35 3a+3b
attB35 4a+3b
attP41 1a+1b
attP41 2a+2b
attP41 3a+3b
attP41 4a+3b
attB35 1a+1b
attB35 1a+2b
attB35 2a+1b
attB35 2a+2b
attP41 1a+1b
attP41 1a+2b
attP41 2a+1b
attP41 2a+2b
attB35 3a+3b
attB35 4a+4b
attP41 3a+3b
attP41 4a+4b
c
PE (PhiC31 variable att, ROSA26), Integration (6.6kb donor), HEK293T
% PE and Integration
PE
Integration
attB 30
attB 35
attB 40
attB 45
attB 50
attP 35
attP 41
attP 45
attP 50

**Supplementary Fig. S1. Optimization of PE at genomic safe harbor sites. a,** Schematic of twin PE and integrase insertion (adapted from Anzalone et al.)(3). The prime editor (PE5) consists of the SpCas9 nickase fused to a reverse transcriptase. Spacers designed to target opposite strands generate nicks in the DNA by PE5. The epegRNA serves as template for reverse transcription extending from the nick site for insertion of new sequence. Resulting single strand flaps anneal to form an edited DNA duplex that is resolved to generate the desired insertion of the attB site. The attB site now found in the genome is recombined with an attP site found on a donor plasmid by the integrase, resulting in insertion of the donor plasmid. **b**, Amplicon sequencing compared with ddPCR results for detection of PE inserted att sites at AAVS1, Xq22.1, and ROSA26. Cells were transfected with PE5max and two epegRNA expression plasmids (epegRNA sequences are listed in Supplementary Table S11). Products for amplicon sequencing were generated from primers flanking the genomic target site. A forward primer and probe located in the genome as well as a reverse primer in the att site were used for ddPCR. All ddPCR measurements were normalized using the product/reference ratio (PRR) to account for differences in the number of target sequences in the genome of cell lines (Supplementary Note S1 and Supplementary Table S1). n = 2. **c**, Comparison of PE and integration efficiency using different PhiC31 attB and attP site lengths. epegRNA pairs designed to insert PhiC31 attB or attP sites varying in size from 30 bp to 50 bp, were co-transfected with PE components, a PhiC31 P3 helper expression plasmid, and a 6.6 kb donor plasmid containing the full size PhiC31 attP or attB site respectively. PE and integration efficiencies were measured by ddPCR. n=3. Data are shown as mean + s.d.

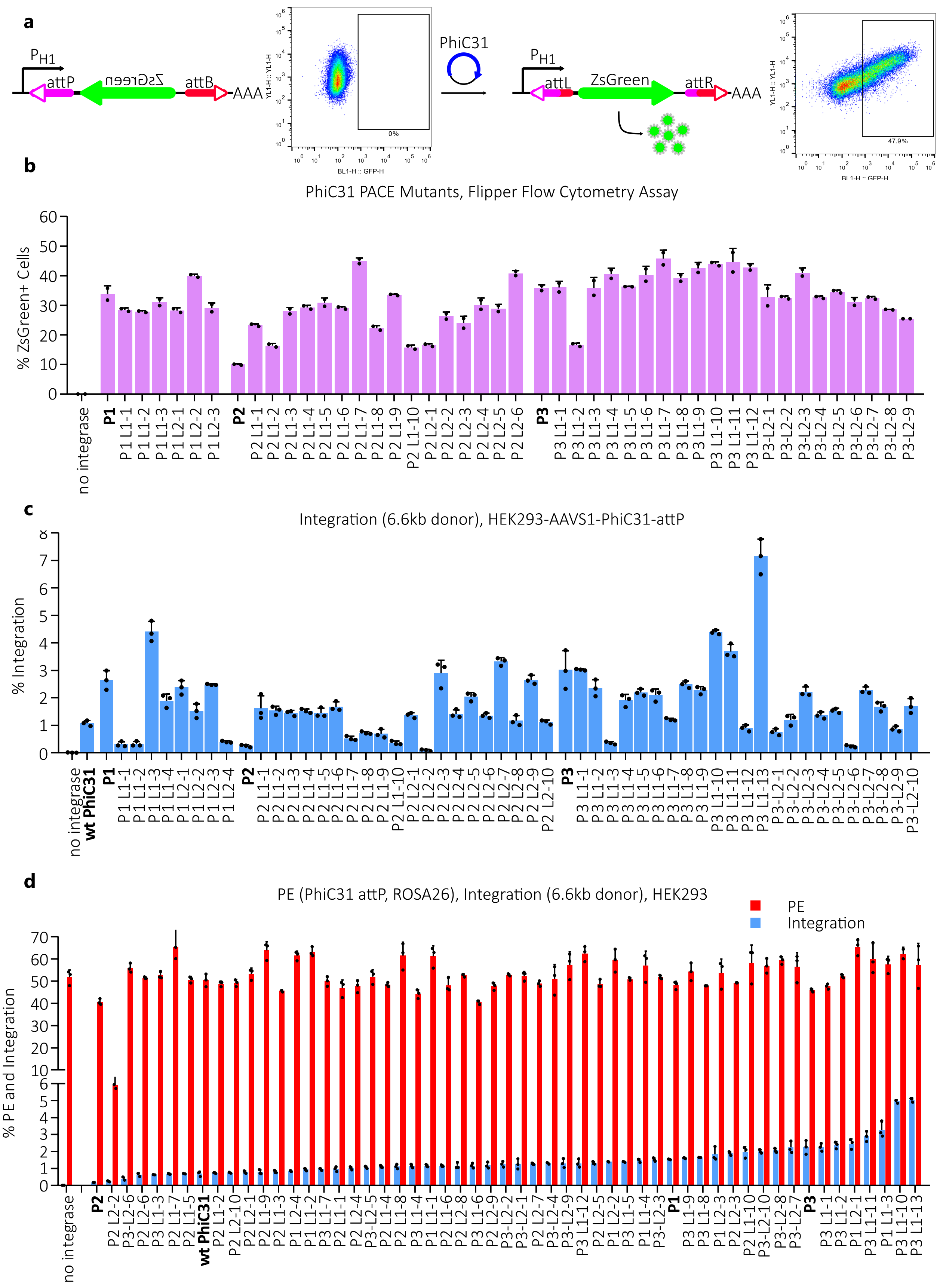

a
PH1
attP
ZsGreen
attB
AAA
PhiC31
attL
attR
0%
47.9%
BL1-H :: GFP-H
YL1-H :: YL1-H
b
PhiC31 PACE Mutants, Flipper Flow Cytometry Assay
% ZsGreen+ Cells
no integrase
P1
P2
P3
c
Integration (6.6kb donor), HEK293-AAVS1-PhiC31-attP
% Integration
no integrase
wt PhiC31
d
PE (PhiC31 attP, ROSA26), Integration (6.6kb donor), HEK293
PE
Integration
% PE and Integration

**Supplementary Fig. S2. Efficiency of IntePACE evolved PhiC31 integrase mutants. a**, Schematic and example from the flipper flow cytometry assay. Integrase clones isolated from IntePACE were cloned into mammalian expression plasmids (helper plasmids). HEK293 cells were co-transfected with evolved integrase helper plasmids and a flipper plasmid containing a promoter oriented away from a ZsGreen gene that was flanked by PhiC31 attP and attB sites. Successful recombination flips the ZsGreen gene to align it with the promoter. Resulting expression is measured by flow cytometry and reported as percent ZsGreen positive cells. **b**, Activity of evolved PhiC31 integrase variants assayed by the flipper flow cytometry assay. Helper plasmid clone names containing starting variant P1, P2, or P3, lagoon number L1 or L2, and clone number are listed on the x-axis. Unevolved controls shown in bold. n = 2. **c**, Evolved integrase helper plasmids were co-transfected with a 6.6 kb donor plasmid into a clonal HEK293 cell line containing a pre-inserted attP site at AAVS1. Integration efficiency was measured with ddPCR. n = 3. **d**, PE insertion of the att site and integration efficiency of the donor plasmid at ROSA26 with IntePACE evolved PhiC31 integrases. n = 3. Data are shown as mean + s.d.

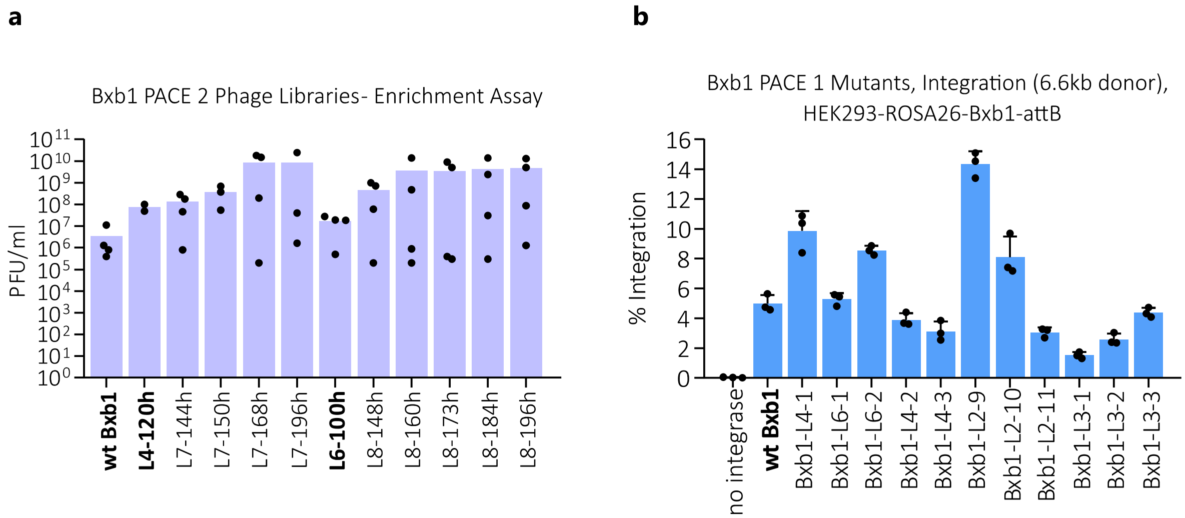


**Supplementary Fig. S3. Efficiency of IntePACE evolved Bxb1 integrase mutants. a**, Pooled phage libraries from IntePACE time points were compared by the overnight enrichment assay using a CP containing the promoter ProA. Starting material from PACE 1 used to inoculate PACE 2 is shown in bold. n = 4. **b**, Integration efficiency of PACE 1 evolved Bxb1 integrase mutants in a clonal HEK293 cell line with a preinstalled ROSA26 attB site measured by ddPCR. Mutations are listed in Supplementary Table S8. n = 4. Data are shown as mean + s.d.

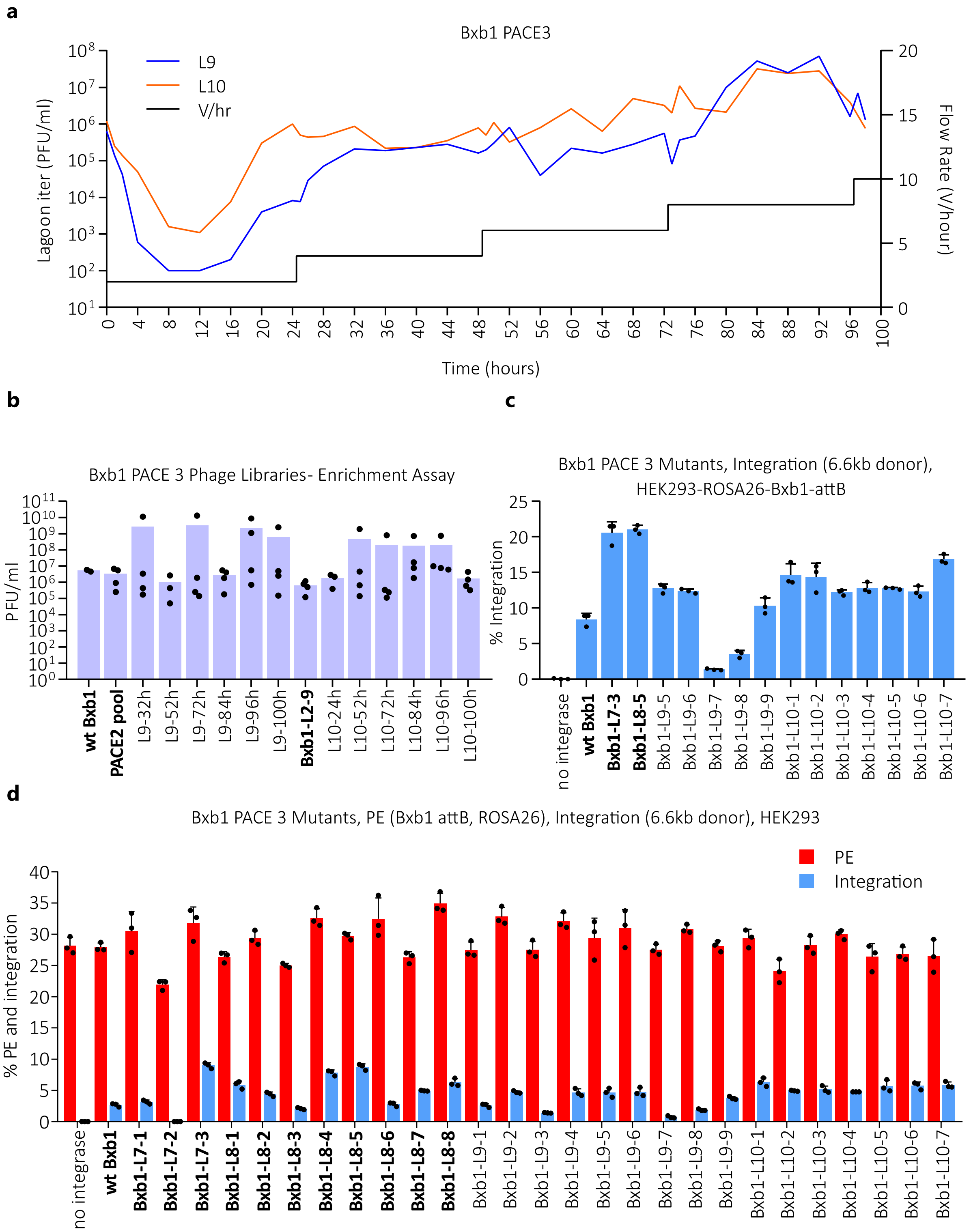
a
Bxb1 PACE3
L9
L10
V/hr
Lagoon iter (PFU/ml)
Flow Rate (V/hour)
Time (hours)
b
Bxb1 PACE 3 Phage Libraries- Enrichment Assay
PFU/ml
wt Bxb1
PACE2 pool
L9-32h
L9-52h
L9-72h
L9-84h
L9-96h
L9-100h
Bxb1-L2-9
L10-24h
L10-52h
L10-72h
L10-84h
L10-96h
L10-100h
c
Bxb1 PACE 3 Mutants, Integration (6.6kb donor),
HEK293-ROSA26-Bxb1-attB
% Integration
no integrase
wt Bxb1
Bxb1-L7-3
Bxb1-L8-5
Bxb1-L9-5
Bxb1-L9-6
Bxb1-L9-7
Bxb1-L9-8
Bxb1-L9-9
Bxb1-L10-1
Bxb1-L10-2
Bxb1-L10-3
Bxb1-L10-4
Bxb1-L10-5
Bxb1-L10-6
Bxb1-L10-7
d
Bxb1 PACE 3 Mutants, PE (Bxb1 attB, ROSA26), Integration (6.6kb donor), HEK293
PE
Integration
% PE and integration
no integrase
wt Bxb1
Bxb1-L7-1
Bxb1-L7-2
Bxb1-L7-3
Bxb1-L8-1
Bxb1-L8-2
Bxb1-L8-3
Bxb1-L8-4
Bxb1-L8-5
Bxb1-L8-6
Bxb1-L8-7
Bxb1-L8-8
Bxb1-L9-1
Bxb1-L9-2
Bxb1-L9-3
Bxb1-L9-4
Bxb1-L9-5
Bxb1-L9-6
Bxb1-L9-7
Bxb1-L9-8
Bxb1-L9-9
Bxb1-L10-1
Bxb1-L10-2
Bxb1-L10-3
Bxb1-L10-4
Bxb1-L10-5
Bxb1-L10-6
Bxb1-L10-7

**Supplementary Fig. S4. Additional rounds of IntePACE did not result in improved Bxb1 integrase variants. a**, For the third Bxb1 experiment, PACE 3, two lagoons (L9 and L10) were run for 96 hours with a flow rate increase every 24 hours (black line). L9 was seeded with selected timepoint pools from PACE 2 (L7-44h, L7-72h, L8-24h, L8-60h, and L8-72h) and L10 was seeded with the PACE 2 clone Bxb1 L2-9 (in bold). Lagoon phage titers were measured by activity-independent plaque assay at least every 4 hours. Phage titers are shown in log scale on the y-axis. **b**, IntePACE time point phage libraries were compared by the overnight enrichment assay. Starting material for PACE is shown in bold. n = 4. **c**, Integration efficiency of PACE 3 mutants in a clonal HEK293 cell line with a preinstalled ROSA26 attB site. Select clonal Bxb1 mutants isolated from PACE 2 lagoons L7 and L8 were compared (shown in bold). n = 3. **d**, PE and integration efficiency at ROSA26 by PACE 3 evolved Bxb1 mutants in HEK293 cells. PACE 2 mutants from lagoons L7 and L8 (shown in bold) were compared to PACE 3 mutants from lagoons L9 and L10. n = 3 (for additional details see Supplementary Note S5). Data are shown as mean + s.d.

**a**

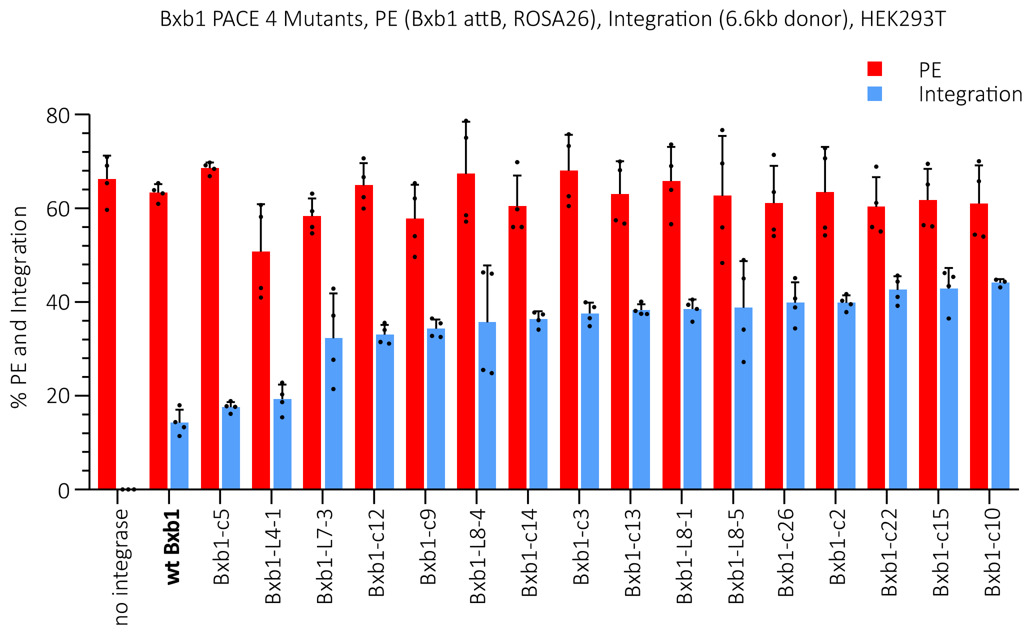

Bxb1 PACE 4 Mutants, PE (Bxb1 attB, ROSA26), Integration (6.6kb donor), HEK293T
PE
Integration
% PE and Integration
80
60
40
20
0
no integrase
wt Bxb1
Bxb1-c5
Bxb1-L4-1
Bxb1-L7-3
Bxb1-c12
Bxb1-c9
Bxb1-L8-4
Bxb1-c14
Bxb1-c3
Bxb1-c13
Bxb1-L8-1
Bxb1-L8-5
Bxb1-c26
Bxb1-c2
Bxb1-c22
Bxb1-c15
Bxb1-c10

**b**

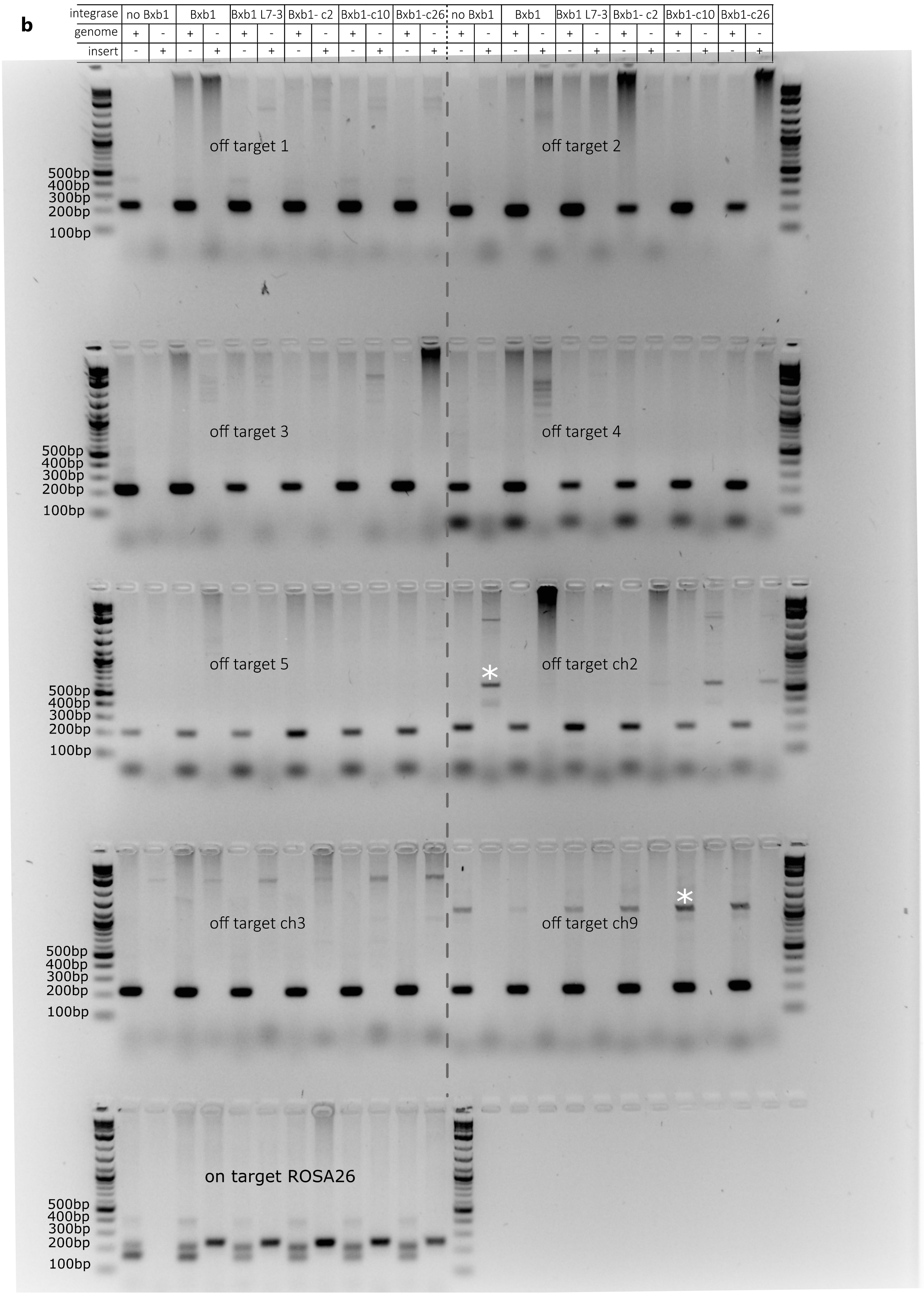
integrase
no Bxb1
Bxb1
Bxb1 L7-3
Bxb1- c2
Bxb1-c10
Bxb1-c26
genome
insert
off target 1
off target 2
off target 3
off target 4
off target 5
off target ch2
off target ch3
off target ch9
on target ROSA26
500bp
400bp
300bp
200bp
100bp
*

**Supplementary Fig. S5. Integration efficiency of Bxb1 variants in HEK293T cells and off-target analysis**. HEK293T cells were transfected with PE5max, epegRNA, integrase helper, and donor plasmids and lysed after three days. **a**, PE insertion of the att site and integration efficiency of the donor plasmid at ROSA26 of top integrase variants from IntePACE and combination mutants in HEK293T cells. n = 4. **b**, Genomic PCR was used to detect insertions at predicted Bxb1 pseudo sites from Anzalone et al.(3) (off target 1-5) and Yarnall et al. (4) (off target ch2, ch3, ch9). Six samples were chosen from the evolved integrase experiment depicted in (**a**). For each site, both a control primer pair and target primer pair were used. The control primers flanked the site in the genome. The target primer pair contained one primer in the genome and one in the donor plasmid. Control on-target primer pairs were used to detect insertion at ROSA26. * sequenced non-specific products. n = 2 (one gel shown).

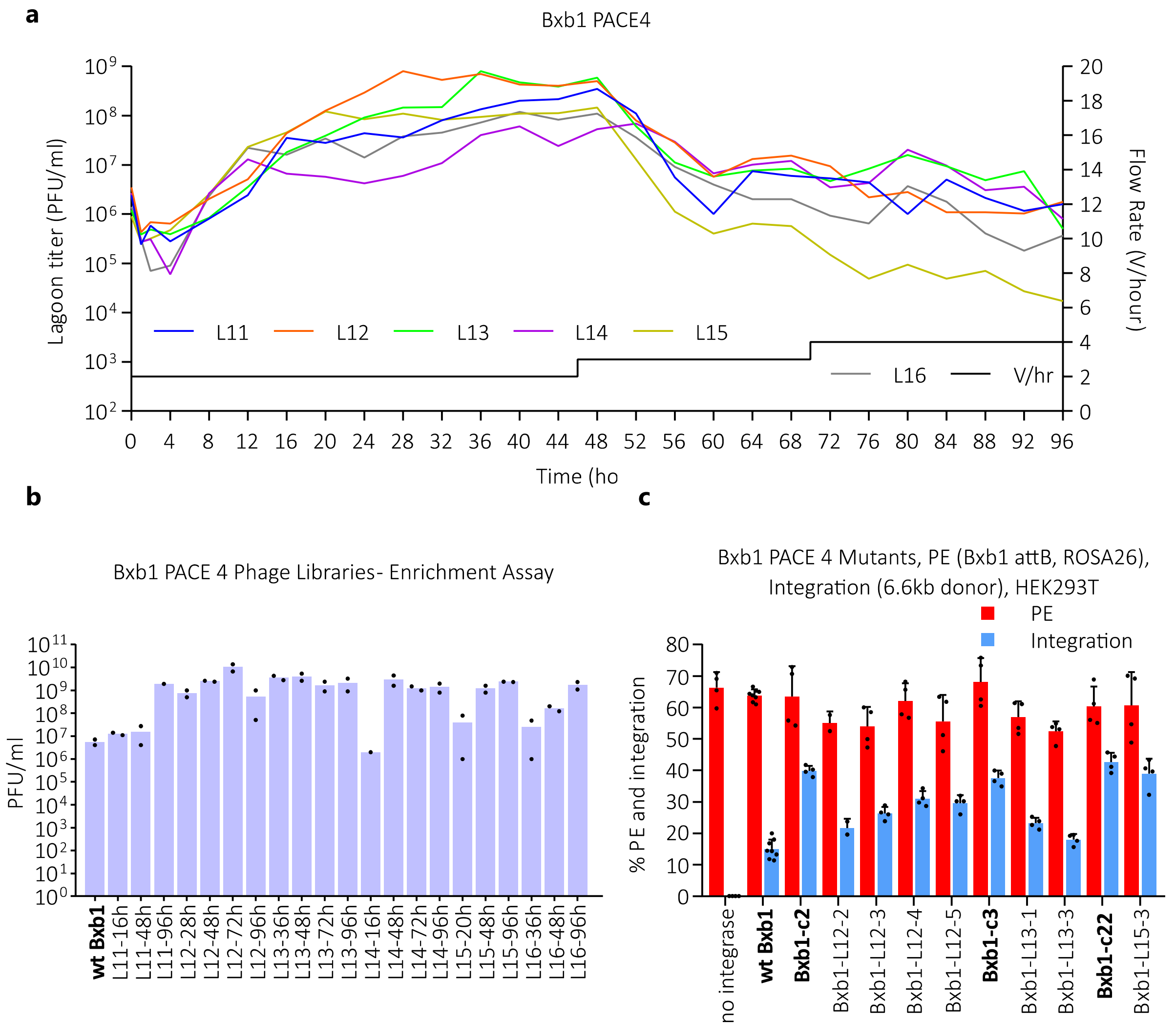


**Supplementary Fig. S6. Additional rounds of IntePACE did not improve Bxb1 combination mutants.** **a**, For the fourth Bxb1 experiment, PACE 4, six lagoons were seeded from wildtype or hyperactive combination mutants (see Supplementary Table S2) and run for 96 hours using AP and CP with a low copy origin of replication (R6K gamma) for increased stringency. By reducing the number of AP for available recombination and gIII expression, only highly active variants are expected to generate enough pIII to sustain progeny in the system. To further increase stringency, the flow rate was increased at 48 and 72 hours. Lagoon phage titers were measured by activity-independent plaque assay at least every 4 hours. **b**, IntePACE time point phage libraries were compared by the overnight enrichment assay. n = 2. **c**, A subset of PACE 4 evolved clones was selected from time point phage libraries and PE and integration efficiency at ROSA26 was assayed in HEK293T cells. Starting material Bxb1 combination mutants Bxb1-c2, Bxb1-c3, and Bxb1-c22 are shown in bold. n = 3. Data are shown as mean + s.d.

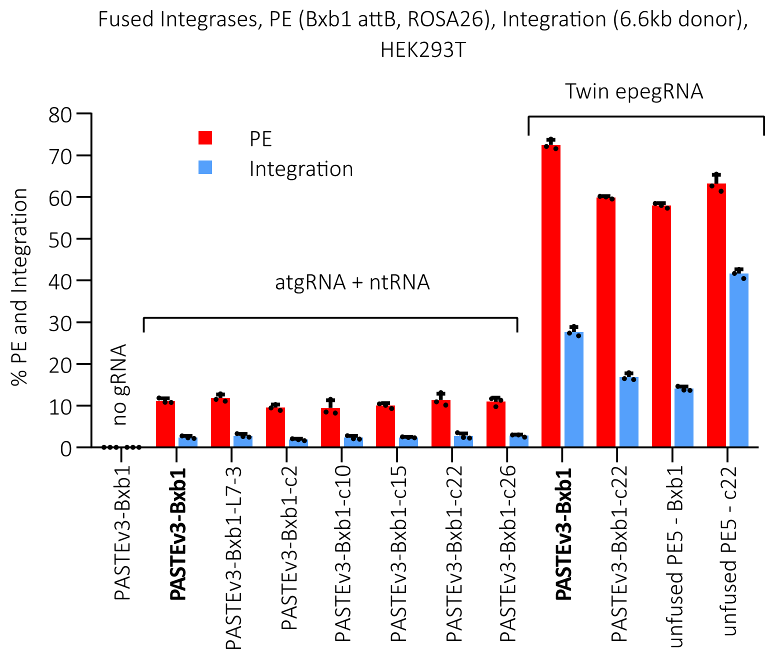


**Supplementary Fig. S7. Comparison of traditional PE PASTE, twin PE PASTE, and untethered PE integrase insertion efficiencies with and without hyperactive mutations.** PASTE uses a wildtype Bxb1 integrase tethered to a Cas9 nickase and reverse transcriptase (PASTEv3-Bxb1). During traditional PE PASTE described by Yarnall et al. (4), a single pegRNA plasmid containing the attB site (called the atgRNA) is co-transfected with a nicking guide RNA (ngRNA) to insert the Bxb1 attB site. Hyperactive mutations recovered from IntePACE were cloned into PASTEv3-Bxb1 and PE and integration efficiency at ROSA26 was measured by ddPCR. For comparison, PASTE was modified to use twin PE by co-transfecting PASTEv3-Bxb1 with twin epegRNA. Both wildtype Bxb1 and IntePACE derived Bxb1-c22 were compared. Twin PE with untethered wildtype Bxb1, described by Anzalone et al. (3), as well as the Bxb1-c22 mutant, described here, were compared for PE and integration efficiency using the same spacers designed to target ROSA26. n = 3. Data are shown as mean + s.d.

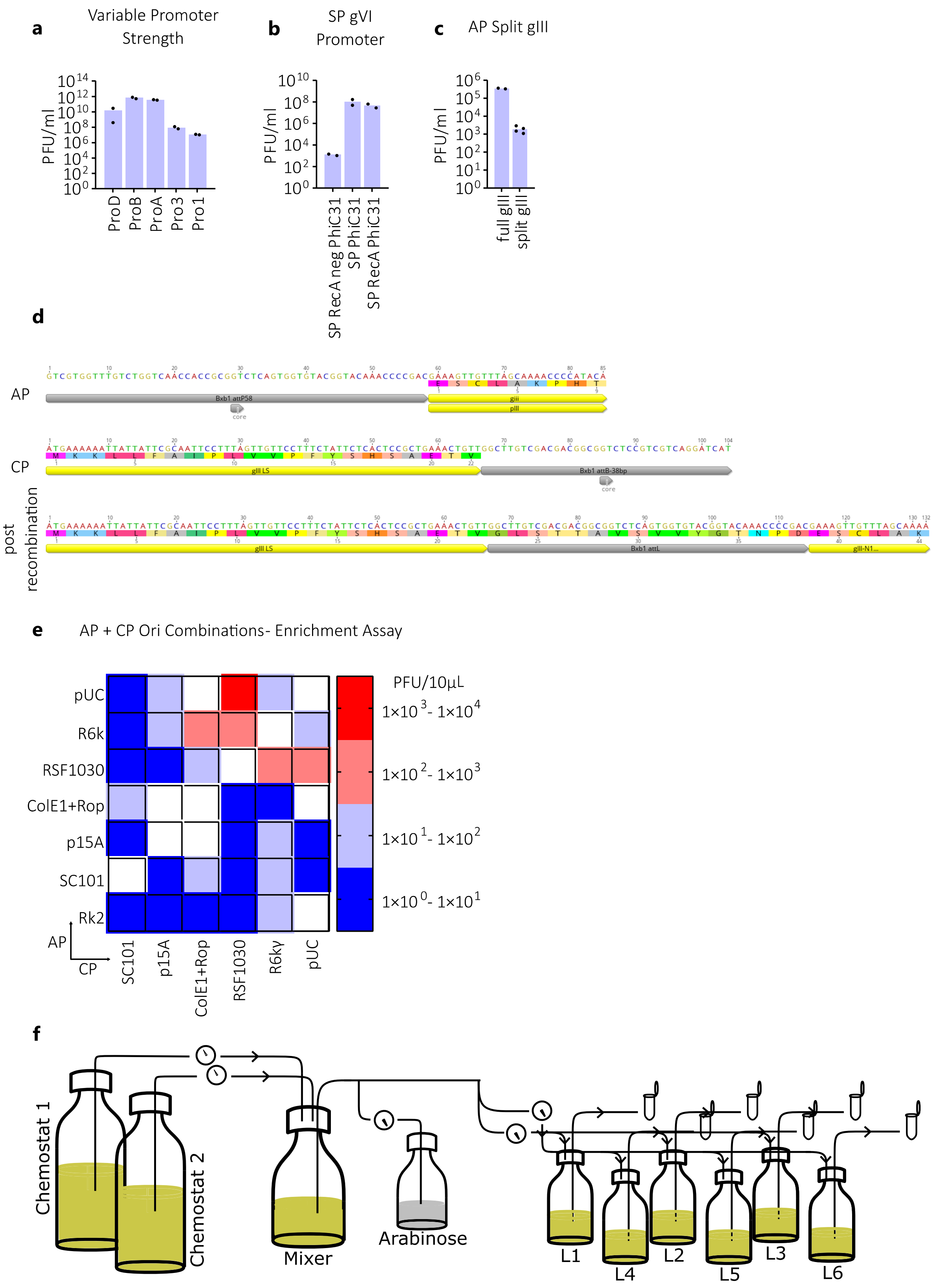
a
Variable Promoter Strength
PFU/ml
ProD
ProB
ProA
Pro3
Pro1
b
SP gVI Promoter
SP RecA neg PhiC31
SP PhiC31
SP RecA PhiC31
c
AP Split gIII
full gIII
split gIII
d
AP
Bxb1 attP58
core
gIII
pIII
CP
gIII LS
Bxb1 attB-38bp
post recombination
Bxb1 attL
gIII-N1...
e
AP + CP Ori Combinations- Enrichment Assay
pUC
R6k
RSF1030
ColE1+Rop
p15A
SC101
Rk2
AP
CP
SC101
p15A
ColE1+Rop
RSF1030
R6kγ
pUC
PFU/10μL
1x10^3- 1x10^4
1x10^2- 1x10^3
1x10^1- 1x10^2
1x10^0- 1x10^1
f
Chemostat 1
Chemostat 2
Mixer
Arabinose
L1
L4
L2
L5
L3
L6

**Supplementary Fig. S8. IntePACE optimizations. a**, Promoter strength for gIII was used to tune stringency. The overnight enrichment assay was used to assay different strength promoters on the CP of the Bxb1 IntePACE system. n = 2. Phage titers for graphs (**a-c)** are shown in log scale. **b**, Traditional PACE typically uses the gVI promoter to drive the protein of interest found in the SP. Because the gVI promoter sequence shares homology with the AP, undesired recombination between these sequences can occur resulting in recombinant wildtype phage that can outcompete other phage. To overcome this issue, a modified SP was used in which the gVI promoter was replaced with the bacterial RecA promoter. Phage expressing PhiC31 integrase under control of the gVI (SP PhiC31) or RecA promoter (SP RecA PhiC31) were used in an overnight enrichment assay to compare the RecA promoter to the gVI promoter's activity. SP RecA neg PhiC31 encoded a catalytically inactive integrase. n = 2. **c**, Although a full-size gIII lacks a promoter on the AP, mis-expression could occur, leading to leaky gIII and undesired propagation of phage without increased recombination activity. By splitting gIII, a recombination event between two plasmids is required to generate full-size gIII. To assay if a split gIII reduces leaking, a split gIII system was tested against a full gIII system using a catalytically inactive PhiC31 integrase in an overnight enrichment assay. n =2. **d**, Sequence of pre- and post-recombination of the AP and CP. The resulting attL site is encoded in-frame between the leader sequence (LS) and the rest of gIII. **e**, During IntePACE, stringency can be increased by lowering the copy number of the AP and CP. Reducing the copy number of available plasmids for recombination and subsequent gIII expression places a higher demand for activity of the evolving integrase. The origin of replication in the IntePACE AP and CP were optimized in the PhiC31 integrase system. Host cells containing all compatible combinations were infected with phage containing PhiC31 integrase. The heat map represents the total number of phage in 10 µl of overnight culture media. Incompatible origins of replication are shown in white. Combinations resulting in high phage titer (red) are suited for low stringency experiments and combinations with low phage titer (blue) are more stringent. The origins of replication used in IntePACE experiments are listed in Supplementary Table S2. n = 2. **f**, Schematic of the PACE apparatus. Host cells flow from two chemostats, are combined in the mixer, and are distributed to six lagoons containing arabinose media and phage. A second chemostat and a mixer were added to a typical PACE setup to accommodate flow rates for six lagoons (for additional details see Supplementary Note 4).

**Supplementary Table S4. Full attB and attP sites for PhiC31, Bxb1, and PaO1 integrases.**

| Phage Integrase | attB | attP | ref |
|---|---|---|---|
| PhiC31 | ctcgaagccgcggtgcgggtgccagggc gtgcccttgggctccccgggcgcgtactccacctcacccatc | cgggagtagtgccccaactggggtaacctttgagttctctcagttgggggcgtagggtcg | Groth 2000 (5) |
| Bxb1 | tcggccggcttgtcgacgacggc ggtctccgtcgtcaggatcatccgggc | gtggtttgtctggtcaaccaccgcg gtctcagtggtgtacggtacaaaccca | Ghosh 2003 (6) |
| PaO1 | ccggtttcccttcgcacccgcaccgcggcttcgaga Ccgtgacctacatgctcgaagggcgtatgcgcc acgaagaccacctcggcaatcgcggcctgctcaag | gtaacgctcttcgagaaagcagattctcatatccatcttgagt cttctttctcgcaagacaacacgaaatagacacagtctcttccctagctgtacactgagcc | Durrant 2023 (7) |

Full attB and attP sites for PhiC31, Bxb1, and PaO1 integrases. Minimal functional att site sequences used in this study are colored red.

### Supplementary Table S5. Un-edited and expected edited amplicon sequences for genome editing analysis using CRISPResso2.

| TARGET | SEQUENCE |
|---|---|
| AAV 1a+1b (Un-edited) | CACTAAGGCAATTGGGGTGCAGGAATGGGGGCAGGGTACCAGCCTCACCAAGTGGTTGATAAACCCACGTGGGGTACCCTAAGAACTTGGGAACAGCCACAGCAGGGGGGCGATGCTTGGGGACCTGCCTGGAGAAGGATGCAGGACGAGAAACACAGCCCCAGGTGGAGAAACTGGCCGGGAATCAAGAGTCACCCAGAGACAGTGACCAACCATCCCTGTTTTCCTAGGACTGAGGGTTTCAGTGCTAAAACTAGGC |
| AAV 1a+1b attB35 | CACTAAGGCAATTGGGGTGCAGGAATGGGGGCAGGGTACCAGCCTCACCAAGTGGTTGATAAACCCGTGCCAGGGCGTGCCCTTGGGCTCCCCGGGCGCGTGTGGAGAAACTGGCCGGGAATCAAGAGTCACCCAGAGACAGTGACCAACCATCCCTGTTTTCCTAGGACTGAGGGTTTCAGTGCTAAAACTAGGC |
| AAV 1a+1b attP41 | CACTAAGGCAATTGGGGTGCAGGAATGGGGGCAGGGTACCAGCCTCACCAAGTGGTTGATAAACCCGCCCCCAACTGAGAGAACTCAAAGGTTACCCCAGTTGGGGCGTGGAGAAACTGGCCGGGAATCAAGAGTCACCCAGAGACAGTGACCAACCATCCCTGTTTTCCTAGGACTGAGGGTTTCAGTGCTAAAACTAGGC |
| AAV 2a+2b (Un-edited) | GCCAGGACGGGGCTGGCTACTGGCCTTATCTCACAGGTAAAACTGACGCACGGAGGAACAATATAAATTGGGGACTAGAAAGGTGAAGAGCCAAAGTTAGAACTCAGGACCAACTTATTCTGATTTTGTTTTTCCAAACTGCTTCTCCTCTTGGGAAGTGTAAGGAAGCTGCAGCACCAGGATCAGTGAAACGCACCAGACAGCCGCGTCAGAGCAGCTCAGGTTCTGGGAGAGGGTAGCGCAGGGTGGCC |
| AAV 2a+2b attB35 | GCCAGGACGGGGCTGGCTACTGGCCTTATCTCACAGGTAAAACTGACGCAGTGCCAGGGCGTGCCCTTGGGCTCCCCGGGCGCGTGGGAAGTGTAAGGAAGCTGCAGCACCAGGATCAGTGAAACGCACCAGACAGCCGCGTCAGAGCAGCTCAGGTTCTGGGAGAGGGTAGCGCAGGGTGGCC |
| AAV 2a+2b attP41 | GCCAGGACGGGGCTGGCTACTGGCCTTATCTCACAGGTAAAACTGACGCAGCCCCCAACTGAGAGAACTCAAAGGTTACCCCAGTTGGGGCGGGAAGTGTAAGGAAGCTGCAGCACCAGGATCAGTGAAACGCACCAGACAGCCGCGTCAGAGCAGCTCAGGTTCTGGGAGAGGGTAGCGCAGGGTGGCC |
| AAV 3a+3b (Un-edited) | AACTGCTTCTCCTCTTGGGAAGTGTAAGGAAGCTGCAGCACCAGGATCAGTGAAACGCACCAGACAGCCGCGTCAGAGCAGCTCAGGTTCTGGGAGAGGGTAGCGCAGGGTGGCCACTGAGAACCGGGCAGGTCACGCATCCCCCCCTTCCCTCCCACCCCCTGCCAAGCTCTCCCTCCCAGGATCCTCTCTGGCTCCATCGTAAGCAAACCTTAGAGGTTCTGGCAAGGAGAGAGATGGCTCCAGGAAATGGGGG |
| AAV 3a+3b (3) attP | AACTGCTTCTCCTCTTGGGAAGTGTAAGGAAGCTGCAGCACCAGGATCAGTGAAACGCACCAGGTTTGTCTGGTCAACCACCGCGGTCTCAGTGGTGTACGGTACAAACCTCCTCTCTGGCTCCATCGTAAGCAAACCTTAGAGGTTCTGGCAAGGAGAGAGATGGCTCCAGGAAATGGGGG |
| AAV 3a+3b attB35 | AACTGCTTCTCCTCTTGGGAAGTGTAAGGAAGCTGCAGCACCAGGATCAGTGAAACGCACCGTGCCAGGGCGTGCCCTTGGGCTCCCCGGGCGCGTTCCTCTCTGGCTCCATCGTAAGCAAACCTTAGAGGTTCTGGCAAGGAGAGAGATGGCTCCAGGAAATGGGGG |
| AAV 3a+3b attP41 | AACTGCTTCTCCTCTTGGGAAGTGTAAGGAAGCTGCAGCACCAGGATCAGTGAAACGCACCGCCCCCAACTGAGAGAACTCAAAGGTTACCCCAGTTGGGGCTCCTCTCTGGCTCCATCGTAAGCAAACCTTAGAGGTTCTGGCAAGGAGAGAGATGGCTCCAGGAAATGGGGG |
| AAV 4a+3b (Un-edited) | AACTGCTTCTCCTCTTGGGAAGTGTAAGGAAGCTGCAGCACCAGGATCAGTGAAACGCACCAGACAGCCGCGTCAGAGCAGCTCAGGTTCTGGGAGAGGGTAGCGCAGGGTGGCCACTGAGAACCGGGCAGGTCACGCATCCCCCCCTTCCCTCCCACCCCCTGCCAAGCTCTCCCTCCCAGGATCCTCTCTGGCTCCATCGTAAGCAAACCTTAGAGGTTCTGGCAAGGAGAGAGATGGCTCCAGGAAATGGGGG |
| AAV 4a+3b (3) attP | AACTGCTTCTCCTCTTGGGAAGTGTAAGGAAGCTGCAGCACCAGGATCAGTGAAACGCACCAGACAGCCGCGTCAGAGCAGCTCAGGTTCTGGGAGGTTTGTCTGGTCAACCACCGCGGTCTCAGTGGTGTACGGTACAAACCTCCTCTCTGGCTCCATCGTAAGCAAACCTTAGAGGTTCTGGCAAGGAGAGAGATGGCTCCAGGAAATGGGGG |
| AAV 4a+3b attB35 | AACTGCTTCTCCTCTTGGGAAGTGTAAGGAAGCTGCAGCACCAGGATCAGTGAAACGCACCAGACAGCCGCGTCAGAGCAGCTCAGGTTCTGGGGTGCCAGGGCGTGCCCTTGGGCTCCCCGGGCGCGTTCCTCTCTGGCTCCATCGTAAGCAAACCTTAGAGGTTCTGGCAAGGAGAGAGATGGCTCCAGGAAATGGGGG |
| AAV 4a+3b attP41 | AACTGCTTCTCCTCTTGGGAAGTGTAAGGAAGCTGCAGCACCAGGATCAGTGAAACGCACCAGACAGCCGCGTCAGAGCAGCTCAGGTTCTGGGGCCCCCAACTGAGAGAACTCAAAGGTTACCCCAGTTGGGGCTCCTCTCTGGCTCCATCGTAAGCAAACCTTAGAGGTTCTGGCAAGGAGAGAGATGGCTCCAGGAAATGGGGG |

| | |
|---|---|
| Rosa 3a+3b (Un-edited) | GGGAGGGAAGCACTGGTTTCTCAAGCAAAAGCTAAAATTTTTCTATTAAGATTTAACCTGATGCTACACTTTGGTGGTGCAGCAAGGGTCTCAAATGGTATAAAACTCAGGTGATCATGCTTTATGTCTGTCTCTAGAAAAATGCTCCAAAAATGATAAGTAGTGATAATCCGCAGTCTCGTTGCATAAAATCAGCCCCAGGTGAATGACTAAGCTCCATTTCCCTACCCCACCCTTATTACAATAACC |
| Rosa 3a+3b attB35 | GGGAGGGAAGCACTGGTTTCTCAAGCAAAAGCTAAAATTTTTCTATTAAGATTTAACCTGATGCTACACTTTGGTGGTGCAGCAAGGGTCTCAAATGGTATAAAAGTGCCAGGGCGTGCCCTTGGGCTCCCCGGGCGCGTGTGAATGACTAAGCTCCATTTCCCTACCCCACCCTTATTACAATAACC |
| Rosa 3a+3b attP41 | GGGAGGGAAGCACTGGTTTCTCAAGCAAAAGCTAAAATTTTTCTATTAAGATTTAACCTGATGCTACACTTTGGTGGTGCAGCAAGGGTCTCAAATGGTATAAAAGCCCCCAACTGAGAGAACTCAAAGGTTACCCCAGTTGGGGCGTGAATGACTAAGCTCCATTTCCCTACCCCACCCTTATTACAATAACC |
| Rosa 4a+4b (Un-edited) | GAGAGGGAGAAAGCTAGTGCTATGTCTGAATACTAGAGGAGCAAGTACAACAAATGGAAAATGGGATCAAGTATGAGTGAGAGTTGCTAAGATGCCTGGTAGGGATGCAAAGGGGTAGAGAGCCTGGGGAGAGAGGGTGAGGGAGGGAAGCACTGGTTTCTCAAGCAAAAGCTAAAATTTTTCTATTAAGATTTAACCTGATGCTACACTTTGGTGGTGCAGCAAGGGTCTCAAATGGTATAAAACTCAGG |
| Rosa 4a+4b attB35 | GAGAGGGAGAAAGCTAGTGCTATGTCTGAATACTAGAGGAGCAAGTACAACAAATGGAAAATGGGATCAAGTATGAGTGAGAGTTGCTAAGATGCCTGGTAGGGATGCGTGCCAGGGCGTGCCCTTGGGCTCCCCGGGCGCGTGCTACACTTTGGTGGTGCAGCAAGGGTCTCAAATGGTATAAAACTCAGG |
| Rosa 4a+4b attP41 | GAGAGGGAGAAAGCTAGTGCTATGTCTGAATACTAGAGGAGCAAGTACAACAAATGGAAAATGGGATCAAGTATGAGTGAGAGTTGCTAAGATGCCTGGTAGGGATGCGCCCCCAACTGAGAGAACTCAAAGGTTACCCCAGTTGGGGCGCTACACTTTGGTGGTGCAGCAAGGGTCTCAAATGGTATAAAACTCAGG |
| Xq22 1a+1b (Un-edited) | AAGGAGAATGACAGACAATAGTATATATGAAATATTCTTATCAAAAGATAAGATCTACTCTAATTAAACCTGTAGGGAAAAAAGGGACAAAGGAGCATGTTAAACAATATCATGAGAATAGAATGAGCAAACTCTAGAATGTGGGAAACTTAGGAAGACACCCAGTTTCTTCAACAAAAAAATTGCAAAAATAAAAAGGAACAAGGAGAACCTATAGATTAAATGAGGCATTAAAGACATATCAACCAAATGTAACCTGGG |
| Xq22 1a+1b attB35 | AAGGAGAATGACAGACAATAGTATATATGAAATATTCTTATCAAAAGATAAGATCTACTCTAATTAAACCTGTGCCAGGGCGTGCCCTTGGGCTCCCCGGGCGCGTGATTAAATGAGGCATTAAAGACATATCAACCAAATGTAACCTGGG |
| Xq22 1a+1b attP41 | AAGGAGAATGACAGACAATAGTATATATGAAATATTCTTATCAAAAGATAAGATCTACTCTAATTAAACCTGCCCCCAACTGAGAGAACTCAAAGGTTACCCCAGTTGGGGCGATTAAATGAGGCATTAAAGACATATCAACCAAATGTAACCTGGG |
| Xq22 1a+2b (Un-edited) | AAGGAGAATGACAGACAATAGTATATATGAAATATTCTTATCAAAAGATAAGATCTACTCTAATTAAACCTGTAGGGAAAAAAGGGACAAAGGAGCATGTTAAACAATATCATGAGAATAGAATGAGCAAACTCTAGAATGTGGGAAACTTAGGAAGACACCCAGTTTCTTCAACAAAAAAATTGCAAAAATAAAAAGGAACAAGGAGAACCTATAGATTAAATGAGGCATTAAAGACATATCAACCAAATGTAACCTGGGGACCATTTAAAATACCTG |
| Xq22 1a+2b attB35 | AAGGAGAATGACAGACAATAGTATATATGAAATATTCTTATCAAAAGATAAGATCTACTCTAATTAAACCTGTGCCAGGGCGTGCCCTTGGGCTCCCCGGGCGCGTGTAACCTGGGGACCATTTAAAATACCTG |
| Xq22 1a+2b attP41 | AAGGAGAATGACAGACAATAGTATATATGAAATATTCTTATCAAAAGATAAGATCTACTCTAATTAAACCTGCCCCCAACTGAGAGAACTCAAAGGTTACCCCAGTTGGGGCGTAACCTGGGGACCATTTAAAATACCTG |
| Xq22 2a+1b (Un-edited) | AAGGAGAATGACAGACAATAGTATATATGAAATATTCTTATCAAAAGATAAGATCTACTCTAATTAAACCTGTAGGGAAAAAAGGGACAAAGGAGCATGTTAAACAATATCATGAGAATAGAATGAGCAAACTCTAGAATGTGGGAAACTTAGGAAGACACCCAGTTTCTTCAACAAAAAAATTGCAAAAATAAAAAGGAACAAGGAGAACCTATAGATTAAATGAGGCATTAAAGACATATCAACCAAATGTAACCTGGGGACCATTTAAAATACCTG |
| Xq22 2a+1b attB35 | AAGGAGAATGACAGACAATAGTATATATGAAATATTCTTATCAAAAGATAAGATCTACTCTAATTAAACCTGTAGGGAAAAAAGGGACAAAGGAGCATGTTAAACAATATCATGAGAATAGAATGAGCAAACTCTAGAGTGCCAGGGCGTGCCCTTGGGCTCCCCGGGCGCGTGATTAAATGAGGCATTAAAGACATATCAACCAAATGTAACCTGGGGACCATTTAAAATACCTG |
| Xq22 2a+1b attP41 | AAGGAGAATGACAGACAATAGTATATATGAAATATTCTTATCAAAAGATAAGATCTACTCTAATTAAACCTGTAGGGAAAAAAGGGACAAAGGAGCATGTTAAACAATATCATGAGAATAGAATGAGCAAACTCTAGAGCCCCCAACTGAGAGAACTCAAAGGTTACCCCAGTTGGGGCGATTAAATGAGGCATTAAAGACATATCAACCAAATGTAACCTGGGGACCATTTAAAATACCTG |

| | |
|---|---|
| Xq22 2a+2b (Un-edited) | AAGGAGAATGACAGACAATAGTATATATGAAATATTCTTATCAAAAGATAAGATCTACTCTAATTAAACCTGTAGGGAAAAAAGGGACAAAGGAGCATGTTAAACAATATCATGAGAATAGAATGAGCAAACTCTAGAATGTGGGAAACTTAGGAAGACACCCAGTTTCTTCAACAAAAAAATTGCAAAAATAAAAAGGAACAAGGAGAACCTATAGATTAAATGAGGCATTAAAGACATATCAACCAAATGTAACCTGGGGACCATTTAAAATACCTG |
| Xq22 2a+2b attB35 | AAGGAGAATGACAGACAATAGTATATATGAAATATTCTTATCAAAAGATAAGATCTACTCTAATTAAACCTGTAGGGAAAAAAGGGACAAAGGAGCATGTTAAACAATATCATGAGAATAGAATGAGCAAACTCTAGAGTGCCAGGGCGTGCCCTTGGGCTCCCCGGGCGCGTGTAACCTGGGGACCATTTAAAATACCTG |
| Xq22 2a+2b attP41 | AAGGAGAATGACAGACAATAGTATATATGAAATATTCTTATCAAAAGATAAGATCTACTCTAATTAAACCTGTAGGGAAAAAAGGGACAAAGGAGCATGTTAAACAATATCATGAGAATAGAATGAGCAAACTCTAGAGCCCCCAACTGAGAGAACTCAAAGGTTACCCCAGTTGGGGCGTAACCTGGGGACCATTTAAAATACCTG |

### Supplementary Table S6. List of Primers

| Primer name | sequence | Figure |
|---|---|---|
| NGS-AAV-1F | TCGTCGGCAGCGTCAGATGTGTATAAGAGACAG GATGACCT aactgcttctcctcttgggaag | Amplicon Seq |
| NGS-AAV-2F | TCGTCGGCAGCGTCAGATGTGTATAAGAGACAG CTGTCTGT aactgcttctcctcttgggaag | Amplicon Seq |
| NGS-AAV-3F | TCGTCGGCAGCGTCAGATGTGTATAAGAGACAG CACACAGT aactgcttctcctcttgggaag | Amplicon Seq |
| NGS-AAV-4F | TCGTCGGCAGCGTCAGATGTGTATAAGAGACAG AACCGGTT aactgcttctcctcttgggaag | Amplicon Seq |
| NGS-AAV-5F | TCGTCGGCAGCGTCAGATGTGTATAAGAGACAG TCTCTCAG aactgcttctcctcttgggaag | Amplicon Seq |
| NGS-AAV-6F | TCGTCGGCAGCGTCAGATGTGTATAAGAGACAG GAGTCACT aactgcttctcctcttgggaag | Amplicon Seq |
| NGS-AAV-25F | TCGTCGGCAGCGTCAGATGTGTATAAGAGACAG AGACTGAG aactgcttctcctcttgggaag | Amplicon Seq |
| NGS-AAV-26F | TCGTCGGCAGCGTCAGATGTGTATAAGAGACAG GGAATACG aactgcttctcctcttgggaag | Amplicon Seq |
| NGS-AAV-27F | TCGTCGGCAGCGTCAGATGTGTATAAGAGACAG AGACCTGT aactgcttctcctcttgggaag | Amplicon Seq |
| NGS-AAV-28F | TCGTCGGCAGCGTCAGATGTGTATAAGAGACAG TAGGCGTT aactgcttctcctcttgggaag | Amplicon Seq |
| NGS-AAV-29F | TCGTCGGCAGCGTCAGATGTGTATAAGAGACAG TGACGACT aactgcttctcctcttgggaag | Amplicon Seq |
| NGS-AAV-7F | TCGTCGGCAGCGTCAGATGTGTATAAGAGACAG AGAGACTG aactgcttctcctcttgggaag | Amplicon Seq |
| NGS-AAV-8F | TCGTCGGCAGCGTCAGATGTGTATAAGAGACAG ATATGCCG aactgcttctcctcttgggaag | Amplicon Seq |
| NGS-AAV-9F | TCGTCGGCAGCGTCAGATGTGTATAAGAGACAG CCTACGAA cactaaggcaattggggtgc | Amplicon Seq |
| NGS-AAV-10F | TCGTCGGCAGCGTCAGATGTGTATAAGAGACAG CGTAGGAA cactaaggcaattggggtgc | Amplicon Seq |
| NGS-AAV-11F | TCGTCGGCAGCGTCAGATGTGTATAAGAGACAG CTCTGACT gccaggacggggctg | Amplicon Seq |
| NGS-AAV-12F | TCGTCGGCAGCGTCAGATGTGTATAAGAGACAG GTCATCAG gccaggacggggctg | Amplicon Seq |
| NGS-Xq22-1F | TCGTCGGCAGCGTCAGATGTGTATAAGAGACAG TCACTCTG aaggagaatgacagacaatagtatatatgaaat | Amplicon Seq |
| NGS-Xq22-2F | TCGTCGGCAGCGTCAGATGTGTATAAGAGACAG CTGACAGT aaggagaatgacagacaatagtatatatgaaat | Amplicon Seq |
| NGS-Xq22-3F | TCGTCGGCAGCGTCAGATGTGTATAAGAGACAG AGTGACAG aaggagaatgacagacaatagtatatatgaaat | Amplicon Seq |
| NGS-Xq22-4F | TCGTCGGCAGCGTCAGATGTGTATAAGAGACAG ACCTACCT aaggagaatgacagacaatagtatatatgaaat | Amplicon Seq |
| NGS-Xq22-5F | TCGTCGGCAGCGTCAGATGTGTATAAGAGACAG CATGTGGT aaggagaatgacagacaatagtatatatgaaat | Amplicon Seq |
| NGS-Xq22-6F | TCGTCGGCAGCGTCAGATGTGTATAAGAGACAG TTCGTAGG aaggagaatgacagacaatagtatatatgaaat | Amplicon Seq |
| NGS-Xq22-7F | TCGTCGGCAGCGTCAGATGTGTATAAGAGACAG GTCTTGAG aaggagaatgacagacaatagtatatatgaaat | Amplicon Seq |
| NGS-Xq22-8F | TCGTCGGCAGCGTCAGATGTGTATAAGAGACAG ACAGAGTG aaggagaatgacagacaatagtatatatgaaat | Amplicon Seq |
| NGS-ROSA-1F | TCGTCGGCAGCGTCAGATGTGTATAAGAGACAG TTCGAACG gtgaatgactaagctccatttccc | Amplicon Seq |
| NGS-ROSA-2F | TCGTCGGCAGCGTCAGATGTGTATAAGAGACAG GATGAGGT gtgaatgactaagctccatttccc | Amplicon Seq |
| NGS-ROSA-3F | TCGTCGGCAGCGTCAGATGTGTATAAGAGACAG CCTACGTT gtgaatgactaagctccatttccc | Amplicon Seq |
| NGS-ROSA-4F | TCGTCGGCAGCGTCAGATGTGTATAAGAGACAG GAGTTGAG gtgaatgactaagctccatttccc | Amplicon Seq |
| NGS-ROSA-1F_2 | TCGTCGGCAGCGTCAGATGTGTATAAGAGACAG TTCGAACG ctccatttccctacccca | Amplicon Seq |
| NGS-ROSA-2F_2 | TCGTCGGCAGCGTCAGATGTGTATAAGAGACAG GATGAGGT ctccatttccctacccca | Amplicon Seq |
| NGS-ROSA-3F_2 | TCGTCGGCAGCGTCAGATGTGTATAAGAGACAG CCTACGTT ctccatttccctacccca | Amplicon Seq |
| NGS-ROSA-4F_2 | TCGTCGGCAGCGTCAGATGTGTATAAGAGACAG GAGTTGAG ctccatttccctacccca | Amplicon Seq |
| NGS-ROSA-5F | TCGTCGGCAGCGTCAGATGTGTATAAGAGACAG GGTTCGAT gggagggaagcactggttt | Amplicon Seq |
| NGS-ROSA-6F | TCGTCGGCAGCGTCAGATGTGTATAAGAGACAG ATCGTTGG gggagggaagcactggttt | Amplicon Seq |
| NGS-ROSA-7F | TCGTCGGCAGCGTCAGATGTGTATAAGAGACAG AGACTCTG gagagggagaaagctagtgctat | Amplicon Seq |
| NGS-ROSA-8F | TCGTCGGCAGCGTCAGATGTGTATAAGAGACAG CAGTACTG gagagggagaaagctagtgctat | Amplicon Seq |
| NGS-AAV-13R | GTCTCGTGGGCTCGGAGATGTGTATAAGAGACAG CATCCAAG cccccatttcctggagcc | Amplicon Seq |
| NGS-AAV-14R | GTCTCGTGGGCTCGGAGATGTGTATAAGAGACAG CGTTGGAT cccccatttcctggagcc | Amplicon Seq |
| NGS-AAV-15R | GTCTCGTGGGCTCGGAGATGTGTATAAGAGACAG GATCTGGT cccccatttcctggagcc | Amplicon Seq |
| NGS-AAV-16R | GTCTCGTGGGCTCGGAGATGTGTATAAGAGACAG GTTCCTTG cccccatttcctggagcc | Amplicon Seq |
| NGS-AAV-17R | GTCTCGTGGGCTCGGAGATGTGTATAAGAGACAG CTAGTGGT cccccatttcctggagcc | Amplicon Seq |
| NGS-AAV-18R | GTCTCGTGGGCTCGGAGATGTGTATAAGAGACAG CAGTCAGT cccccatttcctggagcc | Amplicon Seq |
| NGS-AAV-30R | GTCTCGTGGGCTCGGAGATGTGTATAAGAGACAG GTGTACTG cccccatttcctggagcc | Amplicon Seq |
| NGS-AAV-31R | GTCTCGTGGGCTCGGAGATGTGTATAAGAGACAG GTTGTCCT cccccatttcctggagcc | Amplicon Seq |
| NGS-AAV-32R | GTCTCGTGGGCTCGGAGATGTGTATAAGAGACAG ACCATGGT cccccatttcctggagcc | Amplicon Seq |
| NGS-AAV-33R | GTCTCGTGGGCTCGGAGATGTGTATAAGAGACAG CACAACTG cccccatttcctggagcc | Amplicon Seq |
| NGS-AAV-34R | GTCTCGTGGGCTCGGAGATGTGTATAAGAGACAG CTCTCTCT cccccatttcctggagcc | Amplicon Seq |
| NGS-AAV-19R | GTCTCGTGGGCTCGGAGATGTGTATAAGAGACAG TAGGAACG cccccatttcctggagcc | Amplicon Seq |
| NGS-AAV-20R | GTCTCGTGGGCTCGGAGATGTGTATAAGAGACAG TCTCCAGT cccccatttcctggagcc | Amplicon Seq |
| NGS-AAV-21R | GTCTCGTGGGCTCGGAGATGTGTATAAGAGACAG AGTCGACT gcctagttttagcactgaaaccc | Amplicon Seq |
| NGS-AAV-22R | GTCTCGTGGGCTCGGAGATGTGTATAAGAGACAG AACGTAGG gcctagttttagcactgaaaccc | Amplicon Seq |
| NGS-AAV-23R | GTCTCGTGGGCTCGGAGATGTGTATAAGAGACAG ACTGGACT ggccaccctgcgctac | Amplicon Seq |
| NGS-AAV-24R | GTCTCGTGGGCTCGGAGATGTGTATAAGAGACAG ACTCACAG ggccaccctgcgctac | Amplicon Seq |
| NGS-Xq22-9R | GTCTCGTGGGCTCGGAGATGTGTATAAGAGACAG CTACGTAG cccaggttacatttggttgatatgtc | Amplicon Seq |
| NGS-Xq22-10R | GTCTCGTGGGCTCGGAGATGTGTATAAGAGACAG TTCCGGTT cccaggttacatttggttgatatgtc | Amplicon Seq |
| NGS-Xq22-11R | GTCTCGTGGGCTCGGAGATGTGTATAAGAGACAG GACATGAG caggtattttaaatggtccccagg | Amplicon Seq |
| NGS-Xq22-12R | GTCTCGTGGGCTCGGAGATGTGTATAAGAGACAG CAACCATG caggtattttaaatggtccccagg | Amplicon Seq |
| NGS-Xq22-13R | GTCTCGTGGGCTCGGAGATGTGTATAAGAGACAG ACACCTCT caggtattttaaatggtccccagg | Amplicon Seq |
| NGS-Xq22-14R | GTCTCGTGGGCTCGGAGATGTGTATAAGAGACAG CTCAACAG caggtattttaaatggtccccagg | Amplicon Seq |
| NGS-Xq22-15R | GTCTCGTGGGCTCGGAGATGTGTATAAGAGACAG ACGTTGGT caggtattttaaatggtccccagg | Amplicon Seq |
| NGS-Xq22-16R | GTCTCGTGGGCTCGGAGATGTGTATAAGAGACAG AACGATGG caggtattttaaatggtccccagg | Amplicon Seq |

| | | |
|---|---|---|
| NGS-ROSA-9R | GTCTCGTGGGCTCGGAGATGTGTATAAGAGACAG ACTGTGTG agaagaggtcagaaagccagtc | Amplicon Seq |
| NGS-ROSA-10R | GTCTCGTGGGCTCGGAGATGTGTATAAGAGACAG TGGTGAAG agaagaggtcagaaagccagtc | Amplicon Seq |
| NGS-ROSA-11R | GTCTCGTGGGCTCGGAGATGTGTATAAGAGACAG CTTCTCCT agaagaggtcagaaagccagtc | Amplicon Seq |
| NGS-ROSA-12R | GTCTCGTGGGCTCGGAGATGTGTATAAGAGACAG GAGTGTCT agaagaggtcagaaagccagtc | Amplicon Seq |
| NGS-R_2OSA-9R_2 | GTCTCGTGGGCTCGGAGATGTGTATAAGAGACAG ACTGTGTG gtcagaaagccagtcgcg | Amplicon Seq |
| NGS-R_2OSA-10R_2 | GTCTCGTGGGCTCGGAGATGTGTATAAGAGACAG TGGTGAAG gtcagaaagccagtcgcg | Amplicon Seq |
| NGS-R_2OSA-11R_2 | GTCTCGTGGGCTCGGAGATGTGTATAAGAGACAG CTTCTCCT gtcagaaagccagtcgcg | Amplicon Seq |
| NGS-R_2OSA-12R_2 | GTCTCGTGGGCTCGGAGATGTGTATAAGAGACAG GAGTGTCT gtcagaaagccagtcgcg | Amplicon Seq |
| NGS-ROSA-13R | GTCTCGTGGGCTCGGAGATGTGTATAAGAGACAG TCTCAGAG ggttattgtaataagggtggggtagg | Amplicon Seq |
| NGS-ROSA-14R | GTCTCGTGGGCTCGGAGATGTGTATAAGAGACAG CAGTTCAG ggttattgtaataagggtggggtagg | Amplicon Seq |
| NGS-ROSA-15R | GTCTCGTGGGCTCGGAGATGTGTATAAGAGACAG AGTGCACT cctgagttttataccatttgagaccc | Amplicon Seq |
| NGS-ROSA-16R | GTCTCGTGGGCTCGGAGATGTGTATAAGAGACAG ACCTAGGT cctgagttttataccatttgagaccc | Amplicon Seq |
| NGS-ROSA-1F | TCGTCGGCAGCGTCAGATGTGTATAAGAGACAG TTCGAACG gtgaatgactaagctccatttccc | Amplicon Seq |
| NGS-ROSA-2F | TCGTCGGCAGCGTCAGATGTGTATAAGAGACAG GATGAGGT gtgaatgactaagctccatttccc | Amplicon Seq |
| NGS-ROSA-3F | TCGTCGGCAGCGTCAGATGTGTATAAGAGACAG CCTACGTT gtgaatgactaagctccatttccc | Amplicon Seq |
| NGS-ROSA-4F | TCGTCGGCAGCGTCAGATGTGTATAAGAGACAG GAGTTGAG gtgaatgactaagctccatttccc | Amplicon Seq |
| NGS-ROSA-1F_2 | TCGTCGGCAGCGTCAGATGTGTATAAGAGACAG TTCGAACG ctccatttccctacccca | Amplicon Seq |
| NGS-ROSA-2F_2 | TCGTCGGCAGCGTCAGATGTGTATAAGAGACAG GATGAGGT ctccatttccctacccca | Amplicon Seq |
| NGS-ROSA-3F_2 | TCGTCGGCAGCGTCAGATGTGTATAAGAGACAG CCTACGTT ctccatttccctacccca | Amplicon Seq |
| NGS-ROSA-4F_2 | TCGTCGGCAGCGTCAGATGTGTATAAGAGACAG GAGTTGAG ctccatttccctacccca | Amplicon Seq |
| NGS-ROSA-9R | GTCTCGTGGGCTCGGAGATGTGTATAAGAGACAG ACTGTGTG agaagaggtcagaaagccagtc | Amplicon Seq |
| NGS-ROSA-10R | GTCTCGTGGGCTCGGAGATGTGTATAAGAGACAG TGGTGAAG agaagaggtcagaaagccagtc | Amplicon Seq |
| NGS-ROSA-11R | GTCTCGTGGGCTCGGAGATGTGTATAAGAGACAG CTTCTCCT agaagaggtcagaaagccagtc | Amplicon Seq |
| NGS-ROSA-12R | GTCTCGTGGGCTCGGAGATGTGTATAAGAGACAG GAGTGTCT agaagaggtcagaaagccagtc | Amplicon Seq |
| NGS-R_2OSA-9R_2 | GTCTCGTGGGCTCGGAGATGTGTATAAGAGACAG ACTGTGTG gtcagaaagccagtcgcg | Amplicon Seq |
| NGS-R_2OSA-10R_2 | GTCTCGTGGGCTCGGAGATGTGTATAAGAGACAG TGGTGAAG gtcagaaagccagtcgcg | Amplicon Seq |
| NGS-R_2OSA-11R_2 | GTCTCGTGGGCTCGGAGATGTGTATAAGAGACAG CTTCTCCT gtcagaaagccagtcgcg | Amplicon Seq |
| NGS-R_2OSA-12R_2 | GTCTCGTGGGCTCGGAGATGTGTATAAGAGACAG GAGTGTCT gtcagaaagccagtcgcg | Amplicon Seq |
| SG Rosa 1a+1b For | gtgaatgactaagctccatttccc | Amplicon Seq |
| SG Rosa 1a+1b rev | agaagaggtcagaaagccagtc | Amplicon Seq |
| SG Rosa 1a+1b For 2 | ctccatttccctacccca | Amplicon Seq |
| SG Rosa 1a+1b rev2 | gtcagaaagccagtcgcg | Amplicon Seq |
| SG Rosa 1a+1b For 3 | ccctaccccacccttattaca | Amplicon Seq |
| SG Rosa 1a+1b rev 3 | gaggtcagaaagccagtcgc | Amplicon Seq |
| ddPCR pCMV Rev | gtttgtccaaactcagcggc | ddPCR |
| ddPCR pCMV OPP Rev | gaaggtacgcctgcaggtac | ddPCR |
| ddPCR pCMV OPP Rev2 | atccagcctccggactctag | ddPCR |
| ddPCR attB35 Rev | ccggggagcccaaggg | ddPCR |
| ddPCR attP41 Rev | gagttctctcagttgggggc | ddPCR |
| ddPCR attP41 Rev2 | ccaactggggtaacctttgagt | ddPCR |
| ddPCR Rosa1a For | tccgcagtctcgttgcataa | ddPCR |
| ddPCR Rosa3a For | acaacaaatggaaaatgggatca | ddPCR |
| ddPCR Rosa3a For2 | agaacgtgaactagggagga | ddPCR |
| ddPCR Xq1a For | agcctctatattctaatccacttgt | ddPCR |
| ddPCR Xq1a For2 | ctgtgcctagcctaagcctc | ddPCR |
| ddPCR Xq1a For3 | tgctggaattacaggcgtga | ddPCR |
| ddPCR AAV3a For | gcttctcctcttgggaagtgtaa | ddPCR |
| ddPCR AAV2a For | aagggagttttccacacgga | ddPCR |
| ddPCR AAV1a For | cactaaggcaattggggtgc | ddPCR |
| ddPCR BxBI attB38 Rev | gtcgacgacggcggtctc | ddPCR |
| ddPCR BxBI attB38 Rev2 | CTCCGTCGTCAGGATCATCC | ddPCR |
| ddPCR BxBI attP48 Rev | gtacaccactgagaccgcg | ddPCR |
| ddPCR Pa01 attB33 Rev | CCGTGACCTACATGCTCGC | ddPCR |
| ddPCR Pa01 attB33 Rev2 | CTCGAAGGGCGTATGCGC | ddPCR |
| ddPCR Rosa1a Probe | ccccaggtgaatgactaagctccatttccctac | ddPCR |
| Opti Rosa1a Probe | ggggagtgagcagctgtaag | ddPCR |
| ddPCR Rosa3a Probe | agtgagagttgctaagatgcctggtagggatgc | ddPCR |
| Opti Rosa3a Probe | tgctgcaccaccaaagtgta | ddPCR |
| ddPCR Xq1a Probe | tagggccctgatatgggcacccaaatgtagctt | ddPCR |
| Opti Xq1a Probe | aacatgctcctttgtccctt | ddPCR |
| ddPCR AAV3a Probe | ctgcagcaccaggatcagtgaaacgcac | ddPCR |
| Opti AAV3a Probe | gttctcagtggccaccctg | ddPCR |
| ddPCR AAV2a Probe | cccctcctcaccacagccctgcca | ddPCR |
| Opti AAV2a Probe | ggctcttcacctttctagtccc | ddPCR |
| ddPCR AAV1a Probe | taccagcctcaccaagtggttgataaacccacg | ddPCR |
| Opti AAV1a Probe | ctcgtcctgcatccttctcc | ddPCR |

| | | |
|---|---|---|
| ACTB For | acactgtgcccatctac | ddPCR |
| ACTB Rev | aatgtcacgcacgatttc | ddPCR |
| RPP30 For | AGATTTGGACCTGCGAGCG | ddPCR |
| RPP30 Rev | GAGCGGCTGTCTCCACAAGT | ddPCR |
| UBE2D2 For | GTACTCTTGTCCATCTGTTCTCTG | ddPCR |
| UBE2D2 Rev2 | GGCCGATATTCAGCCCTTAAAC | ddPCR |
| ACTB Probe | /5HEX/CGGGACCTG/ZEN/ACTGACTACCTCAT/3IABkF | ddPCR |
| RPP30 Probe | HEX-TTCTGACCTGAAGGCTCTGCGCG-BHQ1 | ddPCR |
| UBE2D2 Probe | /5HEX/CCGAGCAAT/ZEN/CTCAGGCACTAAAGGA/3IABkFQ | ddPCR |
| Rosa OnT Rev | ttctagagacagacataaagcatgatca | Off-target |
| DL OT1 fwd | GGAAATAAGTTATCACAATGGGAAAT | Off-target |
| DL OT1 rev | TCGCGATTCTTAAAAGGAGAGG | Off-target |
| DL OT2 fwd | CCATTCATATTTTGAAACAAAAGG | Off-target |
| DL OT2 rev | GCATTGCACTCCTACATACAACA | Off-target |
| DL OT3 fwd | GCTGTGGTTATTCCCAGCTC | Off-target |
| DL OT3 rev | CTGGGAACACTGGACAAAATCC | Off-target |
| DL OT4 fwd | GGAAAGCTTTGACAAGTGGAA | Off-target |
| DL OT4 rev | GCCTACTTGCCCTTCTTCCT | Off-target |
| PASTE ch2 off For | AGGGACCTTTGCCTGTGTGAGTC | Off-target |
| PASTE ch2 off Rev | cactcacacagacagaggcc | Off-target |
| PASTE ch3 off For | CCAGGTGAGAGTCAGGGTAGTGTTCA | Off-target |
| PASTE ch3 off Rev | gcttgcgcggcctacgt | Off-target |
| PASTE ch9 off For | TCAGCTCTGTGCTGAGGCGAA | Off-target |
| PASTE ch9 off Rev | GCACAACCCTGGCTGTCC | Off-target |
| ddPCR pCMV Rev2 | tgaatgcaattgttgttgttaacttgt | Off-target |

### Supplementary Table S7. PhiC31 Integrase Mutants

| List of mutants | N-term addition | Internal Mutations |
|---|---|---|
| WT PhiC31 | | |
| P1 | MEQGVVSG | M1E, D2V, V41I |
| P2 | | D32A, D36A, D44A |
| P3 | MTMITPSAQLTLTKGNKSWSSLVTAASVLEFATVIQGVAG | M1V, D2V, D32A, D36A, V41I, D44A |
| P1 L1-1 | MIQEVVSG | V41I, M278L |
| P1 L1-2 | MIQGVVSG | V41I, G344V |
| P1 L1-3 | MIQGVVSG | V41I, E431V |
| P1 L2-1 | MIQGVVSG | V41I, R43R, P230S, Q355Q, K520K |
| P1 L2-2 | MIQGVVSG | V41I, V117V, H228Y, A255G, A580V |
| P1 L2-3 | MIQGVVSG | V41I, G505V, P587P, D586N |
| P2 L1-1 | | D32A, A199T, P535P |
| P2 L1-2 | | D252G, |
| P2 L1-3 | | D32A, D36A, D44A, S396N, R551R |
| P2 L1-4 | | N406S, E512K, R621L |
| P2 L1-5 | | G344D, A397T, A616V |
| P2 L1-6 | | D32A, D36A, D44A, S55S, E77E, I153I, G344D, S359S, V552V, A585A |
| P2 L1-7 | | D32A, D36A, D44A, K266K, D362G |
| P2 L1-8 | | D32A, D36A, D44A, F231L, G505S, D592G |
| P2 L1-9 | | D32A, D36A, D44A, I103I, T302A, D362G, G399G, E449D, E475D |
| P2 L1-10 | | D32A, D36A, D44A, E77E, I153I, H200H, P274S, G344D, K382K, T536A |
| P2 L2-1 | | A498A |
| P2 L2-2 | | D32A, D36A, D44A, R320R, A333T, G429S, A516T, A604S |
| P2 L2-3 | | D32A, D36A, D44A, A238A, R457R, P535L |
| P2 L2-4 | | E14G, D32A, D36A, D44A, E176D, S396R, K586Q |
| P2 L2-5 | | D32A, D36A, D44A, E176D, A333D |
| P2 L2-6 | | D32A, D36A, D44A, I74I, R96R, E176D, A238A, E378K, D590G |
| P3 L1-1 | MTMITPSAQLTLTKGNKSWSSLVTAASVLEFATVIQGVAG | M1E, D2V, D32A, D36A, V41I, D44A |
| P3 L1-2 | MTMITPSAQLTLTKGNKSWSSLVTAASVLEFATVIQGVAG | M1E, D2V, D32A, D36A, V41I, D44A, S235S, L364M |
| P3 L1-3 | MTMITPSAQLTLTKGNKSWSSLVTAASVLEFATVIQGVAG | M1E, D2V, D32A, D36A, V41I, D44A, G346S, G505G |
| P3 L1-4 | MTMITPSAQLTLTKGNKSWSSLVTAASVLEFATVIQGVAG | M1E, D2V, D32A, D36A, V41I, D44A, D362N, L468L, V603I |
| P3 L1-5 | MTMITPSAQLTLTKGNKSWSSLVTAASVLEFATVIQGVAG | M1E, D2V, D32A, D36A, V41I, D44A, D362N |
| P3 L1-6 | MTMITPSAQLTLTKGNKSWSSLVTAASVLEFATVIQGVAG | M1E, D2V, D36A, V41I, D44A, N188N, W448R, P609S |
| P3 L1-7 | MTMITPSAQLTLTKGNKSWSSLVTAASVLEFATVIQGVAG | M1E, D2V, D32A, D36A, V41I, D44A, D362G |
| P3 L1-8 | MTMITPSAQLTLTKGNKSWSSLVTAASVLEFATVIQGVAG | M1E, D2V, D32A, D36A, V41I, D44A, F231L, A410T |
| P3 L1-9 | MTMITPSAQLTLTKGNKSWSSLVTAASVLEFATVIQGVAG | M1E, D2V, D32A, D36A, V41I, D44A, T262T, D362N, V393V, L436L |
| P3 L1-10 | MTMITPSAQLTLTKSNKSWSSLVTAASVLEFATVIQGVTG | M1E, D2M, D32A, V41I, D44A, G45G, G264G, E331E, L351V, G445G, V563V, A580V, V603V |
| P3 L1-11 | MTMITPSAQLTLTKGNKSWSSLVTAASVLEFATVIQGVAG | M1E, D2V, A24A, D32A, D36A, V41I, D44A, W438R, P609S |
| P3 L1-12 | MTMITPSAQLTLTKGNKSWSSLVTAASVLEFATVIQGVAG | M1E, D2V, A24A, D32A, D36A, V41I, D44A, H240R, A333S, A340V, W448R, T600S, P609S |
| P3-L2-1 | MTMITPSAQLTLTKGNKSWSSLVTAASVLEFATVIQGVAG | M1E, D2V, D32A, D36A, V41I, D44A, R347K, I424I |
| P3-L2-2 | MTMITPSAQLTLTKDNKSWSSLVTAASVLEFATVIQGVAG | M1E, D2V, D32A, D36A, V41I, D44A, V322V, L351Q, D549D |
| P3-L2-3 | MTMITPSAQLTLTKGNKSWSSLVTAASVLEFATVIQGVAG | M1E, D2V, D32A, D36A, V41I, D44A, S107S, D362N |
| P3-L2-4 | MTMITPSAQLTLTKSNKSWSSLVTAASVLEFATVIQGVAG | M1E, D2V, S12N, D32A, D36A, V41I, D44A, E153V, A580V, V603A |
| P3-L2-5 | MTMITPSAQLTLTKGNKSWSSLVTAASVLEFATVIQGVAG | M1E, D2V, D32A, D36A, V41I, D44A, D362N |
| P3-L2-6 | MTMITPSAQLTLTKSNKSWSSLVTAASVLEFATVIQGVAG | M1E, D2V, S18N, D32A, D36A, V41I, D44A, V51M, I103I, I153V, A450D, V603A |
| P3-L2-7 | MTMITPSAQLTLTKSNKSWSSLVTAASVLEFATVIQGVAG | M1E, D2V, D32A, D36A, V41I, D44A, I153V, S269N, A450D, E517E, V603A |
| P3-L2-8 | MTMITPSAQLTLTKGNKSWSSLVTAASVLEFATVIQGVAG | M1E, D2V, D32A, D36A, V41I, D44A, I153V, G259G, L436L, A450D, L501L, V603A |
| P3-L2-9 | MTMITPSAQLTLTKGNKSWSSLVTAASVLEFATVIQGVAG | M1E, D2V, D32A, D36A, V41I, D44A, I153V, L351V, A450D, E452K, V603A |

### Supplementary Table S8. Bxb1 Integrase Mutants

| Bxb1 Mutant | Mutations |
|---|---|
| WT Bxb1 | |
| Bxb1-L4-1 | H111P, E434G |
| Bxb1-L6-1 | T285A |
| Bxb1-L6-2 | G156G, A369P |
| Bxb1-L4-2 | R63K, R88R, A110A, P295P |
| Bxb1-L4-3 | D36A, R63K, P292P, L479I |
| Bxb1-L2-9 | H95Y |
| Bxb1-L2-10 | V5V, S106S, E434G |
| Bxb1-L2-11 | D51D, H189N |
| Bxb1-L3-1 | L164L, H203Y, Q480STOP |
| Bxb1-L3-2 | F67S, V187I, A261T |
| Bxb1-L3-3 | L4I |
| Bxb1-L7-3 | I87L, V187V, A218A, E419E, A425T, E483E |
| Bxb1-L8-1 | R63K, A119S, P295P, E361E, M499T |
| Bxb1-L8-2 | R63K, V122M, P295P |
| Bxb1-L8-3 | R63K, A347E |
| Bxb1-L8-4 | V40I, R63K, V122M, P295P, R319K, S428S |
| Bxb1-L8-5 | R63K, V122M, P295P, R487R |
| Bxb1-L8-7 | L278L, P295P, P332H, A369T, V380I |
| Bxb1-L8-8 | E42K, R63K, D99N, E133E, A145T, K153K, P295P, A369E, R494S |
| Bxb1-L9-1 | V40A, R63K, H89G, Q191Q, P295P, L302L, C307C |
| Bxb1-L9-2 | V40V, R63K, P295P, R362K, G370G, G468G |
| Bxb1-L9-3 | G209G, A288V, A311V, A398S, R416K, T453I, H496N, E42K, G209G |
| Bxb1-L9-4 | A288V, A311V, A398S, R416K, T453I, H496N |
| Bxb1-L9-5 | S18S, R63K, E69E, I87V, P295P, P332H, H334R, A369E |
| Bxb1-L9-6 | S18S, E42K, R63K, E69E, E133E, P295P, P332H, H334R |
| Bxb1-L9-7 | P268P, A288V, P268P, A311V, D359D, T388M, T453I, V466V, H496N |
| Bxb1-L9-8 | A288V, A311V, R319G, A398S, R416K, T453I, H496N, E434G |
| Bxb1-L9-9 | E42K, A288V, A311V, A398S, R416K, T453I, H496N, E434G |
| Bxb1-L10-1 | H95Y, R287R, K313R |
| Bxb1-L10-2 | H95Y, A280T |
| Bxb1-L10-3 | H95Y, V264A |
| Bxb1-L10-4 | H95Y, T254S |
| Bxb1-L10-5 | V46V, H95Y, V179A, R223R |
| Bxb1-L10-6 | H95Y, V264A, E434G |
| Bxb1-L10-7 | H95Y, A62A, V466M |
| Bxb1-L11-1 | A280A, A360T |
| Bxb1-L11-2 | L61F, E434G |
| Bxb1-L11-3 | L174L, S231S, A288T, E434G, T463I |
| Bxb1-L11-4 | V40V, A130A, V283V, A288T, E434G, G468D |
| Bxb1-L11-5 | E229K, K313K, A405A, T453I |
| Bxb1-L11-6 | L90L, E229K, N251K, D359N, A360T, V375V, R494R |
| Bxb1-L11-7 | Q92Q, V179I, R181K, E229K, M239I, A360T, R444L |
| Bxb1-L12-1 | E69A, I87L, N251N, H321P, D355N |
| Bxb1-L12-2 | D51N, I87L, R272Q, G489G |
| Bxb1-L12-3 | I87L, del L488 frameshift |
| Bxb1-L12-4 | I87L, V105A, H321N, S328S, R397R, A411V |
| Bxb1-L12-5 | R79R, I87L, Q484K |
| Bxb1-L12-6 | I87L, I137I, F331S, R409H, del L488 frameshift |
| Bxb1-L13-1 | E69A, H95Y, R461R |
| Bxb1-L13-2 | H95Y |
| Bxb1-L13-3 | H95Y, L282L |
| Bxb1-L14-1 | I87L, A369P, F476F |
| Bxb1-L14-2 | G34D, I87L, T166I, E229K, A369P, T435T, G489G |
| Bxb1-L14-3 | I87L, R88R, R140R, K333K, A369P, A414A, V466M, F476F |
| Bxb1-L15-1 | I87L, A369S, E434G |
| Bxb1-L15-2 | I87L, A369P, A414V, A425A, E434G |
| Bxb1-L15-3 | R85R, I87L, A248T, V306V, A369P, A415S, E434G |
| Bxb1-L16-1 | I87L, P160P, V175V, V187V, A218A, E281E, V353I, E419E, A425T, E483E |
| Bxb1-L16-2 | I87L, Q92H, P160P, V175V, P178P, V187V, A218A, E281E, V353I, E419E, A425T, E483E |
| Bxb1-L16-3 | E24E, I87L, H100N, V187V, A218A, E419E, A425T, E483E |

| | |
|---|---|
| Bxb1-L16-4 | I87L, V187V, A218A, A369E, E419E, A425T, E483E |
| Bxb1-L16-5 | I87L, V187V, A218A, E419E, A425T, E434G, E483E |
| Bxb1-c1 | |
| Bxb1-c2 | I87L |
| Bxb1-c3 | H95Y |
| Bxb1-c4 | V122M |
| Bxb1-c5 | A369P |
| Bxb1-c6 | E434G |
| Bxb1-c7 | I87L, H95Y |
| Bxb1-c8 | I87L, V122M |
| Bxb1-c9 | I87L, A369P |
| Bxb1-c10 | I87L, E434G |
| Bxb1-c11 | H95Y, V122M |
| Bxb1-c12 | H95Y, A369P |
| Bxb1-c13 | H95Y, E434G |
| Bxb1-c14 | V122M, A369P |
| Bxb1-c15 | V122M, E434G |
| Bxb1-c16 | A369P, E434G |
| Bxb1-c17 | I87L, H95Y, V122M |
| Bxb1-c18 | I87L, H95Y, A369P |
| Bxb1-c19 | I87L, H95Y, E434G |
| Bxb1-c20 | I87L, V122M, A369P |
| Bxb1-c21 | I87L, V122M, E434G |
| Bxb1-c22 | I87L, A369P, E434G |
| Bxb1-c23 | H95Y, V122M, A369P |
| Bxb1-c24 | H95Y, V122M, E434G |
| Bxb1-c25 | H95Y, A369P, E434G |
| Bxb1-c26 | V122M, A369P, E434G |
| Bxb1-c27 | I87L, H95Y, V122M, A369P |
| Bxb1-c28 | I87L, H95Y, V122M, E434G |
| Bxb1-c29 | I87L, H95Y, A369P, E434G |
| Bxb1-c30 | I87L, V122M, A369P, E434G |
| Bxb1-c31 | H95Y, V122M, A369P, E434G |
| Bxb1-c32 | I87L, H95Y, V122M, A369P, E434G |
| Bxb1-c33 | L4I, R63K, I87L, H95Y, H111P, V122M, A280T, A369P, E434G, V466M |
| Bxb1-c34 | R63K, V122M |

### Supplementary Table S9. List of promoters.

| Promoter | Relative strength | Sequence |
|---|---|---|
| ProD | 1.000 | tctagagCACAGCTAACACCACGTCGTCCCTATCTGCTGCCCTAGGTCTATGAGTGGTTGCTGGATAACTTTACGGGCA<br>TGCATAAGGCTCGTATAATATATTCAGGGAGACCACAACGGTTTCCCTCTACAAATAATTTTGTTTAACTTTtactaga |
| ProB | 0.119 | tctagagCACAGCTAACACCACGTCGTCCCTATCTGCTGCCCTAGGTCTATGAGTGGTTGCTGGATAACTTTACGGGCA<br>TGCATAAGGCTCGTAATATATATTCAGGGAGACCACAACGGTTTCCCTCTACAAATAATTTTGTTTAACTTTtactaga |
| ProA | 0.030 | tctagagCACAGCTAACACCACGTCGTCCCTATCTGCTGCCCTAGGTCTATGAGTGGTTGCTGGATAACTTTACGGGCA<br>TGCATAAGGCTCGTAGGCTATATTCAGGGAGACCACAACGGTTTCCCTCTACAAATAATTTTGTTTAACTTTtactaga |
| Pro3 | 0.017 | tctagagCACAGCTAACACCACGTCGTCCCTATCTGCTGCCCTAGGTCTATGAGTGGTTGCTGGATAACTTTACGGGCA<br>TGCATAAGGCTCGGAGGATATATTCAGGGAGACCACAACGGTTTCCCTCTACAAATAATTTTGTTTAACTTTtactaga |
| Pro1 | 0.009 | tctagagCACAGCTAACACCACGTCGTCCCTATCTGCTGCCCTAGGTCTATGAGTGGTTGCTGGATAACTTTACGGGCA<br>TGCATAAGGCTCGGTATCTATATTCAGGGAGACCACAACGGTTTCCCTCTACAAATAATTTTGTTTAACTTTtactaga |

Insulated promoters used to titrate gIII expression in PACE described by Davis et. al 2010 (2). The -35 hexamer is underlined and the -10 hexamer is colored red. The relative strengths of the promoters were determined by GFP expression in *E. coli*.

### Supplementary Table S10. List of Plasmids

| Name | Antibiotic | Ori | Figure |
|---|---|---|---|
| AP LinkRec PhiC31 attP pUC | Carbenicillin | pUC | 3a-b, S8a-b |
| AP LinkRec PhiC31 attP R6Kgamma | Carbenicillin | R6K gamma | S8a |
| AP LinkRec PhiC31 attP RSF1030 | Carbenicillin | RSF1030 | S8a |
| AP LinkRec PhiC31 attP p15A | Carbenicillin | p15A | S8a |
| AP LinkRec PhiC31 attP ColE1Rop | Carbenicillin | ColE1 Rop | S8a |
| AP LinkRec PhiC31 attP RK2 | Carbenicillin | RK2 | S8a |
| AP LinkRec PhiC31 attP SC101 | Carbenicillin | SC101 | S8a |
| AP-P rec PhiC31 attP | Carbenicillin | pUC | S8c |
| AP LinkRec Bxb1 attP pUC | Carbenicillin | pUC | 4b, S3a, S4a-b , S8d |
| AP LinkRec Bxb1 attP R6Kgamma | Carbenicillin | R6K gamma | S6a-b |
| CP LinkDon PhiC31 attB ProB | Kanamycin | RSF1030 | 3a |
| CP LinkDon PhiC31 attB ProA | Kanamycin | RSF1030 | 3a-b, S8b |
| CP LinkDon PhiC31 attB Pro3 | Kanamycin | RSF1030 | 3a |
| CP LinkDon PhiC31 attB Pro1 RSF1030 | Kanamycin | RSF1030 | 3a, S8a |
| CP LinkDon PhiC31 attB Pro1 R6Kgamma | Kanamycin | R6K gamma | S8a |
| CP LinkDon PhiC31 attB Pro1 pUC | Kanamycin | pUC | S8A |
| CP LinkDon PhiC31 attB Pro1 p15A | Kanamycin | p15A | S8a |
| CP LinkDon PhiC31 attB Pro1 ColE1Rop | Kanamycin | ColE1 Rop | S8a |
| CP LinkDon PhiC31 attB Pro1 RK2 | Kanamycin | RK2 | S8a |
| CP LinkDon PhiC31 attB Pro1 SC101 | Kanamycin | SC101 | S8a |
| CP don PhiC31 attB ProA | Kanamycin | RSF1030 | S8c |
| CP LinkDon Bxb1 attB ProD | Kanamycin | RSF1030 | 4b, S8d |
| CP LinkDon Bxb1 attB ProB | Kanamycin | RSF1030 | 4b, S8d |
| CP LinkDon Bxb1 attB ProA | Kanamycin | RSF1030 | 4b, S8d, S3a |
| CP LinkDon Bxb1 attB Pro3 | Kanamycin | RSF1030 | 4b, S8d |
| CP LinkDon Bxb1 attB Pro1 | Kanamycin | RSF1030 | 4b, S8d, S4a-b, S6a-b |
| pJC175e (8) | Carbenicillin | SC101 | 3b, S3a, S4b, S6b, S8a-d |
| MP6 (9) | Chloramphenicol | CloDF13 | 3a, 4b, S4a, S6a |
| SP RecA PhiC31 | | M13 | 3a |
| SP RecA PhiC31 P1Int | | M13 | 3a-b |
| SP RecA PhiC31 P2Int | | M13 | 3a-b |
| SP RecA PhiC31 P3Int | | M13 | 3a-b |
| SP RecA neg-Int | | M13 | S8b-c |
| SP RecA PhiC31 032-10 | | M13 | S8b |
| SP PhiC31 032-10 | | M13 | S8b |
| SP RecA Bxb1 | | M13 | 4b, S3a, S4b, S6b, S8d |
| pCMV2 PhiC31 wt | Carbenicillin | pBR322 | 1c-d, 3d-e, 4a, S2b-d |
| pCMV2 PhiC31 P1 | Carbenicillin | pBR322 | 1c-d, 3d-e, S2b-d |
| pCMV2 PhiC31 P2 | Carbenicillin | pBR322 | 1c-d, 3d-e, S2b-d |
| pCMV2 PhiC31 P3 | Carbenicillin | pBR322 | 1c-d, 3d-e, S1c, S3b-d |
| pCMV2 HuOpt Bxb1 | Carbenicillin | pBR322 | 4a, 4c-d, 5a-b, 6a-h, S3b, S4c-d, S5a-b, S6c, S7 |
| pCMV2 HuOpt Pa01 | Carbenicillin | pBR322 | 4a |
| pH1 EukFlip zsGreen PhiC31 Hygro | Carbenicillin | pBR322 | S2b |
| pCMV PhiC31 attB dsRed Hygro | Carbenicillin | pBR322 | 1c-d, 3d-e, 4a, S2b-d |
| pCMV Bxb1 attP dsRed Hygro | Carbenicillin | pBR322 | 4a, 4c-d, 5a-b, 6a-e, S3b, S4c-d, S5a-b, S6c-d, S7 |
| pCMV Pa01 attP dsRed Hygro | Carbenicillin | pBR322 | 4a |
| pCMV-PEmax-P2A-hmlh1dn (PE5max) (10) | Carbenicillin | pBR322 | 1b, 1d, 3e, 4a, 4d, 5b, 6a-h, S1b-c, S2d, S4d, S5a, S6c, S7 |
| pDY1052 PASTEv3 pCMV-SpCas9-XTEN3-RT(L139P)-(GGS)6-BxbInt (4) | Carbenicillin | pBR322 | S7 |
| pU6 atgRNA Rosa4a Bxb1 attB38 | Carbenicillin | pBR322 | S7 |
| pU6 ntRNA Rosa4b | Carbenicillin | pBR322 | S7 |
| pU6 pegRNA GG Acceptor (11) | Carbenicillin | pBR322 | |
| pCDNA3.1-WT-VWF Bxb1 attP HIR Donor | Carbenicillin | pBR322 | 6g-h |

### Supplementary Table S11. epegRNA Sequences

| Name | Spacer | Scaffold | RT Template | PBS | Linker | TevopreQ1 |
|---|---|---|---|---|---|---|
| aav peg spacer1a attB35 | aagtggttgataaacccacg | cr772 | ggggagcccaagggcacgccctggcac | gggtttatcaacc | GGAATGCC | tevopreQ1 |
| aav peg spacer1b attB35 | ccggccagtttctccacctg | cr772 | ggcgtgcccttgggctccccgggcgcgt | gtggagaaactgg | AATTAATG | tevopreQ1 |
| aav peg spacer2a attB35 | caggtaaaactgacgcacgg | cr772 | ggggagcccaagggcacgccctggcac | tgcgtcagtttta | AGGTTCAA | tevopreQ1 |
| aav peg spacer2b attB35 | gcttccttacacttcccaag | cr772 | ggcgtgcccttgggctccccgggcgcgt | gggaagtgtaagg | TCAAATGA | tevopreQ1 |
| aav peg spacer3a attB35 | gatcagtgaaacgcaccaga | cr772 | ggggagcccaagggcacgccctggcac | ggtgcgtttcact | CTATAAGA | tevopreQ1 |
| aav peg spacer3b attB35 | gatggagccagagaggatcc | cr772 | ggcgtgcccttgggctccccgggcgcgt | tcctctctggctc | GTAATAAT | tevopreQ1 |
| aav peg spacer4a attB35 | gcagctcaggttctgggaga | cr772 | ggggagcccaagggcacgccctggcac | cccagaacctgag | AATTAATT | tevopreQ1 |
| aav peg spacer1a attP41 | aagtggttgataaacccacg | cr772 | tggggtaacctttgagttctctcagttgggggc | gggtttatcaacc | TTCAGACC | tevopreQ1 |
| aav peg spacer1b attP41 | ccggccagtttctccacctg | cr772 | actgagagaactcaaaggttaccccagttggggc | gtggagaaactgg | AATTATAA | tevopreQ1 |
| aav peg spacer2a attP41 | caggtaaaactgacgcacgg | cr772 | tggggtaacctttgagttctctcagttgggggc | tgcgtcagtttta | AAGGTCAC | tevopreQ1 |
| aav peg spacer2b attP41 | gcttccttacacttcccaag | cr772 | actgagagaactcaaaggttaccccagttggggc | gggaagtgtaagg | CCTCAATT | tevopreQ1 |
| aav peg spacer3a attP41 | gatcagtgaaacgcaccaga | cr772 | tggggtaacctttgagttctctcagttgggggc | ggtgcgtttcact | TTATAATA | tevopreQ1 |
| aav peg spacer3b attP41 | gatggagccagagaggatcc | cr772 | actgagagaactcaaaggttaccccagttggggc | tcctctctggctc | AACTGAAA | tevopreQ1 |
| aav peg spacer4a attP41 | gcagctcaggttctgggaga | cr772 | tggggtaacctttgagttctctcagttgggggc | cccagaacctgag | ATATAACA | tevopreQ1 |
| xq22 peg spacer1a attB35 | ctactctaattaaacctgta | cr772 | ggggagcccaagggcacgccctggcac | aggtttaattaga | TGCCATAA | tevopreQ1 |
| xq22 peg spacer1b attB35 | taatgcctcatttaatctat | cr772 | ggcgtgcccttgggctccccgggcgcgt | gattaaatgaggc | CCGAAGAA | tevopreQ1 |
| xq22 peg spacer2a attB35 | aatgagcaaactctagaatg | cr772 | ggggagcccaagggcacgccctggcac | tctagagtttgct | GCCCGGAT | tevopreQ1 |
| xq22 peg spacer2b attB35 | aatggtccccaggttacatt | cr772 | ggcgtgcccttgggctccccgggcgcgt | gtaacctggggac | GTCCGACA | tevopreQ1 |
| xq22 peg spacer1a attP41 | ctactctaattaaacctgta | cr772 | tggggtaacctttgagttctctcagttgggggc | aggtttaattaga | TTCCCATC | tevopreQ1 |
| xq22 peg spacer1b attP41 | taatgcctcatttaatctat | cr772 | actgagagaactcaaaggttaccccagttggggc | gattaaatgaggc | CGAAATGC | tevopreQ1 |
| xq22 peg spacer2a attP41 | aatgagcaaactctagaatg | cr772 | tggggtaacctttgagttctctcagttgggggc | tctagagtttgct | AAGGGACA | tevopreQ1 |

| | | | | | | |
|---|---|---|---|---|---|---|
| xq22 peg spacer2b attP41 | aatggtccccaggttacatt | cr772 | actgagagaactcaaaggttaccccagttggggc | gtaacctggggac | AAGAATTA | tevopreQ1 |
| rosa peg spacer1a attB35 | tcgacaccaactctagtccg | cr772 | ggggagcccaagggcacgccctggcac | actagagttggtg | AATAGTAC | tevopreQ1 |
| rosa peg spacer1b attB35 | agtcgcttctcgattatggg | cr772 | ggcgtgcccttgggctccccgggcgcgt | ataatcgagaagc | AAATAACG | tevopreQ1 |
| rosa peg spacer2b attB35 | ccctgggcgttgccctgcag | cr772 | ggcgtgcccttgggctccccgggcgcgt | cagggcaacgccc | GCTTAACT | tevopreQ1 |
| rosa peg spacer3a attB35 | tctcaaatggtataaaactc | cr772 | ggggagcccaagggcacgccctggcac | ttttataccattt | TACACACG | tevopreQ1 |
| rosa peg spacer3b attB35 | ggagcttagtcattcacctg | cr772 | ggcgtgcccttgggctccccgggcgcgt | gtgaatgactaag | AGAAAGAT | tevopreQ1 |
| rosa peg spacer4a attB30 | atgcctggtagggatgcaaa | cr772 | gcgtgcccttgggctccccgggcgc | gcatccctaccag | AACTCAAG | tevopreQ1 |
| rosa peg spacer4b attB30 | caccaccaaagtgtagcatc | cr772 | cggggagcccaagggcacgccctgg | gctacactttggt | CATTATTA | tevopreQ1 |
| rosa peg spacer4a attB35 | atgcctggtagggatgcaaa | cr772 | ggggagcccaagggcacgccctggcac | gcatccctaccag | ATCTCAGA | tevopreQ1 |
| rosa peg spacer4b attB35 | caccaccaaagtgtagcatc | cr772 | ggcgtgcccttgggctccccgggcgcgt | gctacactttggt | TATTATAA | tevopreQ1 |
| rosa peg spacer4a attB40 | atgcctggtagggatgcaaa | cr772 | ggcgtgcccttgggctccccgggcgcgtac | gcatccctaccag | AATTAAAT | tevopreQ1 |
| rosa peg spacer4b attB40 | caccaccaaagtgtagcatc | cr772 | Ggggagcccaagggcacgccctggcacccg | gctacactttggt | TAACTTAC | tevopreQ1 |
| rosa peg spacer4a attB45 | atgcctggtagggatgcaaa | cr772 | ggcgtgcccttgggctccccgggcgcgtactcc | gcatccctaccag | AATCTATA | tevopreQ1 |
| rosa peg spacer4b attB45 | caccaccaaagtgtagcatc | cr772 | ggggagcccaagggcacgccctggcacccgca | gctacactttggt | CATATATG | tevopreQ1 |
| rosa peg spacer4a attB50 | atgcctggtagggatgcaaa | cr772 | ggcgtgcccttgggctccccgggcgcgtactccac | gcatccctaccag | AATTAATT | tevopreQ1 |
| rosa peg spacer4b attB50 | caccaccaaagtgtagcatc | cr772 | ggggagcccaagggcacgccctggcacccgcaccg | gctacactttggt | AAATTTAT | tevopreQ1 |
| rosa peg spacer1a attP41 | tcgacaccaactctagtccg | cr772 | tggggtaacctttgagttctctcagttgggggc | actagagttggtg | AAACCCTT | tevopreQ1 |
| rosa peg spacer1b attP41 | agtcgcttctcgattatggg | cr772 | actgagagaactcaaaggttaccccagttggggc | ataatcgagaagc | AAATATAA | tevopreQ1 |
| rosa peg spacer2b attP41 | ccctgggcgttgccctgcag | cr772 | actgagagaactcaaaggttaccccagttggggc | cagggcaacgccc | CTTTCAAA | tevopreQ1 |
| rosa peg spacer3a attP41 | tctcaaatggtataaaactc | cr772 | tggggtaacctttgagttctctcagttgggggc | ttttataccattt | TTCTACTA | tevopreQ1 |
| rosa peg spacer3b attP41 | ggagcttagtcattcacctg | cr772 | actgagagaactcaaaggttaccccagttggggc | gtgaatgactaag | AATAAGAA | tevopreQ1 |
| rosa peg spacer4a attP35 | atgcctggtagggatgcaaa | cr772 | ggtaacctttgagttctctcagttggg | gcatccctaccag | ATCTATTC | tevopreQ1 |

| | | | | | | |
|---|---|---|---|---|---|---|
| rosa peg spacer4b attP35 | caccaccaaagtgtagcatc | cr772 | gagagaactcaaaggttaccccagttgg | gctacactttggt | ATAAGACT | tevopreQ1 |
| rosa peg spacer4a attP41 | atgcctggtagggatgcaaa | cr772 | tggggtaacctttgagttctctcagttgggggc | gcatccctaccag | ATCTAATG | tevopreQ1 |
| rosa peg spacer4b attP41 | caccaccaaagtgtagcatc | cr772 | actgagagaactcaaaggttaccccagttggggc | gctacactttggt | TAATAACG | tevopreQ1 |
| rosa peg spacer4a attP45 | atgcctggtagggatgcaaa | cr772 | gggtaacctttgagttctctcagttgggggcgt | gcatccctaccag | ATTTAAAT | tevopreQ1 |
| rosa peg spacer4b attP45 | caccaccaaagtgtagcatc | cr772 | agagaactcaaaggttaccccagttggggcac | gctacactttggt | ATAAACAA | tevopreQ1 |
| rosa peg spacer4a attP50 | atgcctggtagggatgcaaa | cr772 | gggtaacctttgagttctctcagttgggggcgtag | gcatccctaccag | ATTAACTA | tevopreQ1 |
| rosa peg spacer4b attP50 | caccaccaaagtgtagcatc | cr772 | agagaactcaaaggttaccccagttggggcactac | gctacactttggt | ACATACAA | tevopreQ1 |
| rosa peg spacer4a Bxb1attP48 | atgcctggtagggatgcaaa | cr772 | taccgtacaccactgagaccgcggtggttgaccagacaaacct | gcatccctaccag | AAACCACA | tevopreQ1 |
| rosa peg spacer4b Bxb1attP48 | caccaccaaagtgtagcatc | cr772 | gtctggtcaaccaccgcggtctcagtggtgtacggtacaaacct | gctacactttggt | CTCATCTT | tevopreQ1 |
| rosa peg spacer4a Bxb1attB38 | atgcctggtagggatgcaaa | cr772 | acgacggcggtctccgtcgtcaggatcat | gcatccctaccag | AACTTCGT | tevopreQ1 |
| rosa peg spacer4b Bxb1attB38 | caccaccaaagtgtagcatc | cr772 | acgacggagaccgccgtcgtcgacaagcc | gctacactttggt | CCATATAA | tevopreQ1 |
| rosa peg spacer 4a PaO1 attB | atgcctggtagggatgcaaa | cr772 | cctacatgctcgaagggcgtatgcgcc | gcatccctaccag | ATCCTACA | tevopreQ1 |
| rosa peg spacer 4b PaO1 attB | caccaccaaagtgtagcatc | cr772 | acgcccttcgagcatgtaggtcacgg | gctacactttggt | CCATATAA | tevopreQ1 |

| | |
|---|---|
| cr772 Scaffold (12) | GTTTAAGAGCTAAGCTGGAAACAGCATAGCAAGTTTAAATAAGGCTAGTCCGTTATCAACTCGAAAGAGTGGCACCGAGTCGGTGC |
| TevopreQ1 (13) | CGCGGTTCTATCTAGTTACGCGTTAAACCAACTAGAA |

List of epegRNA plasmids. The optimized cr772 scaffold (12) was substituted for the epegRNA scaffold on pegLIT website (13) to generate linker sequences.
